# IFNγ-induced IRF1 synergizes with TLR7 signals to tune the IRF4-IRF8 axis and drive pathogenic effector B cell fate

**DOI:** 10.64898/2026.02.06.704376

**Authors:** Eddie-Williams Owiredu, Ashley J. Denslow, Suki Chen, Betty Mousseau, Jeremy B. Foote, Guang Yang, Jessica N. Peel, Rebecca Burnham, Kelsey Browning, Christopher D. Scharer, Troy D. Randall, Esther Zumaquero-Martinez, Frances E. Lund

## Abstract

Interferon regulatory factor 1 (IRF1), a transcription factor encoded within the 5q31 locus harboring systemic lupus erythematosus (SLE) associated variants, promotes inflammatory responses by T and myeloid cells. Although IFNγ-activated B cells also express IRF1, its role in B cell biology and SLE is unclear. Here, we use a mouse SLE model, single-cell multiomics, and human B cells to show that IRF1 intrinsically regulates *Irf4* gene chromatin accessibility and expression in B cells to control the balance between the antibody secreting cell (ASC) lineage commitment factor, IRF4, and the B cell identity factor, IRF8. We demonstrate that IRF1, through its integration of IFNγ and TLR7 induced transcriptional programs, tips B cells toward a terminal effector inflammatory AC fate at the expense of preserving more stem-like, resting and regulatory B cells that do not elicit autoantibody-associated pathology in SLE. Thus, IRF1 serves as a central node controlling B cell-driven autoimmune disease.

## INTRODUCTION

Systemic lupus erythematosus (SLE) is a prototypic systemic autoimmune disease characterized by loss of self-tolerance and multi-organ inflammation that can progress to life-threatening complications.^1^ A major driver of SLE pathology is the B cell lineage-derived autoantibodies (autoAbs) that form immune complexes, activate Fc receptor and complement pathways, and propagate inflammatory circuits.^2^ This places B lymphocytes at the core of SLE pathogenesis, both as immunoregulatory hubs that shape immune activation and as progenitors of autoAb-secreting cells.

Multiple B cell subsets can differentiate into the short- and long-lived antibody secreting cells (ASC) that produce autoAbs in SLE. Germinal center (GC) responses generate affinity-matured class-switched long-lived ASC that can sustain autoAb levels over years.^3^ GC-derived memory B cells (MBC), which are also long-lived, can serve as a persistent autoreactive reservoir as these MBC may rapidly differentiate into proliferating short-lived autoAb-producing ASC upon exposure to inflammatory signals and autoantigen.^4^ Conversely, the GC-independent extrafollicular pathway^5^ leads to the generation of DN2 (CD27^−^IgD^−^ CD11c^+^Tbet^+^CXCR5^−^) B cells^6^ or their related age-associated B cells in mice^7^ (ABC), which can enter the MBC compartment or rapidly generate short-lived ASC, particularly during flares.

A defining molecular feature of SLE is a type I and type II interferon (IFN) signature.^8,9^ Indeed, IFNγ levels are often elevated in SLE patients years before onset of clinical symptoms and diagnosis^10,11^ while IFNα levels correlate with disease activity and organ involvement.^12^ Both type I and type II IFNs activate JAK/STAT pathways and induce antiviral and inflammatory transcriptional modules^13^ and both cytokines can support B cell differentiation into ASC.^14–16^ Moreover, IFNγ signals license B cells to differentiate^15^ in response to TLR7 ligands, including nucleic acid-containing autoantigens. TLR7 engagement in B cells^17,18^ leads to MYD88-dependent induction of the IRAK and TRAF nodes and converges on activation of transcription factors (TF) in the NFκB, AP-1, and interferon regulatory factor (IRF) families. In turn, these TFs support proliferation and differentiation programs that can accelerate ASC formation through GC-dependent and extrafollicular pathways.^19,20^ Importantly, IFN and TLR7 pathways further reinforce one another as IFN priming enhances responsiveness to TLR ligands and TLR7 ligation amplify IFN-driven programs, creating a feed forward circuit that favors pathogenic B cell states.^9,15,19^ Indeed, TLR7 gain-of-function variants, elevated TLR7 expression, and exaggerated TLR7-IFN signaling represent major drivers of SLE.^21–23^

Integration of TLR and inflammatory sensing pathways is executed through TF networks, which include IRF TFs that link cytokine and innate-sensing inputs to lineage-specific differentiation and functional programs. Nine different IRF TFs have been described,^24^ with IRF1 considered the prototypic example of an IFN-inducible TF downstream of type I and type II IFNR signaling.^25^ Although IRF1 was initially characterized as an antiviral and tumor suppressor,^26,27^ IRF1 is now recognized as a broad regulator of antigen presentation, apoptosis, cytokine production, and immuno-metabolic programs across immune lineages.^24^ Consistent with its role as an amplifier of inflammatory responses, IRF1 and its downstream pro-inflammatory gene programs are activated in immune cells isolated from the blood and kidney of SLE patients.^28–30^ Moreover, human genetic analyses suggest that IRF1 functions as an autoimmune susceptibility node, as IRF1-associated SNPs have been identified in the 5q31 locus that is linked to SLE and other autoimmune diseases.^31–34^

Despite the links between IFNs, IRF1 and SLE, a potential role for IRF1 in B cell-driven autoimmunity remains poorly defined. However, it is well appreciated that other IRFs, including IRF5,^35–37^ IRF4^38–41^ and IRF8^40,42^ play important roles in B cell biology. Indeed, IRF4 and IRF8 form a core axis that governs B cell fate decisions with IRF4 driving ASC programming and survival and IRF8 restraining terminal ASC differentiation and maintaining B lineage identity.^43^ We previously showed that IRF1 is upregulated in IFNγ-activated B effector 1 (Be1) cells and that its expression correlates with the differentiation potential of these cells.^14^ Given the known role for IFNγ and its downstream regulator STAT1 in driving B cell differentiation and Ab production in the settings of antiviral immunity^14,44^ and autoimmunity,^45,46^ we hypothesized that IRF1 might couple IFNγ and TLR7 signals to the core ASC differentiation machinery. We tested this hypothesis using a TLR7-driven lupus-prone mouse model^21^ and B cells from healthy donors and SLE patients. We show *Irf1*/*IRF1* links IFNγ- and TLR7-driven B cell differentiation in both mouse and human B cells and that B cell selective deletion of *Irf1* shifts B cell fate away from pathogenic ABC and ASC programs toward resting, stem-like MBC fates, with concurrent reductions in autoAb production, decreased kidney pathology, and improved survival. Mechanistically, IRF1 tunes chromatin accessibility at the *Irf4* locus and controls the balance of *Irf4*/*IRF4* and *Irf8*/*IRF8* in human and mouse ASC precursors. Collectively, these data support a conserved role for IRF1 as an inflammatory sensing rheostat that is directly wired into the IRF4- and IRF8-governed B cell differentiation landscape in the setting of autoimmunity.

## RESULTS

### *Irf1* regulates the composition of the B cell compartment in Yaa.*Fcgr2b*^−/−^ mice

Although IRF1 activates type I and type II IFN gene networks^25^ that are known to be dysregulated in immune cells from SLE patients,^9,28–30^ it is not known whether IRF1 also drives autoAb responses in SLE. To address this question, we intercrossed C57BL/6J Yaa.*Fcgr2b*^−/−^ mice (YaaFc), which develop a TLR7-dependent spontaneous SLE-like disease,^21^ with *Irf1*^−/−^ mice^47^ to generate an autoimmune-prone mouse strain lacking *Irf1* in all cell types (YaaFc.*Irf1*^−/−^). We then performed flow cytometric analysis on spleen and mediastinal lymph node (medLN) cells from the mice at five months of age (Fig. 1A) – a time when lupus-like clinical manifestations are easily measured in YaaFc mice. Consistent with our previous analysis of C57BL/6J *Irf1*^−/−^ mice,^48^ global *Irf1* deletion resulted in a significant reduction of the splenic marginal zone B (MZB) cell compartment in the YaaFc.*Irf1*^−/−^ mice (Fig. S1A-B). By contrast, YaaFc.*Irf1*^−/−^ spleens (Fig. 1B-C) and medLN (Fig. S1C-D) harbored higher frequencies and numbers of GCB cells compared to YaaFc controls. Consistent with the increased size of the GC compartment, spleen (Fig. 1D-E) and medLN (Fig. S1E-F) IgG^+^ MBC were also increased in YaaFc.*Irf1^−/−^*mice. However, the frequency and number of CD11c^+^T-bet^+^CD21^neg^CD23^neg^ ABC^49–51^ were significantly decreased in spleen (Fig. 1F-G) and medLN (Fig. S1G-H) of YaaFc.*Irf1^−/−^*mice compared to YaaFc mice. Finally, total (Fig. 1H-I) and IgG^+^ (Fig. 1J-K) CD138^+^ ASC were reduced in the spleen and medLN (Fig. S1I-L) of YaaFc.*Irf1*^−/−^ mice. The loss of ASC was further confirmed by ELISPOT on bulk splenocytes, which revealed that IgG-secreting ASC (Fig. 1L-M) were significantly reduced in the YaaFc.*Irf1*^−/−^ spleens. Thus, IRF1 appears to remodel the composition of the splenic and LN B cell compartments in this SLE-prone mouse strain.

**Figure 1.**
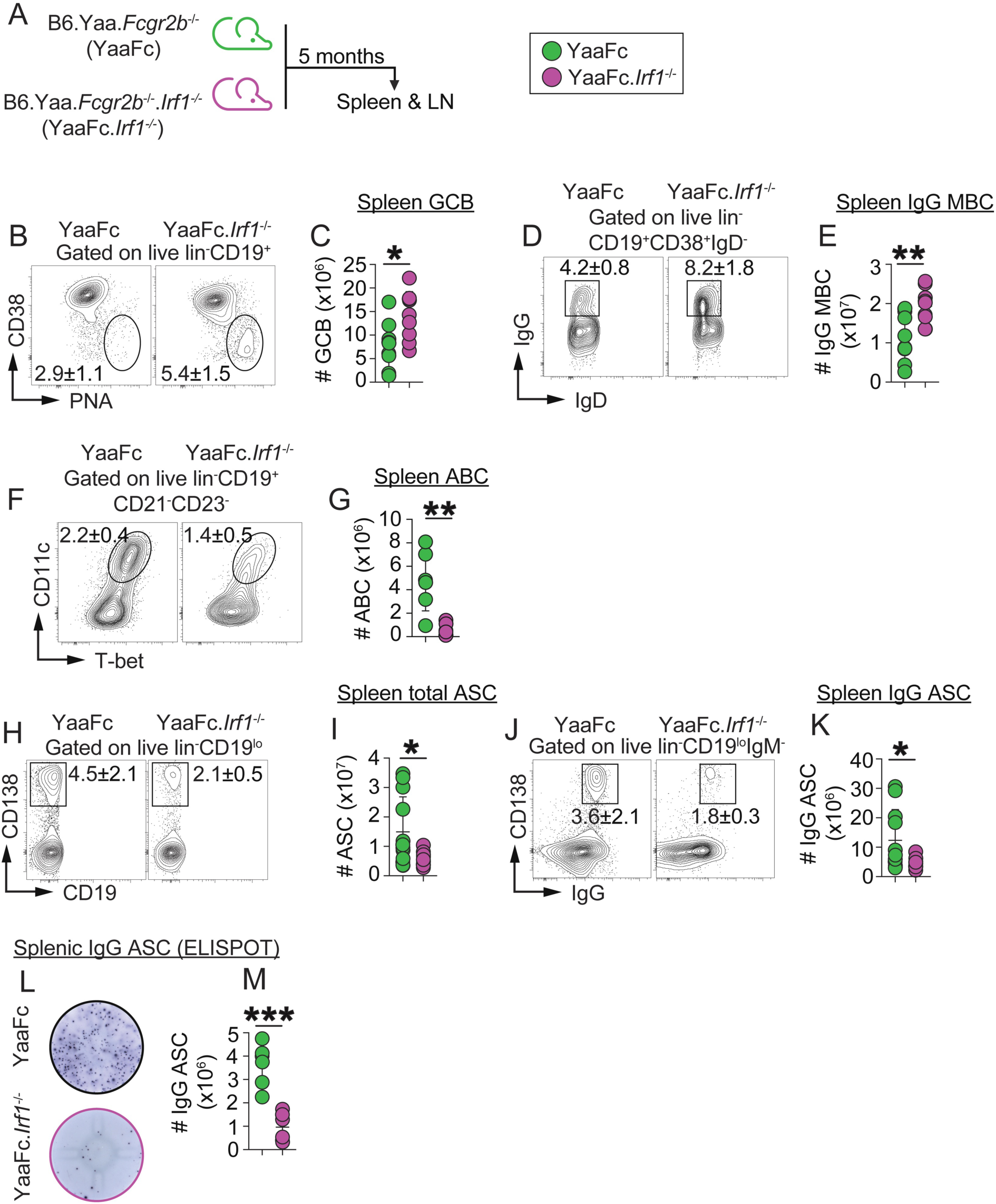
*Irf1* regulates homeostasis of the B cell compartment in lupus-prone Yaa.*Fcgr2b*^−/−^ mice. (**A**) Experimental design schematic. Spleens and medLN of Yaa.*Fcgr2b*^−/−^ (YaaFc) and Yaa.*Fcgr2b*^−/−^.*Irf1^−/−^* (YaaFc.*Irf1*^−/−^) lupus-prone mice were harvested and analyzed at 5 months. (**B-K**) Enumeration of B cell subsets in the spleens of YaaFc and YaaFc.*Irf1*^−/−^ mice by flow cytometry. Representative flow plots showing frequencies (mean±SD) of CD19^+^CD38^lo^PNA^+^ GCB cells (**B**), CD19^+^CD38^+^IgD^−^IgG^+^ MBC (**D**), CD19^+^CD21^−^CD23^−^CD11c^+^T-bet^+^ ABC (**F**), CD19^lo^CD138^+^ ASC (**H**) and CD19^lo^CD138^+^IgG^+^ ASC (**J**). Data reported as absolute numbers of splenic GCB cells (**C**), IgG^+^ MBC (**E**), ABC (**G**), ASC (**I**) and IgG^+^ ASC (**K**). (**L-M**) ELISPOT analysis of splenic YaaFc and YaaFc.*Irf1*^−/−^ IgG-producing ASC. Representative ELISPOTs shown (**L**) and reported as absolute numbers (**M**) of IgG^+^ ASC. Data shown from individual mice (**B-K, M**). Data representative of two independent experiments (n ≥ 5 mice/group). Significance determined by unpaired two-tailed Student’s t tests. *p < 0.05, **p < 0.01, ***p < 0.001. See **Fig. S1** for analysis of YaaFc and YaaFc.*Irf1*^−/−^ medLN.

### *Irf1* promotes autoreactive ASC responses and kidney pathology in lupus-prone mice

To determine whether *Irf1* deficiency also attenuated the humoral hallmarks of lupus, we next measured the frequencies of splenic autoAb-producing ASC in YaaFc and YaaFc.*Irf1*^−/−^ mice and observed a significant reduction in the number of splenic dsDNA-reactive IgG^+^ ASC in YaaFc.*Irf1*^−/−^ mice relative to YaaFc controls (Fig. 2A-B). To determine whether this defect translated to reduced systemic autoAb load, we measured serum antinuclear autoAb (ANA) by indirect immunofluorescence using HEp-2 cells. Although both cohorts produced detectable IgG^+^ ANA, sera from YaaFc.*Irf1*^−/−^ mice displayed weaker nuclear staining (Fig. 2C-D, Table S1) and serial dilution of the sera showed that titers of IgG^+^ ANA were significantly decreased in the YaaFc.*Irf1*^−/−^ mice (Fig. S1M-N, Table S1).

**Figure 2.**
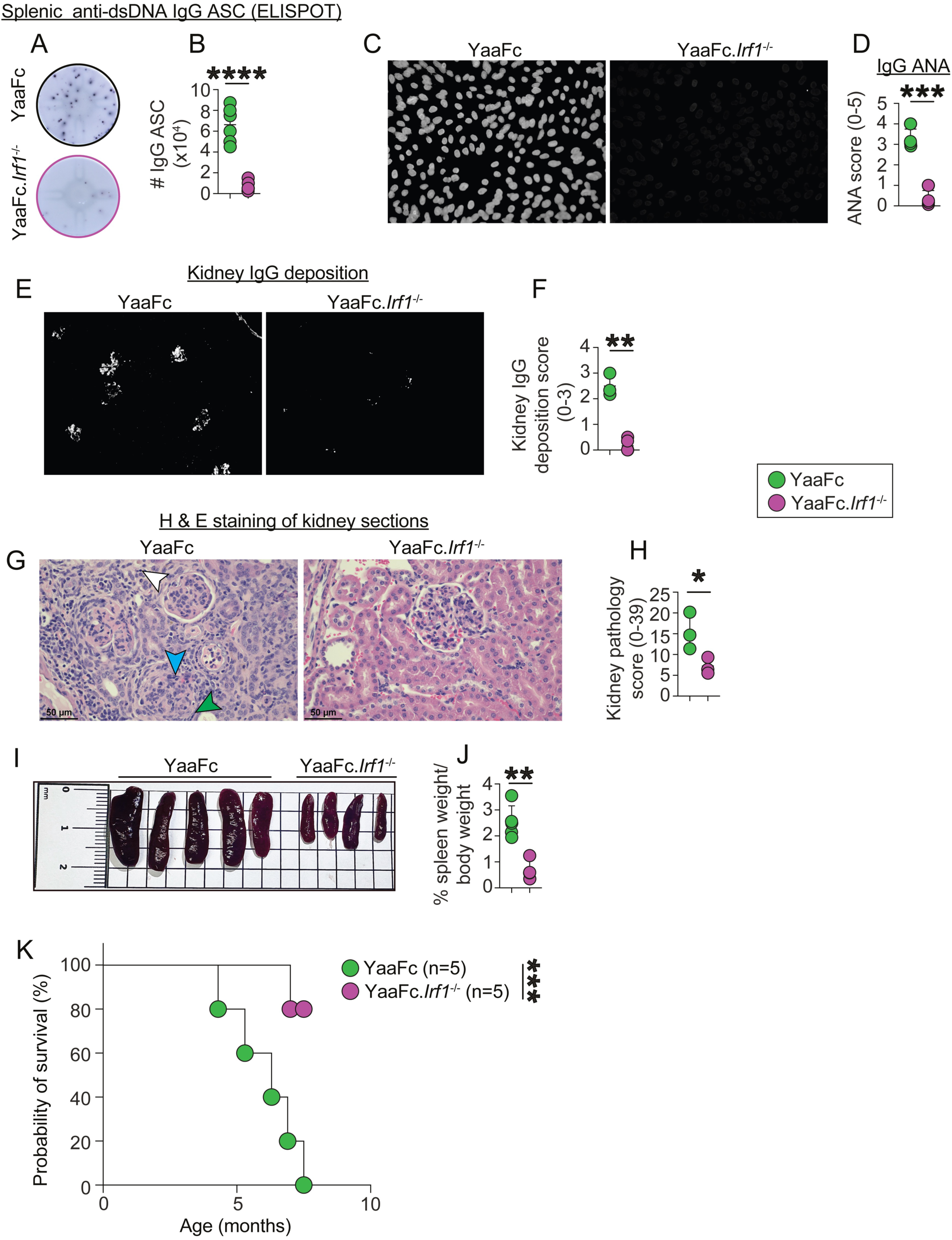
*Irf1* promotes autoreactive ASC responses and kidney pathology in lupus-prone mice. (**A-H**) Analysis of autoreactive ASC and Ab from 5-month YaaFc and YaaFc.*Irf1*^−/*-*^ mice. ELISPOT analysis of splenic anti-dsDNA specific IgG^+^ ASC. Representative ELISPOTs shown (**A**) and data reported as numbers (**B**) of IgG^+^ autoreactive splenic ASC. Binding of serum (1:400 dilution) IgG anti-nuclear antibodies (ANAs) to HEp-2 cells with representative ANA images shown (**C**) and data reported as ANA scores (**D**). Immunofluorescence analysis of glomerular IgG deposition in kidney sections with representative immunofluorescence images shown (**E**) and data reported as IgG deposition scores (**F**). (**G-H**) Histopathology analysis of kidney tissue from 5-month YaaFc and YaaFc.*Irf1*^−/−^ mice. Sections stained with H&E. Representative images (**G**) shown and data reported as kidney pathology scores (**H**). Glomerular hypercellularity (blue arrowhead); Crescent formation (green arrowhead); Interstitial inflammation (white arrowhead). (**I-J**) Size and weight of spleens from 5-month YaaFc and YaaFc.*Irf1*^−/−^ mice. Representative spleens shown (**I**) and data reported as the percent spleen weight to body weight (**J**) for each animal. (**K**) Survival analysis of YaaFc and YaaFc.*Irf1*^−/−^ mice with data reported until 100% of the YaaFc mice succumbed to disease. Data shown from individual mice (**B, D, F, H, J**). Data representative of two independent experiments (n ≥ 5 mice/group (**A-B, I-K**), 4 mice/group (**C-D**) or 3 mice/group (**E-H**). Significance determined using unpaired two-tailed Student’s t test (**B, D, F, H, J**) or log-rank (Mantel-Cox) test (**K**). See IgG ANA, renal IgG deposition, and histopathology scores in **Table S1**. *p < 0.05, **p < 0.01, ***p < 0.001, ****p < 0.0001.

Given these results, we next examined immune complex deposition in the kidneys, as Ab deposition is a hallmark of end-organ pathology in SLE.^1^ Immunofluorescence staining for IgG Ab revealed easily detected immune complex deposition in YaaFc kidney sections, while kidney sections from YaaFc.*Irf1*^−/−^ mice showed markedly attenuated Ab deposition (Fig. 2E-F, Table S1). To determine whether reduced immune complex deposition corresponded with less tissue damage, we stained kidney sections from both strains of mice with hematoxylin and eosin (H&E) and quantitated changes in glomerular hypercellularity, crescent formation, and interstitial inflammation. While these changes were evident in YaaFc kidneys (Fig. 2G-H, Table S1), these features were significantly diminished in YaaFc.*Irf1*^−/−^ mice, resulting in a lower total kidney pathology score (Fig. 2H). Finally, other systemic indicators of inflammation were also improved in the *Irf1* deficient mice. For example, splenomegaly, a cardinal feature of lupus that reflects extramedullary hematopoietic cell expansion,^52^ was significantly reduced in YaaFc.*Irf1*^−/−^ mice (Fig. 2I-J). Moreover, the reduction in inflammation was associated with prolonged survival of the YaaFc.*Irf1*^−/−^ mice compared to the autoimmune YaaFc animals (Fig. 2K). Taken together, these data indicate that IRF1 promotes systemic autoimmunity in a SLE mouse model and does so, at least in part, by driving autoreactive ASC responses and exacerbating immune complex-driven tissue pathology.

### B cell-intrinsic *Irf1* controls expansion of pathogenic B cell subsets and autoreactive ASC in lupus-prone mice

Given the attenuated autoreactive ASC and Ab response in the YaaFc.*Irf1*^−/−^ mice and the fact that IFNγ signaling in B cells can drive the autoimmune humoral response in mice,^45,46^ we predicted that IRF1, an IFNγ-inducible TF in B cells^14,48^ would intrinsically regulate B cell autoimmune responses. To test this, we generated 80:20 bone marrow (BM) chimeras by reconstituting lethally irradiated B cell deficient YaaFc mice (YaaFc.μMT mice^15^) with either 80% YaaFc.μMT BM + 20% YaaFc BM (B-YaaFc mice) or with 80% YaaFc.μMT BM + 20% YaaFc.*Irf1*^−/−^ BM (B-YaaFc.*Irf1*^−/−^ mice). The B-YaaFc mice were *Irf1* sufficient in all cell types while the B-YaaFc.*Irf1*^−/−^ mice lacked *Irf1* in 100% of B lineage cells but were competent to express *Irf1* in 80% of all other BM-derived cells and 100% of all radiation resistant cells (Fig. 3A). While irradiation and BM reconstitution slowed the development of autoimmunity, by 10 months post-reconstitution, the B-YaaFc mice exhibited most hallmarks of disease, with easily detected splenic ABC (Fig. 3B-C) and ASC (Fig. 3D-G). Strikingly, much like the animals with a global deletion of *Irf1*, mice with *Irf1* selectively deleted in all B cells exhibited a near absence of splenic CD11c^+^Tbet^+^ ABC (Fig. 3B-C), CD138^+^ASC (Fig. 3D-E), IgG^+^CD138^+^ ASC (Fig. 3F-G) and CD138^+^IRF4^+^ ASC (Fig. S2A-C) as measured by flow cytometry. Similar results were observed in the medLN (Fig. S2D-L). These results were confirmed by ELISPOT, which revealed a near complete loss of IgG^+^ ASC (Fig. 3H-I) and anti-dsDNA IgG^+^ ASC (Fig. 3J) in the spleens of B-YaaFc.*Irf1*^−/−^ mice. By contrast and identical to what we observed in the mice with a global *Irf1* deficiency, the GCB (Fig. 3K-L) and MBC (Fig. 3M-N) compartments were increased in the spleen (Fig. 3K-N) and medLN (Fig. S2M-P) of the B-YaaFc.*Irf1*^−/−^ mice. Finally, we found that spleen size was reduced in B-YaaFc.*Irf1*^−/−^ mice (Fig. 3O-P), and that these animals exhibited attenuated kidney pathology (Fig. 3Q, Table S1) and improved overall survival (Fig. 3R) compared to B-YaaFc mice. Thus, B cell intrinsic IRF1 expression is both necessary and sufficient to induce expansion of the pathogenic ABC and autoreactive ASC subsets and promote progression of systemic autoimmunity in the YaaFc model.

**Figure 3.**
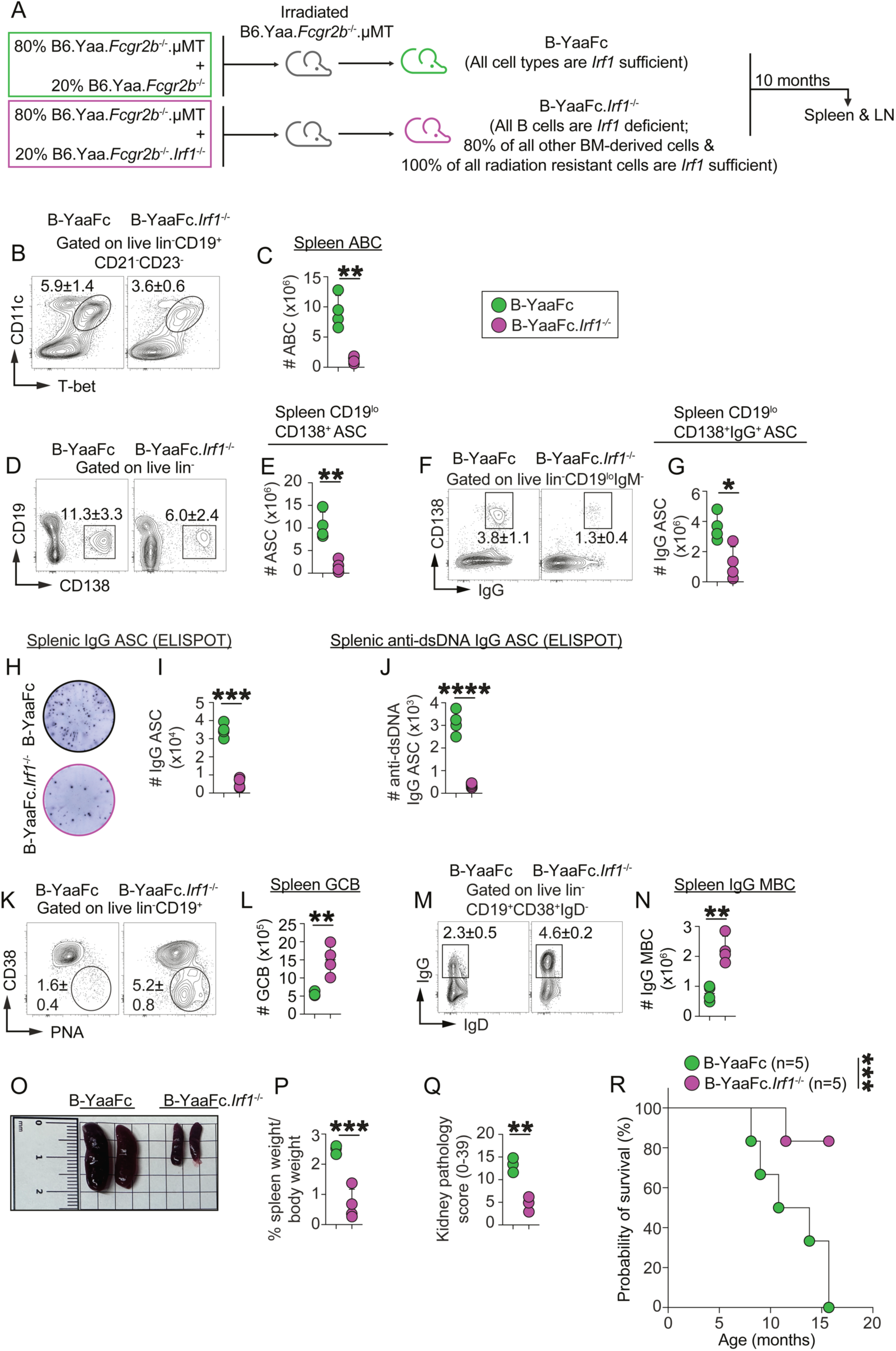
B cell-intrinsic *Irf1* is required for the expansion of pathogenic B cell subsets and autoreactive ASC in lupus-prone mice. (**A**) Experimental design schematic. The 80:20 bone marrow (BM) chimeras were generated by reconstituting lethally irradiated B cell deficient YaaFc mice (YaaFc.μMT mice^15^) with either 80% YaaFc.μMT BM + 20% YaaFc BM (B-YaaFc mice) or with 80% YaaFc.μMT BM + 20% YaaFc.*Irf1*^−/−^ BM (B-YaaFc.*Irf1*^−/−^ mice). B-YaaFc and B-YaaFc.*Irf1*^−/−^ mice were analyzed at 10 months post-reconstitution (**B-Q**) or were entered into survival studies (**R**). (**B-N**) Flow cytometry (**B-G, K-N**) and ELISPOT (**H-J**) analysis of splenic B cell subsets and ASC in B-YaaFc and B-YaaFc.*Irf1*^−/−^ mice. Representative flow plots showing frequencies of ABC (**B**), CD138^+^ ASC (**D**), IgG-expressing CD138^+^ ASC (**F**), GCB cells (**K**), and IgG^+^ MBC (**M**). Data reported as absolute numbers of ABC (**C**), CD18^+^ ASC (**E**), IgG expressing CD138^+^ ASC (**G**), GCB cells (**L**) and IgG^+^ MBC (**N**). ELISPOT analysis of splenic splenic YaaFc and YaaFc.*Irf1*^−/−^ IgG^+^ ASC (**H-I**) and ds-DNA specific IgG^+^ ASC (**J**). Representative ELISPOTs (**H**) shown with absolute numbers of ASC (**I-J**) reported. (**O–P**) Size and weight of B-YaaFc and B-YaaFc.*Irf1*^−/−^ spleens. Representative spleens shown (**O**) with data reported as percentage of spleen weight to body weight (**P**) for each animal. (**Q**) Histopathology analysis of H&E stained kidney sections. Data reported as kidney pathology scores. (**R**) Survival analysis of B-YaaFc and B-YaaFc.*Irf1^−/−^* mice with data reported until 100% of B-YaaFc mice succumbed to disease. Data shown from individual mice (**B-G, I-N, P-R**). Data representative of two independent experiments (n ≥ 4 mice/group). Significance determined using unpaired two-tailed Student’s t tests (**B-G, I-N, P-Q**) or log-rank (Mantel–Cox) test (**R**). See **Figure S2** for medLN analysis and **Table S1** for histopathology scores. *p < 0.05, **p < 0.01, ***p < 0.001, ****p < 0.0001.

### *Irf1* shapes the transcriptional and epigenomic landscape of ASC subsets in lupus-prone mice

Given the significant and direct effect of IRF1 on B cell autoimmune responses, we predicted that IRF1 would regulate the transcriptional and/or epigenomic landscape of B cells in the setting of autoimmunity. To test this, we performed single nuclei multiomics analysis^53,54^ of purified IgD^neg^ antigen-experienced B cells from pooled spleens isolated from B-YaaFc and B-YaaFc.*Irf1*^−/−^ mice at 6 month post-reconstitution. We recovered 12,421 high-quality nuclei with paired RNAseq and ATACseq data that passed all quality control (QC) steps (B-YaaFc: 6,979 cells; B-YaaFc.*Irf1*^−/−^: 5,442 cells) (Fig. S3A). Unsupervised clustering of these cells identified 15 B cell clusters (B-C0 to B-C14) based on joint gene expression and chromatin accessibility (Fig. S3B, Table S2). These clusters were homogeneous (Fig. S3C) as measured by ROGUE scores^55^ and contained 182-2640 B cells per cluster with 68-3939 differentially expressed genes (DEGs) per cluster (Table S2). We then annotated the clusters based on expression of canonical marker genes associated with different B cell subsets (see Fig. S3D-E and Table S2 for genes used in subset annotation) and identified (Fig. 4A) residual naïve B cells (B-C12), MZB (B-C5), GCB (B-C4 light zone GCB, B-C6 dark zone GCB), MBC (B-C3, B-C13, B-C14), non-proliferating ASC (B-C0, B-C2, B-C8, B-C10, B-C11), proliferating ASC (B-C7), ABC (B-C1), and recent GC emigrants (B-C9). Consistent with our flow cytometry data, B-YaaFc.*Irf1*^−/−^ spleens harbored fewer transcriptionally defined MZB, ASC and ABC but appeared to have more GCB cells and recent GC emigrant B cells (Fig. 4B-C).

**Figure 4.**
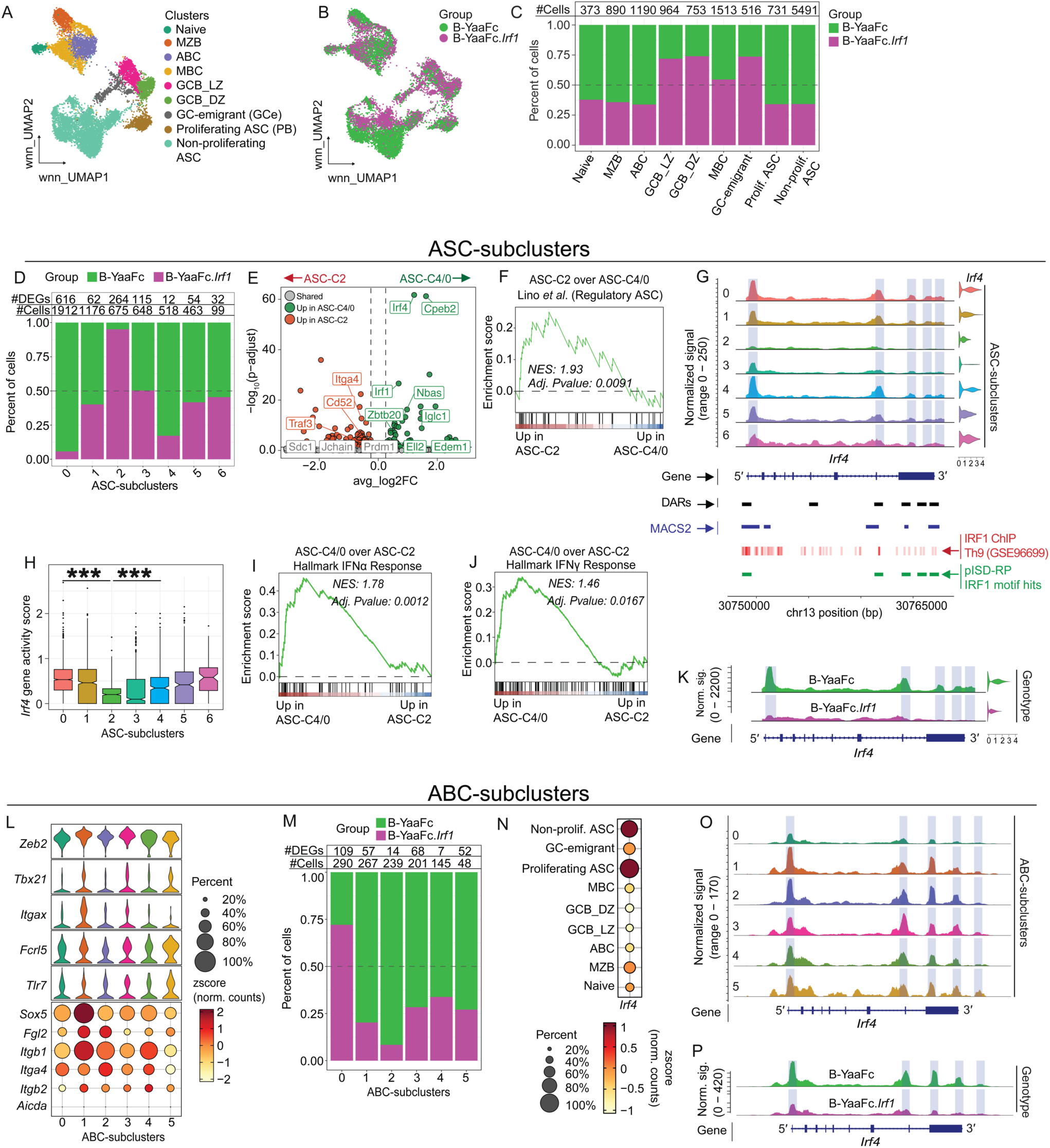
IRF1 controls *Irf4* expression and ASC effector programming in ASC and ABC from lupus prone mice. Single cell multiomics analysis (scRNA-seq, scATAC-seq) was performed on nuclei isolated from purified IgD^neg^ B cells from spleens of B-YaaFc and B-YaaFc.*Irf1*^−/−^ mice at 6-month post-reconstitution. (**A-C**) High-quality nuclei (n=6979 B-YaaFc and 5442 B-YaaFc.*Irf1*^−/−^, **Fig. S3A**) passing QC analysis underwent integrated WNN analysis and were visualized by UMAP (**Fig. S3B-C**). Individual UMAP clusters (B-C0 to B-C14, **Fig. S3B, right**) were assigned to specific B cell subsets (**A**) based on expression of canonical subsetting markers (see **Fig. S3D-E** for gene signatures used to identify B cell subsets). B cells of each genotype (B-YaaFc and B-YaaFc.*Irf1*^−/−^) were identified and displayed in a UMAP plot (**B**) with the data reported in (**C**) as the number of cells present in each subset and the relative abundance of the B-YaaFc and B-YaaFc.*Irf1*^−/−^ cells within the different B cell subsets. (**D-K**) Analysis of ASC subclusters. Unsupervised reclustering was performed using cells assigned as non-proliferating ASC (See **A** and **Fig. S3D-E**). Seven ASC subclusters (ASC-C0 to ASC-C6) were identified and the genotype of the cells in each ASC subcluster was determined (**Fig. S3F**). Data reported in (**D**) as the number of cells and DEG assigned to each ASC subcluster and the relative abundance of the B-YaaFc and B-YaaFc.*Irf1*^−/−^ B cells within the different ASC subclusters. Volcano plot (**E**) comparing gene expression between ASC-C0/C4 and ASC-C2. Shown are DEGs upregulated in ASC-C0/C4 (green), upregulated in ASC-C2 (red) or genes equivalently expressed between ASC-C0/C4 and ASC- C2 (grey). GSEA (**F**) using a published regulatory ASC geneset^63^ to query ranked DEG list of ASC-C2 over ASC-C0/C4. Chromatin accessibility (**G**) surrounding the *Irf4* gene in the ASC-subclusters (**G**, top) showing DAR (G, black bars), MACS2-called peaks (**G**, blue bars), Publicly available (GSE96699) IRF1 Chip-seq signals surrounding the *Irf4* gene in IFNγ-stimulated Th9 cells (**G**, red bars) and predicted IRF1 binding sites within the *Irf4* locus based on multimodal integrated regulatory potential modeling^70^ following probabilistic *in silico* deletion of TF binding regions (pISD-RP)^71^ (**G**, green bars). *Irf4* gene activity score (**H**) for each ASC-subcluster. GSEA (**I-J**) using Hallmark gene lists for Type I (**I**) and Type II (**J**) IFN-response programs to query ranked DEG list of ASC-C0/C4 over ASC-C2. Chromatin accessibility (**K**) surrounding the *Irf4* gene among all non-proliferating ASC, divided by genotype. See **Fig. S3F-I** for additional analysis of ASC subclusters. (**L-O**) Analysis of ABC subclusters. Unsupervised reclustering was performed using cells assigned as ABC (See **A** and **Fig. S3D-E**). Six ABC subclusters (ABC-C0 to ABC-C5) were identified and the genotype of the cells in each ABC subcluster was determined (**Fig. S3J**). Gene expression (**L**) of select ABC-related genes among the ABC subclusters. Relative abundance (**M**) of the B-YaaFc and B-YaaFc.*Irf1*^−/−^ B cells within the different ABC subclusters. *Irf4* gene expression (**N**) by all major B cell subsets. Chromatin accessibility surrounding the *Irf4* gene in the ABC-subclusters (**O**) and in all ABC divided by genotype (**P**). See **Fig. S3K-O** for additional analysis of ABC subclusters. See **Table S2** for DEGs of ASC subclusters, DEGs of ASC-C0/4 over ASC-C2 subclusters, DEGs of ABC-subclusters, average expression of select genes among ABC-subclusters and motif enrichment analysis of ABC-C1 over ABC-C0. DEGs (**D**, **M**) were identified using auROC and Wilcoxon p-value based on Gaussian approximation (presto). Genes with |avg_log2FC| ≥0.1, FDR <0.05 and expressed in ≥10% of cells per cluster or group were considered significant. ***p < 0.001. Adjusted p value and NES scores for each GSEA provided (**F, I-J**, also see **Table S3**).

Next, we focused specifically on the non-proliferating terminally-differentiated ASC clusters (Fig. S3D-E). Despite our ELISPOT data showing that B-YaaFc.*Irf1*^−/−^ spleens contained few functional ASC (Fig. 3H-I), it was evident that some transcriptionally-identified ASC were present in B-YaaFc.*Irf1*^−/−^ spleens (Fig. 4C). Given that the ASC compartment is known to contain transcriptionally and functionally heterogeneous subpopulations,^56–58^ we hypothesized that the B-YaaFc.*Irf1*^−/−^ ASC might not exhibit transcriptional programming consistent with high secretory effector potential. To test this, we combined the cells in the transcriptionally defined non-proliferating ASC clusters (Fig. 4C, Fig. S3D-E) and performed unsupervised reclustering of the ASC to reveal seven distinct ASC subpopulations (ASC-C0 to ASC-C6) based on joint gene expression and chromatin accessibility (Fig. S3F). These subclusters contained 99-1,912 cells and 12-616 DEGs per cluster (Fig. 4D, Table S2). As expected, most ASC subclusters were enriched in B-YaaFc cells, with the ASC-C0 and ASC-C4 subclusters (2,430 cells) populated almost entirely by B-YaaFc cells (Fig. 4D). Since ASC-C0 and ASC-C4 exhibited few transcriptional differences (Table S2) we combined these two clusters for downstream analyses. By contrast, one cluster, ASC-C2 (675 cells), was almost exclusively populated with B-YaaFc.*Irf1*^−/−^ cells (Fig. 4D). ASC-C2 cells were unusual as these cells expressed canonical ASC markers (e.g. *Prdm1*, *Jchain*, *Sdc1* and *Xbp1*) and molecules that support Ab secretion (e.g. *Creb3l2*, *Hspa5*, *Pdia4*), but also expressed higher levels of regulators that oppose robust ASC function/survival, including known immunosuppressive genes (e.g. *Ctla4*, *Traf3*, *Itga4, Pten*, *Cd52*, *H2-Eb1*, *H2-Ab1, Rhoh*)^59–61^ (Fig. 4E, Table S2). Based on the ASC-C2 gene expression signature, we hypothesized that ASC-C2 might represent a regulatory ASC-like state that is reported to emerge in response to innate signals.^62–64^ To test this, we performed gene set enrichment analysis (GSEA) comparing ASC-C2 to the ASC-C0 and ASC-C4 cells using a published transcriptome dataset^63^ from regulatory plasma cells. Consistent with our prediction, ASC-C2 showed significant enrichment for genes that are upregulated in IL-10 producing regulatory plasma cells (Fig. 4F, Table S3) – cells that exhibited immunosuppressive activity and attenuated inflammatory responses.^62–64^ Consistent with this, chromatin accessibility surrounding the *Il10* gene was higher in ASC-C2 and this cluster also had more *Il10* transcripts and a significantly higher *Il10* gene activity score (Fig. S3G-H), which reflects increased chromatin accessibility across the *Il10* promoter and gene body in this cluster. It is known that the IL-10 producing regulatory plasma cells also express higher levels of *Ctla4*^63^ and, consistent with this, *Ctla4* expression was limited to cells in the ASC-C2 cluster (Table S2). Thus, the ASC-C2 cluster, which was enriched with B-YaaFc.*Irf1*^−/−^ ASC, included some ASC with regulatory-like features.

Unlike the ASC-C2 cluster that appeared composed of regulatory-like ASC, the ASC-C0 and ASC-C4 clusters (2,430 cells), which were dominated by B-YaaFc cells, displayed a typical high-secretory ASC transcriptional program. These ASC had upregulated a transcription-elongation factor (*Ell2*) (Fig. 4E, Table S2), which drives expression of the secretory IgH mRNA isoform,^65^ and had higher expression of genes in the unfolded protein response (UPR) and the endoplasmic reticulum (ER) secretory pathway^66,67^ (e.g. *Edem1*, *Nbas, Psmb8*, *Creld2*, *Txn1*, *Ubc*) (Fig. 4E, Table S2), which regulate protein-folding/trafficking and sustained Ab synthesis. The combined ASC-C0/C4 cluster was also marked by high expression of RNA-processing and translation factors (*Cpeb2*, *Mbnl1*, *Tnrc6b*, *Eif4e3*, *Rpl39*) that stabilize the hyper-transcriptional state of ASC, *Zbtb20* (Fig. 4E, Table S2), which reinforces ASC identity and persistence,^68^ and cytoskeletal/trafficking gene modules (*Tns3*, *Arhgap6*, *Rabgap1l*, *Rhobtb1*) as well as metabolic/stress adapters (*Eaf2*, *Lpin2*, *Adk*, *Trp53inp1*) (Table S2), which support a mature, secretory effector state.^69^ Not surprisingly, cells in ASC-C0 and ASC-C4 expressed high levels of *Irf4* (Fig. 4E, G), which is critical for establishment and maintenance^38–41^ of the ASC lineage. Consistent with the high expression of *Irf4* transcripts in ASC-C0 and ASC-C4 cells, these cells exhibited significantly increased chromatin accessibility in the promoter and gene body of *Irf4* (Fig. 4G). In striking contrast, the *Irf4* gene accessibility peaks were significantly decreased in cells from the ASC-C2 cluster (Fig. 4G) and ASC-C2 ASC had a significantly lower *Irf4* gene activity score compared to cells in ASC-C0 and ASC-C4 (Fig. 4H). Moreover, multimodal integrated regulatory potential modeling based on more than 10,000 ChIP-seq datasets from the Cistrome Data Browser (CistromeDB v3.0)^70^ following probabilistic *in silico* deletion of TF binding regions^71^ revealed that 5 out of the 6 significant differential accessible regions (DARs) surrounding the *Irf4* gene, including the promoter regions, contained binding sites for IRF1 (Fig. 4G). These IRF1 binding site predictions were in good concordance with publicly (GSE96699) available IRF1 ChIP data (Fig. 4G). Finally, GSEA comparing ASC-C0 and ASC-C4 cells to ASC-C2 cells revealed enrichment of hallmark type I and II IFN-response signatures in ASC-C0 and ASC-C4 cells (Fig. 4I-J, Table S3), consistent with the IFN-licensed effector programs that can drive pathogenic extrafollicular ASC trajectories implicated in lupus-like autoimmunity and human SLE.

Although ASC-C0, ASC-C4 and ASC-C2 were preferentially enriched by B-YaaFc or B-YaaFc.*Irf1*^−/−^ cells, the other ASC subclusters were composed of roughly equivalent numbers of ASC from each genotype (Fig. 4D, Fig. S3F). Since the clustering was unsupervised, this suggested that cells from both genotypes converged on overlapping transcriptional states. However, if *Irf1* was important for regulating chromatin accessibility at the *Irf4* locus, we hypothesized that all the ASC derived from B-YaaFc.*Irf1*^−/−^ mice should exhibit reduced accessibility across the *Irf4* regulatory elements. Consistent with this prediction, genotype-resolved DARs spanning the *Irf4* gene body and flanking regions revealed uniformly diminished *Irf4* accessibility in B-YaaFc.*Irf1*^−/−^ cells (Fig. 4K), regardless of ASC subcluster assignment (Fig. S3I). Peak-gene linking further connected multiple intragenic IRF1 motif-containing DARs across the *Irf4* locus to the *Irf4* promoter, consistent with coordinated cis regulatory coupling within this region (Fig. S3I). Thus, the ASC compartment in lupus-prone Yaa.Fc mice was composed of multiple transcriptionally distinct subpopulations that exhibited divergent regulatory, inflammatory and effector profiles. This heterogeneous ASC transcriptional landscape was shaped, at least in part, by B cell intrinsic expression of the TF IRF1, which appears to maintain *Irf4* locus accessibility and supports development of pathogenic, highly functional IFN-driven ASC subsets.

### *Irf1* regulates the effector potential of ABC from lupus-prone mice

Our data showed that IRF1 expression in B lineage cells regulates *Irf4* expression and the development of secretory inflammatory ASC. Given that IRF1 is upregulated in activated B cells responding to IFNγ,^14,48^ we predicted that IRF1-dependent programming of ASC development and function likely occurred before commitment to the ASC lineage within one or more of the ASC precursor populations. We first examined the IFNγ- and TLR7-dependent^50,51,72^ extrafollicular ABC population that rapidly differentiates into ASC and has been implicated in SLE immunopathology. Since emerging data suggests that the ABC compartment in mice is heterogenous,^51^ we reclustered the cells defined as ABC-like (B-C1 cluster, 1190 cells, Fig. S3D-E). We identified six distinct subclusters (Fig. S3J, ABC-C0 to ABC-C5, Table S2) and as expected, cells in all 6 ABC-like subclusters expressed *Zeb2* and *Tlr7* (Fig. 4L, Table S2), which are required for ABC development.^20,72,73^ Consistent with extrafollicular B cells, none of the clusters expressed *Aicda* (Fig. 4L, Table S2). Although ABC-C0, which contained 290 cells, was enriched for B-YaaFc.*Irf1*^−/−^ cells, all the remaining ABC subclusters were highly enriched in B-YaaFc cells (Fig. 4M). ABC-C1, the largest (267 cells) of the B-YaaFc cell dominated clusters, also expressed the highest levels of genes that support ABC development and function,^6,7^ including *Tbx21*, *Itgax*, *Itgb1*/*2* and *Sox5* (Fig. 4L). Expression of these genes was consistently lower in the B-YaaFc.*Irf1*^−/−^ dominated ABC-C0 cluster (Fig. 4L). Consistent with the transcriptional profile of the ABC-C1 cells, TF motif analysis revealed that ABC-C1 cells were enriched relative to ABC-C0 cells for chromatin accessible binding motifs for T-box TFs, like T-bet and EOMES (Fig. S3K, Table S2). Moreover, GSEA comparing the transcriptomes of ABC-C1 and ABC-C0 cells revealed that ABC-C1 cells were enriched for expression of the Hallmark type I and II IFN gene sets (Fig. S3L-M, Table S3). Additional GSEA comparing the transcriptomes of ABC-C1 and ABC-C0 cells to a published ABC transcriptome dataset^37^ further indicated that ABC-C1 cells were significantly and positively enriched for the “ABC up-regulated” gene signature and negatively enriched for the “ABC down-regulated” gene set (Fig. S3N-O, Table S3). Finally, while *Irf4* mRNA expression was generally low in the ABC compartment (Fig. 4N), chromatin accessibility within the *Irf4* gene was increased in ABC-C1 relative to ABC-C0 cells (Fig. 4O). Indeed, splitting the chromatin accessibility profile by genotype revealed that ABC-like cells from B-YaaFc.*Irf1*^−/−^ mice generally had lower chromatin accessibility surrounding the *Irf4* gene compared to cells from B-YaaFc mice (Fig. 4P). Taken together, these data suggest that the remaining ABC-like B-YaaFc.*Irf1*^−/−^ cells, while expressing the core ABC-defining TFs, lack the pathogenic, pre-ASC signature that is evident in the B-YaaFc cells.

### *Irf1* regulates the effector potential of GCB and post-GCB compartments in lupus-prone mice

Given the aberrant transcriptional and epigenetic profile of the extrafollicular ABC compartment in B-YaaFc.*Irf1*^−/−^ mice, we next asked whether IRF1 affected the follicular GCB cells in a similar manner. We therefore reclustered the B cells identified as dark and light zone GCB cells^74^ (B-C4 and B-C6, Fig. S3D-E). Not surprisingly, given that the GCB cell compartment was dominated by B-YaaFc.*Irf1*^−/−^ cells (see Fig. 4A-C), all three GCB cell subclusters (GCB-C0 to GCB-C2) were enriched for B-YaaFc.*Irf1*^−/−^ cells (Fig. S4A-B, Table S4). Therefore, we compared the transcriptomes of all GCB cells divided on the basis of genotype (B-YaaFc GC versus B-YaaFc.*Irf1*^−/−^ GC) and looked for potential genotype-specific shifts in GCB cell fate-associated programs. We performed GSEA using a published GC output reference dataset^75^ and compared the transcriptomes of B-YaaFc and B-YaaFc.*Irf1*^−/−^ GCB cells. We found that B-YaaFc GCB cells were enriched for gene signatures associated with ASC differentiation relative to MBC programming (Fig. 5A, Table S3). Consistent with this, B-YaaFc GCB cells upregulated expression of the ASC-associated chaperone *Mzb1* and increased expression of ER and UPR-associated genes^67^ like *Hspa5*, *Hsp90b1*, *Pdia4*, and *Edem1* (Table S4). Thus, the B-YaaFc GCB cell compartment appeared somewhat biased toward an ASC fate.

**Figure 5.**
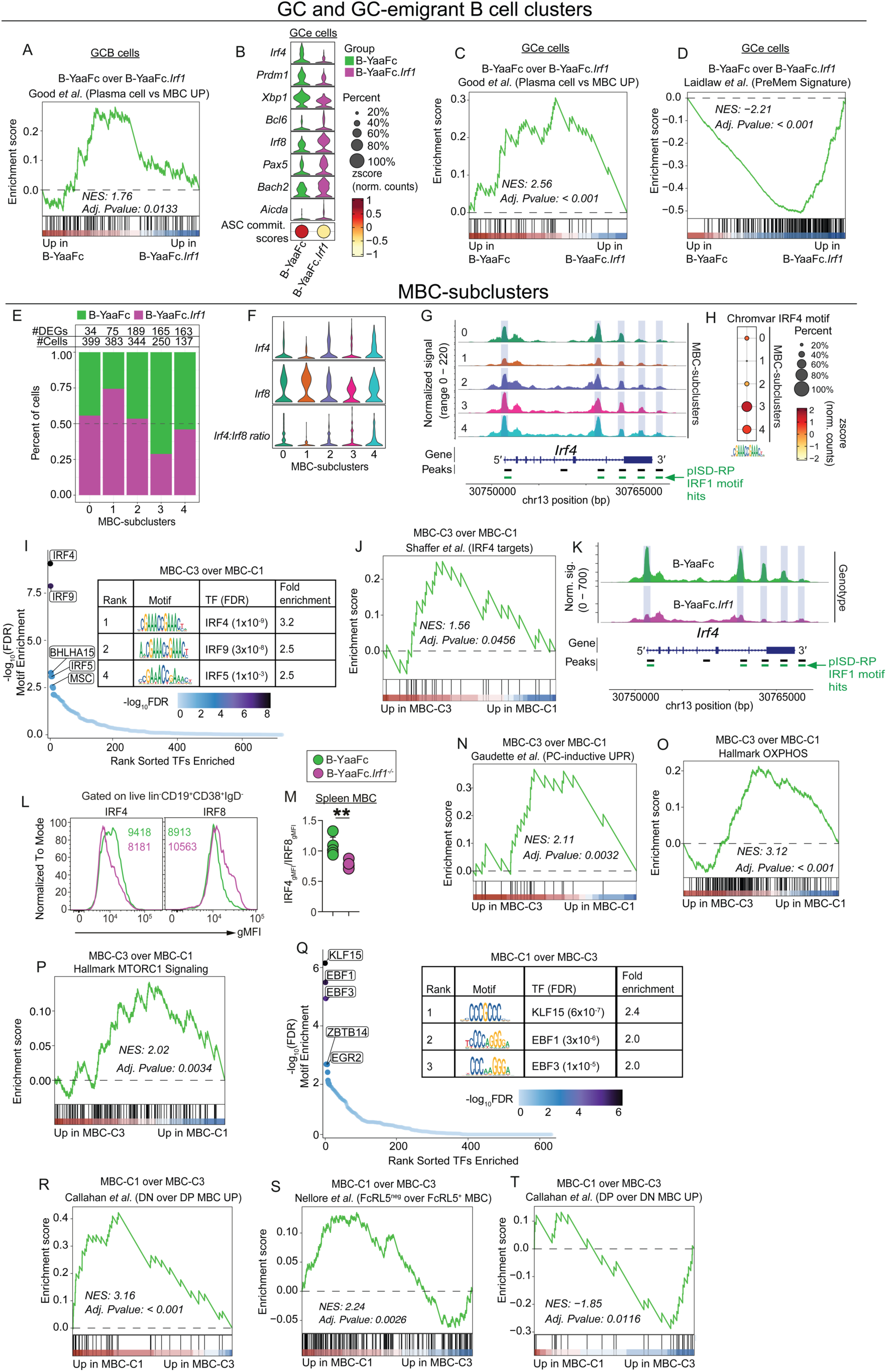
IRF1 regulates the *Irf4*:*Irf8* balance and alters effector potential of post-GCB cells from lupus-prone mice. (**A-D**) Analysis of total GCB cells (**A**) and all cells assigned as recent emigrants from the GC (GCe cells, **B-D**). GSEA (**A**) using published plasma cell vs MBC signatures^75^ gene sets to query ranked DEG list of B-YaaFc GCB cells over B-YaaFc.*Irf1^−/−^* GCB cells. Gene expression profile of B-YaaFc and B-YaaFc.*Irf1^−/−^*GCe cells (**B**, top). Expression of ASC-related genes (*Irf4*, *Prdm1*, *Xbp1*) and core B lineage genes (*Bcl6*, *Irf8*, *Pax5*, *Bach2*, *Aicda*) reported. ASC commitment scores for B-YaaFc and B-YaaFc.*Irf1^−/−^* GCe cells (**B**, bottom) were determined using Seurat AddModuleScore^79^ function with genes curated from a published dataset.^80^ GSEA (**C-D**) using published plasma cell vs MBC^75^ (**C**) and Pre-Mem^85^ (**D**) datasets to query ranked DEG list of B-YaaFc GCe over B-YaaFc.*Irf1^−/−^* GCe. See **Fig. S4A-D** for unsupervised reclustering analysis performed using all cells assigned as GCB or GCe and **Fig. S4E-F** for IRF4 motif enrichment and effector gene set enrichment analyses in B-YaaFc and B-YaaFc.*Irf1^−/−^*GCe. (**E-T**) Multiome analysis of MBC subclusters (**E-K, N-T**) and flow cytometry analysis of MBC from B-YaaFc and B-YaaFc.*Irf1*^−/−^ mice (**L-M**). (**E-K**) Unsupervised reclustering of cells assigned to the MBC lineage (See Fig. 4A and **Fig. S3D-E**). Five MBC subclusters (MBC-C0 to MBC-C4) were identified and the genotype of the cells in each MBC subcluster was determined (**Fig. S4G**). Data reported in (**E**) as the number of cells and DEG assigned to each MBC subcluster and the relative abundance of the B-YaaFc and B-YaaFc.*Irf1*^−/−^ B cells within the different MBC subclusters. *Irf4*, *Irf8* expression levels (**F**) and the *Irf4*:*Irf8* ratio for each MBC subcluster. Chromatin accessibility (**G**) surrounding the *Irf4* gene in each MBC subcluster. ChromVar^137^ IRF4 motif enrichment (**H**) by each MBC subcluster. ChromVar TF-binding motif enrichment^137^ (**I**) analysis comparing enrichment for TF motif accessibility in MBC-C3 over MBC-C1. GSEA (**J**) using an IRF4 target geneset^92^ to query the ranked DEG list of MBC-C3 over MBC-C1. Chromatin accessibility (**K**) surrounding the *Irf4* gene among all B-YaaFc and B-YaaFc.*Irf1*^−/−^ MBC. (**L-M**) Flow cytometry analysis of IRF4 and IRF8 protein expressed by splenic CD19^+^CD38^+^IgD^neg^ MBC from 6-month post-reconstitution B-YaaFc and B-YaaFc.*Irf1^−/−^* chimeras. Representative plots with geometric mean fluorescence intensities (gMFI) for IRF4 and IRF8 (**L**) shown and IRF4:RF8 protein ratio (**M**) reported. (**N-T**) GSEA using a plasma cell-inductive unfolded protein response (UPR) geneset^92^ (**N**), the Hallmark oxidation phosphorylation (OXPHOS) gene set (**O**) and the Hallmark MTORC1 signaling geneset (**P**) to query the ranked DEG list of MBC-C3 over MBC-C1. TF-binding motif enrichment analysis (**Q**) of MBC-C1 over MBC-C3. GSEA (**R-T**) using published datasets of genes upregulated in mouse stem-like CD80^neg^PD-L2^neg^ (DN) MBC relative to effector-like CD80^+^PD-L2^+^ (DP) MBC^104^ (**R**); genes upregulated in stem-like FcRL5^neg^ human MBC relative to effector-like FcRL5^+^ MBC^100^ (**S**); and genes upregulated in mouse effector-like DP MBC relative to stem-like DN MBC^104^ (**T**); to query ranked DEG list of MBC-C1 over MBC-C3 cells. Adjusted p value and NES scores for each GSEA (**A**, **C-D**, **J**, **N-P**, **R-T**) indicated (see also **Table S3**). TF motif enrichment (**I, Q**) was determined using Likelihood Ratios-based DARs using nCount_ATAC as a latent variable. Top 5 motifs are shown. See **Fig. S4I-J** for additional analysis of the MBC subclusters, **Fig. S4H** for IRF4:IRF8 protein ratio for medLN MBC. See **Table S3** for gene list used for ASC commitment scores.^80^ See **Table S4** for number of GCB subclusters based on WNN (RNA+ATAC) analysis, DEG lists for GCB cell GCe cell and MBC, average expression of select genes among all GCe cells by genotype, average expression of select genes among MBC subclusters, and motif enrichment analysis results. DEGs identified using auROC and Wilcoxon p-value based on Gaussian approximation. Genes with |avg_log2FC| ≥0.1, FDR <0.05 and expressed in ≥10% of cells per cluster or group were considered significant. **p < 0.01.

Next, we examined cells in the cluster that appeared to represent recent GC emigrants^76^ that lacked *Ighd*, maintained a follicular/GC positioning signature (*Cxcr5*, *Gna13*), but had downregulated canonical GCB cell genes (e.g. *Bcl6* and *Aicda*) and upregulated GC-exit genes (e.g. *Bach2*, *Ly6d*, and *Cd38*) (Cluster B-C9, Fig. S3D-E). Reclustering of these putative recent GC emigrants, resolved four distinct subclusters (Fig. S4C, GCe-C0 to GCe-C3). Similar to the GC compartment, the GC emigrant B cell compartment was dominated by B-YaaFc.*Irf1*^−/−^ cells (see Fig. 4C) and all three GCe subclusters were enriched for B-YaaFc.*Irf1*^−/−^ cells (Fig. S4C-D). Therefore, we compared the transcriptomes and epigenomes of all GCe cells divided on the basis of genotype (B-YaaFc GCe versus B-YaaFc.*Irf1*^−/−^ GCe) (Table S4). GCe cells from B-YaaFc mice expressed relatively higher levels of TFs like *Prdm1* and *Xbp1* that are required for terminal ASC differentiation and robust Ab production^77^ (Fig. 5B, Table S4), while GCe cells from B-YaaFc.*Irf1*^−/−^ mice expressed higher levels of *Bach2* and *Pax5* (Fig. 5B, Table S4) – TFs that are important for MBC development and maintenance.^78^

Since the data suggested that the recent GC emigrant cells from the B-YaaFc.*Irf1*^−/−^ mice might be biased toward a MBC fate, we hypothesized that *Irf4* transcript levels should be decreased in these cells and that we might also see changes in *Irf8*, which functions to restrain the IRF4-driven ASC program.^40^ Consistent with this hypothesis, GCe cells from B-YaaFc.*Irf1*^−/−^ mice expressed lower *Irf4* but higher *Irf8* levels relative to the GCe cells from B-YaaFc mice, which expressed higher levels of *Irf4* and lower levels of *Irf8* (Fig. 5B, Table S4). Not surprisingly, the higher *Irf4* levels in GCe cells from B-YaaFc mice was accompanied by increased accessibility surrounding IRF4-binding motifs within the genome of the B-YaaFc cells (Fig. S4E).

Given these results, we postulated that the GCe from B-YaaFc mice would be enriched for a pre-ASC transcriptional signature while the GCe from B-YaaFc.*Irf1*^−/−^ mice would exhibit a pre-MBC signature. In agreement with our prediction, the ASC commitment gene score based on Seurat AddModuleScore^79^ function using a published dataset^80^ was significantly higher in GCe cells from B-YaaFc mice compared GCe cells from B-YaaFc.*Irf1*^−/−^ mice (Fig. 5B, Table S3) and GSEA using public dataset GSE13411^75^ to compare the transcriptome of GCe cells from B-YaaFc mice and B-YaaFc.*Irf1*^−/−^ mice revealed that B-YaaFc GCe cells were enriched for expression of genes that are upregulated in ASC relative to MBC (Fig. 5C, Table S3). Likewise, GCe cells from B-YaaFc mice were significantly enriched for gene programs required for robust ASC differentiation and function, such as OXPHOS,^69,81^ MYC targets,^82^ UPR,^83,84^ and MTORC1^84^ (Fig. S4F, Table S3). In striking contrast, GCe cells from B-YaaFc.*Irf1*^−/−^ mice were enriched for genes expressed by PreMem cells (Fig. 5D, Table S3), which represent the transcriptionally and functionally distinct GC-derived MBC precursors that are localized near the GC border.^85–87^ Consistent with this observation, GCe cells from B-YaaFc.*Irf1*^−/−^ mice were negatively enriched for genes that are upregulated in ASC compared to GCB cells^80^ (Fig. S4F). Finally, GCe cells from B-YaaFc.*Irf1*^−/−^ mice were enriched for genes that are upregulated in *Prdm1*^−/−^ and *Xbp1*^−/−^ (Fig. S4F, Table S3) B cells.^88^ Collectively, these data suggest that the recent GC emigrants from B-YaaFc mice lie on a trajectory that tilts toward terminal ASC differentiation, whereas the recent GC emigrants from B-YaaFc.*Irf1*^−/−^ mice are biased away from the ASC lineage toward an MBC fate.

### *Irf1* regulates the IRF4:IRF8 balance in effector and stem-like MBC from lupus-prone mice

The mouse MBC compartment is heterogenous^89^ and composed of effector CD80-expressing MBC, which can rapidly differentiate into ASC,^90,91^ and more stem-like CD80^neg^PDL2^neg^ MBC, which can re-enter GC reactions.^90^ Given our results showing that deletion of *Irf1* in B cells affected the transcriptional fate potential of post-GC emigrant B cells, we hypothesized that *Irf1* might not only regulate the GC fate decision between the MBC and ASC lineages but might also control the balance between the effector and stem-like MBC subsets. To test this, we reclustered all MBC from the multiome dataset (B-C3, B-C13, B-C14 clusters, Fig. S3D-E) and identified five distinct MBC subclusters (MBC-C0 to MBC-C4), which contained 137-399 cells per subcluster and 34-189 DEGs per subcluster (Fig. S4G, Table S4). While the relative proportions of B-YaaFc and B-YaaFc.*Irf1*^−/−^ cells were very similar within subclusters MBC-C0, MBC-C2 and MBC-C4, the MBC-C3 subcluster was dominated by B-YaaFc cells and the MBC-C1 cluster was predominantly composed of B-YaaFc.*Irf1*^−/−^ cells (Fig. 5E). The B-YaaFc biased MBC-C3 subcluster expressed higher levels of *Irf4* and the lowest levels of *Irf8* (Fig. 5F, Table S4). By contrast the YaaFc.*Irf1*^−/−^ dominated MBC-C1 subcluster expressed lower *Irf4* and higher *Irf8*, resulting in the lowest *Irf4*:*Irf8* ratio (Fig. 5F, Table S4). Consistent with these results, chromatin accessibility surrounding IRF1 motif-containing *Irf4* DARs was significantly increased in MBC-C3 cells relative to the MBC-C1 cells (Fig. 5G).

Since IRF1 appeared to regulate *Irf4* expression in B-YaaFc MBC, we predicted that IRF4 would be more highly expressed in B-YaaFc MBC and would drive transcriptional programming in these cells. Indeed, accessible IRF4 binding motifs were highly enriched in the B-YaaFc dominated MBC-C3 and were almost completely absent in MBC-C1 cells (Fig. 5H). Moreover, the IRF4 binding motif was the most enriched accessible TF binding motif when comparing MBC-C3 and MBC-C1 cells (Fig. 5I, Table S4) and GSEA using a curated list of IRF4 target genes in B cells^92^ revealed significant enrichment for expression of IRF4-induced genes in MBC-C3 cells relative to MBC-C1 cells (Fig. 5J, Table S3). These results were not limited to the MBC found in the MBC-C3 and MBC-C1 subclusters as dividing all MBC on the basis of genotype revealed increased chromatin accessibility within the *Irf4* gene in the B-YaaFc MBC relative to the B-YaaFc.*Irf1*^−/−^ MBC (Fig. 5K). Flow cytometry analysis of splenic (Fig. 5L-M) and LN (Fig. S4H) MBC showed that B-YaaFc.*Irf1*^−/−^ MBC expressed lower levels of IRF4 protein and higher levels of IRF8 protein relative to B-YaaFc cells.

Given these results, we hypothesized that the B-YaaFc MBC should be more effector-like and would be enriched for expression of gene programs that support ASC differentiation and function. We observed that binding motifs for other IRF TFs, including IRF5 and IRF9, which are involved in ASC development and function,^35,36,93^ exhibited increased accessibility in MBC-C3 cells compared to MBC-C1 cells (Fig. 5I, Table S4). MBC-C3 cells also upregulated genes that promote or support B cell proliferation and rapid ASC differentiation (Table S4), including *Zeb2*, *St6gal1* and *Dock2,*^73,94,95^ and downregulated genes critical for maintenance of B cell identity, like *Pax5*, *Cr2*, *Foxp1*, *Ebf1*, *Bach2* and *Bank1*.^96–99^ MBC-C3 cells were also positively enriched for the expression of Hallmark gene sets that are upregulated in effector MBC poised for ASC differentiation,^91,100^ including pathways controlling the plasma cell-inductive UPR,^83,84^ MTORC1 signaling,^84^ and mitochondrial oxidative phosphorylation^69,81^ (Fig. 5N-P, Table S3). Finally, MBC-C3 cells were enriched for the expression of effector MBC and effector T cell genes^100–103^ (Fig. S4I, Table S3) and expressed the highest level of *Cd80* (Fig. S4J, Table S4) – a known marker of effector MBC.^90,91,104^

In striking contrast, motifs for TFs that suppress effector B cell potential including EBF1/EBF3, which maintain B-cell identity and antagonize PRDM1/XBP1-driven plasmablast programs,^96^ and KLF family TFs (KLF4, KLF15, KLF16), which enforce B cell quiescence programs,^105^ were enriched in MBC-C1 cells compared to MBC-C3 cells (Fig. 5Q, Table S4). MBC-C1 cells also expressed higher levels of genes that oppose ASC differentiation (Table S4), including *Pax5*, *Foxp1, Ebf1*, and *Bank1*. Finally, GSEA using published datasets^100,104^ to compare MBC-C1 cells to MBC-C3 cells revealed that MBC-C1 cells were positively enriched for the expression of genes that are upregulated in resting or stem-like MBC^100,104^ (Fig. 5R-S, Table S3) and were negatively enriched for expression of effector-like MBC associated genes^104^ (Fig. 5T, Table S3). Thus, IRF1 appears to facilitate development of effector MBC poised for rapid ASC differentiation while MBC lacking *Irf1* take on the transcriptome and epigenome profiles of resting or stem-like MBC.

### *Irf1* controls expansion and differentiation of autoreactive mouse B cells into ASC

The mouse single cell multiome data suggested that IRF1 controls autoimmune B cell fate decisions, at least in part, by regulating the IRF4-RF8 balance that dictates B cell identity and ASC fate programming. To determine whether these IRF1-directed transcriptional and epigenetic changes were linked to B cell functional attributes, we sort-purified follicular B cells (FOB, CD19⁺CD21^lo^CD23^+^PNA^−^CD138^−^) from spleens of 3 month old YaaFc and YaaFc.*Irf1*^⁻/⁻^ mice, labeled the cells with Cell Trace Violet (CTV) and cultured the cells *in vitro* in the presence of anti-Kappa (Fab’2), IFNγ, TLR7/8 ligand (R848) and cytokines (IL-2, IL-21) that support B cell proliferation and differentiation.^91^ Flow cytometric analysis on D4 cultures revealed that FOB cells from both groups proliferated, however YaaFc.*Irf1*^⁻^*^/^*^⁻^ FOB cells divided and expanded less compared to YaaFc FOB cells (Fig. 6A-D). Not surprisingly, given that proliferation of activated B cells is required for optimal ASC differentiation,^106,107^ there was a marked reduction in the frequency and number of CD138⁺ (Fig. 6E-G) and CD93⁺CD138⁺ (Fig. 6E, H-I) ASC in cultures derived from YaaFc.*Irf1*^⁻^*^/^*^⁻^ mice compared to YaaFc mice.

**Figure 6.**
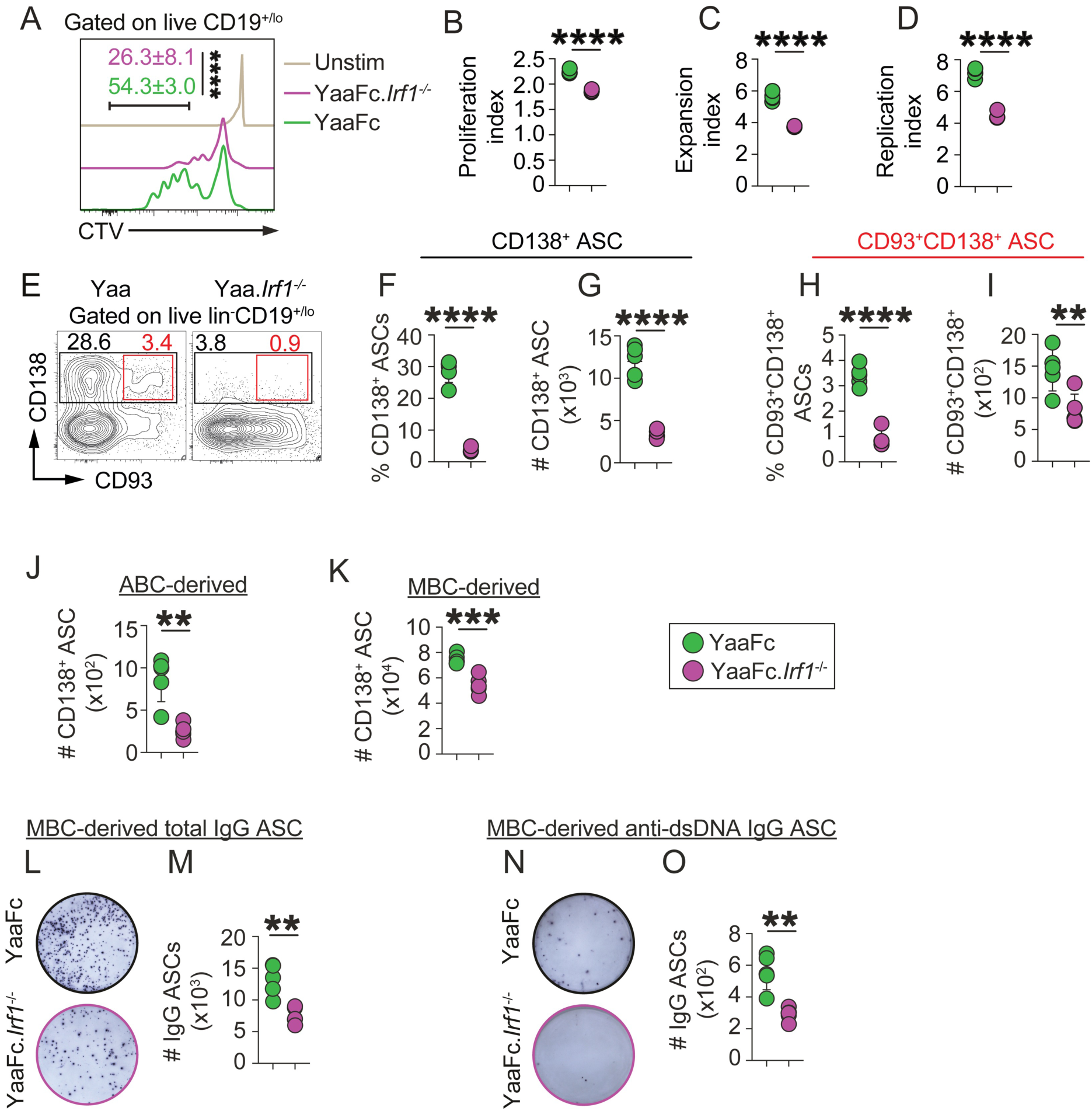
IRF1 controls expansion and differentiation of autoreactive mouse MBC and ABC into ASC. (**A-O**) Proliferation and differentiation analysis of sort-purified CD19⁺CD21^lo^CD23^+^PNA^−^CD138^−^ follicular B (FOB) cells (**A-I**), sort-purified CD19⁺CD21^−^CD23^−^CD11c^+^PNA^−^CD138^−^ ABC (**J**) and sort-purified CD19⁺CD21^lo^CD23^+^CD38^+^IgD^−^PNA^−^CD138^−^ MBC (**K-O**) isolated from spleens of 3-month YaaFc and YaaFc.*Irf1*^⁻^*^/^*^⁻^ mice. Purified B cell subsets were labeled with CellTrace Violet (CTV, **A-D**) and cultured under ASC-inducing conditions (anti-Kappa (Fab’2), IL-2, IL-21, R848, IFNγ). Cell expansion (**A-D**) and ASC differentiation (**E-O**) measured on D4. (**A-D**) Representative CTV dilution profiles from cultured FOB on D4 (**A**), showing average percentage of cells that had undergone ≥ 5 divisions. Data are reported as proliferation index (**B**, total number divisions / number of cells that divided); expansion index (**C**, total cell number / number of cells at start of culture); and replication index (**D**, total number of divided cells / number of cells that entered division). (**E-I**) Flow cytometric analysis of ASC recovered from cultures initiated with FOB cells (**E-I**), ABC (**J**), and MBC (**K**). Representative flow cytometry plots (**E**) identifying CD138^+^ and CD138^+^CD93^+^ ASC in cultures. Data reported as frequencies (**F**) and absolute numbers (**G, J, K**) of CD138^+^ ASC in the cultures or frequencies (**H**) and absolute numbers (**I**) of CD93^+^CD138^+^ ASC in the cultures. (**L-O**) ELISPOT enumeration of total IgG^+^ (**L-M**) and dsDNA-specific IgG^+^ (**N-O**) ASC from cultures initiated with MBC. Data reported as representative ELISPOTs (**L, N**) and absolute numbers (**M, O**) of IgG^+^ and dsDNA-specific IgG^+^ ASC. Data shown from individual mice (**B-D, F-K, M, O**). Data representative of two independent experiments (n ≥ 4 mice/group). Significance determined by unpaired two-tailed Student’s t tests. **p < 0.01, ***p < 0.001, ****p < 0.0001.

Since our data showed that *Irf1* regulates the transcriptional and epigenomic profiles of mouse MBC and ABC, we asked whether IRF1 specifically controls the differentiation potential of these cells. We sorted splenic YaaFc or YaaFc.*Irf1*^⁻^*^/^*^⁻^ MBC (CD19⁺CD21^lo^CD23^+^CD38^+^IgD^−^PNA^−^CD138^−^) and ABC (CD19⁺CD21^−^ CD23^−^CD11c^+^PNA^−^CD138^−^), stimulated the cells with anti-Kappa (Fab’2), R848, IFNγ, IL-2, and IL-21 and enumerated CD138⁺ ASC in the D4 cultures. We observed significantly fewer ASC in YaaFc.*Irf1*^−/−^ MBC and ABC cultures compared to cultures containing YaaFc MBC and ABC (Fig. 6J-K). To confirm these results, we performed ELISPOT assays on D4 MBC cultures and observed that YaaFc.*Irf1*^−/−^ MBC generated significantly fewer IgG⁺ ASC relative to YaaFc MBC (Fig. 6L-M). Moreover, the number of anti-dsDNA-IgG⁺ secreting ASC was significantly decreased in the cultures containing Yaa.*Irf1*^⁻^*^/^*^⁻^ MBC (Fig. 6N-O). Thus, IRF1 appears to directly support expansion and differentiation of effector autoimmune-prone mouse ABC and MBC following exposure to TLR7 ligand and IFNγ.

### *IRF1* expression correlates with IRF4-driven effector programs in human SLE B cells

Our data using an SLE mouse model indicated that IRF1 modulates the expression and activity of the ASC-commitment TF, IRF4. To test whether IRF1 might be required for *IRF4* induction in human B cells, we interrogated a published^108^ single-cell RNA-seq dataset from two patients carrying a nonsense mutation in the *IRF1* coding region. Consistent with our mouse B cell data, IgD^neg^ B cells from *IRF1*-deficient patients displayed markedly reduced *IRF4* expression (Fig. 7A). Next, to address whether the IRF1-dependent circuitry mapped in our TLR7-driven lupus mouse model was activated in human SLE patient B cells, we compared the transcriptional and chromatin landscapes of peripheral B cells from SLE patients with healthy control donors (HC).^109^ Analysis of bulk B cell RNA-seq profiles revealed an increase in *IRF1* mRNA in the SLE B cells relative to HC B cells (Fig. 7B). Consistent with this, chromatin accessibility surrounding IRF1 ISRE (interferon-stimulated response elements) binding motifs was increased in the SLE B cells (Fig. 7C). When the total B cells were divided into discrete subsets, we observed increased *IRF1* mRNA in the SLE-derived resting naïve (rN), transitional 3 (T3), activated naïve (aN), switched MBC (SM) and DN2 B cells (Fig. S5A). SLE B cells also exhibited increased *IRF4* mRNA expression in total B cells (Fig. 7D) and within the rN, T3, aN, and SM subsets relative to the HC counterparts (Fig. S5B). We further observed a significant positive correlation between *IRF1* and *IRF4* levels in SLE B cells but not HC B cells (Fig. S5C), suggesting that *IRF1* and *IRF4* may be co-regulated in SLE B cells.

**Figure 7.**
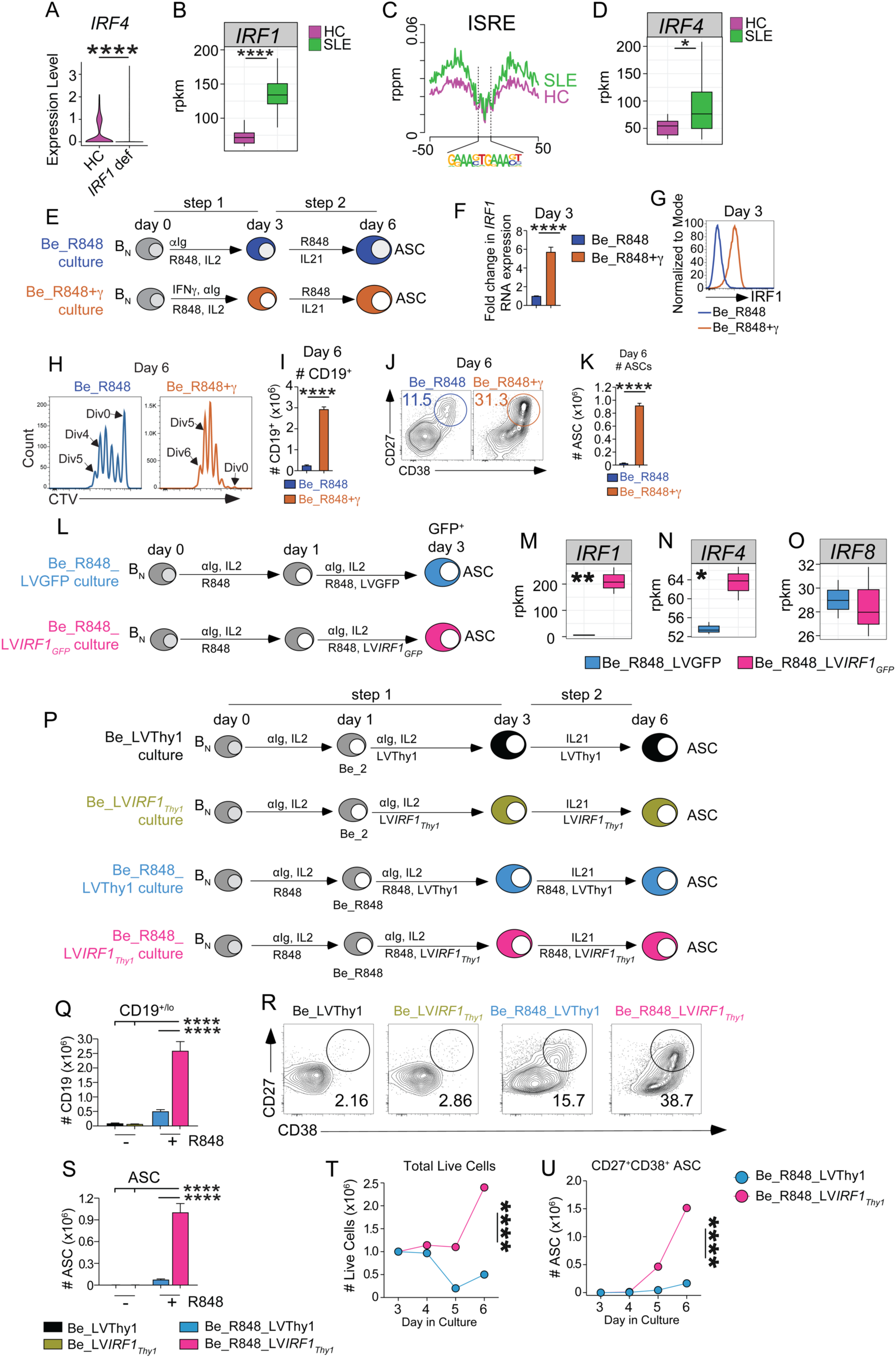
IRF1 regulates *IRF4* expression and synergizes with TLR7 signals to support IFNγ-dependent differentiation of human B cells into ASC. (**A-D**) Analysis of published^108,109^ human B cell data sets showing: (**A**) *IRF4* expression by B cells from *IRF1*-deficient individuals and HC (HC=healthy control donors); (**B**) *IRF1* expression by B cells from HC and SLE patients; (**C**) chromatin accessibility surrounding the interferon-stimulated response element (ISRE) motif (IRF1-binding site) in HC and SLE B cells; and (**D**) *IRF4* expression by HC and SLE B cells. (**E-K**) Analysis of IFNγ- and TLR7-dependent *in vitro* human B cell differentiation cultures. Experimental design schematic (**E**) showing Step 1 (activation) and Step 2 (proliferation and differentiation). Step 1: Activation of CTV-labeled primary tonsil-derived HC naïve B cells with anti-IgM+IL2+R848 (Be_R848 cultures) or with anti-IgM+IL2+R848, and IFNγ (Be_R848+γ cultures). Step 2: Cells from both cultures washed on D3, restimulated with IL21 and R848 and analyzed on D6. *IRF1* mRNA (**F**) in D3 cells, quantitated by qPCR and normalized to *GAPDH.* IRF1 protein levels analyzed by flow cytometry (**G**) in D3 cells. Cell proliferation measured by CTV dilution (**H**) and reported as cell recovery (**I**) on D6. Enumeration of ASC by flow cytometry on D6. Data shown as representative flow cytometry plots with frequency (**J**) and absolute number (**K**) of CD27^+^CD38^+^ ASC reported. (**L-O**) Bulk RNAseq analysis of *in vitro* human B cell differentiation cultures. Experimental design schematic (**L**) showing lentiviral (LV) transduction of primary tonsil-derived HC naive B cells activated with anti-IgM+IL2+IL21+R848. On D1, cells were transduced with a lentiviral vector expressing either GFP (Be_R848_LVGFP) or with a lentiviral vector encoding *IRF1* and GFP (Be_R848_LV*IRF1*_GFP_). Bulk RNAseq performed on D3. Gene expression levels for *IRF1* (**M**), *IRF4* (**N**) and *IRF8* (**O**). (**P-S**) Analysis of IRF1- and TLR7-dependent *in vitro* human B cell differentiation cultures. Experimental design schematic (**P**) showing Step 1 (activation) and Step 2 (proliferation and differentiation) in LV transduced cells. Step 1: primary tonsil-derived HC naive B cells were activated with anti-IgM+IL2 (Be) and transduced on D1 with LV encoding mouse Thy1.1 (Be_LVThy1 (black)) or with LV encoding IRF1 and Thy1 (Be_LV*IRF1*_Thy1_ (olive green) cultures). Alternatively, naïve B cells were activated with anti-IgM+IL2+R848 and then transduced with the control LV (Be_R848+LVThy1, (blue)) or with the *IRF1* LV (Be_R848_LV*IRF1*_Thy1_ (pink) cultures). Step 2: D3 cells from the 4 cultures were washed and transduced with the same LV used in Step 1 and then stimulated with IL21 (Be_LVThy1 and Be_LV*IRF1*_Thy1_ cultures) or with IL21+R848 (Be_R848_LVThy1 and Be_R848_LV*IRF1*_Thy1_ cultures). Cells from D6 cultures were analyzed by flow cytometry (**Q-S**). Data reported as total number of recovered CD19^+/lo^ cells from the four cultures (**Q**), with representative flow cytometry plots showing the frequencies (**R**) and absolute numbers (**S**) of CD27^+^CD38^+^ ASC recovered on D6 from cultures initiated with 1×10^6^ naïve B cells. (**T-U**) Analysis of Be_R848_LVThy1 and Be_R848_LV*IRF1*_Thy1_ cultures containing sorted D3 Thy1^+^ cells labeled with CTV (see **Fig. S6J**). Absolute numbers of total live cells (**T**) and CD27^+^CD38^+^ ASC (**U**) recovered from D3, D4, D5 and D6 cultures shown. Data shown as mean ± SD (**B, D, I, K, M-O, Q, S**) or mean ± SEM (**F**). n=10 SLE and n=8 HC (**B-D**). Data in (**B, D, M-O**) are shown as reads per kilobase per million reads (rpkm) values. Data in (**M-O**) are from 3 independent cultures that were set up with donor-matched samples. Center lines of boxplots indicate the median, lower and upper bounds of boxes indicate the 1^st^ and 3^rd^ quartiles and whiskers indicate the upper and lower limits of the data. Significance determined by Wilcoxon rank sum test (**A**), unpaired two-tailed Student’s t tests (**B, D, F, I, K, M-O**), one-way ANOVA followed by Tukey’s post hoc multiple comparison test (**Q, S**) or by comparing time course area under the curve (AUC) values between R848_LVGFP and Be_R848_LV*IRF1* cultures using unpaired two-tailed Student’s t test (**T-U**). See **Fig. S5** for additional analysis of SLE and HC RNA-seq data. See **Fig. S6** and **Table S5** for additional analyses of the human *in vitro* B cell cultures. *p < 0.05, **p ≤ 0.01, ****p ≤ 0.0001.

To corroborate this result, we analyzed a published^110^ single-cell RNA-seq dataset containing >120,000 non-ASC peripheral B cells from 99 HC and 162 SLE patients. Similar to what we observed with the SLE-prone mouse B cells, GSEA comparing SLE and HC B cells showed that SLE B cells were transcriptionally skewed toward effector-like B cell programs (Fig S5D, Table S3). The SLE B cells also expressed higher levels of *IRF1* (Fig. S5E) and *IRF4* (Fig. S5F) transcripts relative to B cells from HC. Furthermore, we observed a significantly stronger positive correlation between *IRF1* and *IRF4* expression levels in the *IRF1*-expressing SLE B cells (r = 0.56, p = 7.7×10^−31^) relative to the *IRF1*-expressing HC B cells (r = 0.21, p = 2.2×10^−3^) (Fig S5G). Together, these findings show that elevated *IRF1* expression in SLE B cells is associated with increased *IRF4* expression and acquisition of an effector B cell program.

### IFNγ-induced IRF1 licenses TLR7-dependent naïve human B cell differentiation into ASC

The human B cell transcriptome data sets showed that *IRF1* was expressed and upregulated in SLE patient B cells that exhibited effector-like programming. To test whether IRF1 regulates human B cell proliferation and differentiation in the setting of inflammation, we utilized our previously established 2-step activation (step 1), and proliferation + differentiation (step 2) *in vitro* B effector (Be) culture system^15^ (Fig. 7E). This B cell differentiation culture system requires core stimuli (anti-IgM + IL2 + IL21) and additionally requires IFNγ and TLR7 ligand (R848) in step 1 and R848 in step 2. To address whether IRF1 was upregulated in human B cells in an IFNγ-dependent fashion, we activated CTV-labeled HC naïve B cells for 3 days (Fig. 7E) with anti-IgM, IL-2, and R848 (Be_R848) or with anti-IgM, IL-2, R848, and IFNγ (Be_R848+γ). We observed that IFNγ signals, provided during the activation phase, resulted in markedly increased *IRF1* mRNA and protein expression by the Be_R848+γ cells compared to Be_R848 cells (Fig. 7F-G). Consistent with our previous report that IFNγ, but not IFNα, is a major inducer of IRF1 in B cells,^48^ we observed a minimal change in *IRF1* mRNA and protein expression in cells receiving IFNα in place of IFNγ (D3 Be_R848+α cultures, Fig. S6A-B).

Next, we asked whether IRF1, induced by IFNγ signals, was associated with B cell proliferation. As expected,^15^ minimal proliferation and ASC differentiation was seen in either culture during the 1^st^ step activation phase (Fig. S6C-D). By the end of the 2^nd^ phase, the B cells from both cultures had undergone multiple rounds of division (Fig. 7H), however more late division (Div. 5+) cells accumulated in the Be_R848+γ cultures compared to the Be_R848 cultures (Fig. 7H) and significantly more B cells were recovered in the Be_R848+γ cultures (Fig. 7I). Consistent with this enhanced proliferative response, Be_R848+γ cultures yielded substantially higher frequencies and numbers of ASC compared to the Be_R848 cultures (Fig. 7J-K, Fig. S6E). The IFNγ-dependent accumulation of ASC in the culture was also highly dependent on TLR7 signals as removal of R848 from Steps 1 and 2 of the Be_R848+γ cultures prevented ASC development (Fig. S6F). By contrast, higher concentrations of R848 synergized with IFNγ signals to support maximal ASC development (Fig. S6F). Thus, IFNγ licensed TLR7-stimulated human naïve B cells for proliferation and ASC differentiation and did so in parallel with IRF1 upregulation.

### IRF1 cooperates with TLR7 signaling to promote human B cell proliferation and differentiation

Our data showed that IFNγ-induced *IRF1* expression in human B cells was associated with ASC development. To address whether *IRF1* expression in human B cells was sufficient to tip the IRF4-IRF8 balance toward ASC commitment in the absence of IFNγ, we activated primary tonsil-derived human naive B cells with anti-IgM+IL2+IL21+R848. On D1, we transduced the cells with a lentiviral vector expressing either GFP (Be_R848_LVGFP cultures) or with a lentiviral vector encoding *IRF1* and GFP (Be_R848_LV*IRF1*_GFP_, Fig. 7L). We then performed bulk RNAseq on both cultures on D3 – at a timepoint before the cells substantially expand or differentiate into ASC (Fig. S6C-D). This allowed us to identify the immediate-early transcriptional circuitry engaged by IRF1 in activated B cells without the confounding influence of changes in cell number or overt effector differentiation. Consistent with the fact that IFNγ was not added to the cultures, the bulk RNA-seq data showed that *IRF1* was not upregulated by the Be_R848_LVGFP B cells (Fig. 7M, Table S5). By contrast, we observed robust expression of *IRF1* mRNA in the Be_R848_LV*IRF1*_GFP_ cells that had been transduced with the *IRF1* lentivector (Fig 7M, Table S5). Moreover, and consistent with our mouse B cell data, enforced *IRF1* expression in the stimulated human B cells reshaped the expression of key B cell fate regulators as GSEA comparing Be_R848_LV*IRF1*_GFP_ and Be_R848_LVGFP revealed enrichment of effector gene programs in D3 Be_R848_LV*IRF1*_GFP_ cells (Fig. S6G, Table S3). Furthermore, Be_R848_LV*IRF1*_GFP_ cells expressed elevated *IRF4* levels compared to Be_R848_LVGFP B cells (Fig. 7N-O) and the *IRF4*-to-*IRF8* ratio was heavily biased toward *IRF4* in the Be_R848_LV*IRF1*_GFP_ cells compared to the control cells (Fig. S6H). These data therefore show that IRF1 is sufficient to induce *IRF4* upregulation and acquisition of an effector-like transcriptional program in TLR7/8 activated human B cells.

To directly test whether IRF1 was also sufficient to support IFNγ-dependent, TLR7-driven human B cell proliferation and differentiation, we stimulated primary tonsil-derived human naive B cells with the core anti-IgM+IL2+IL21 cocktail. In one culture, we removed IFNγ and R848 from the activation step 1 and in the other culture we removed IFNγ but included R848 (Fig. 7P). We transduced the cultures with a lentivirus encoding a reporter gene (LVThy1) or with a lentivirus containing IRF1 plus the Thy1 reporter gene (LV*IRF1*_Thy1_). In the second step, we re-transduced the cells with the same lentivirus the cells received in step 1 and added R848 to the cultures that received R848 during the activation phase (Fig. 7P).

Cells that were not exposed to IFNγ and R848 throughout the six day culture did not expand (Fig. 7Q) or differentiate (Fig. 7R-S) into ASC, regardless of whether the cells were transduced with the control vector (Be_LVThy1 cultures) or with the *IRF1* vector (Be_LV*IRF1*_Thy1_ cultures), indicating that enforced expression of *IRF1* was not sufficient to elicit B cell proliferation and differentiation in response to IL2 + anti-IgM and IL21. Similarly, only modest expansion (Fig. 7Q) and differentiation (Fig. 7R-S) of B cells was observed in Be_R848 cultures that were transduced with the control vector (Be_R848_LVThy1 cultures), indicating that TLR7 signals, in the absence of IFNγ, also did not support robust B cell expansion and subsequent differentiation. However, when the cells were stimulated with R848 and were transduced with *IRF1* (Be_R848_LV*IRF1*_Thy1_), the B cells underwent extensive expansion resulting in greatly increased cell recovery (Fig. 7Q) compared to cultures transduced with the control vector. This proliferative advantage was mirrored by a robust increase in the frequency and absolute number of ASC in the cultures transduced with *IRF1* (Fig. 7R-S). Given these results and our mouse data (Fig. 6A-D) showing reduced proliferation in the absence of IRF1, we predicted that IRF1 likely supported ASC differentiation, at least in part, by promoting B cell proliferation. To test this, we activated primary tonsil-derived HC naive B cells with anti-IgM+IL2+R848 (Be_R848). On D1, we transduced the cultures with a lentivirus LVThy1 (Be_R848_LVThy1) or LV*IRF1* (Be_R848_LV*IRF1*_Thy1_). On D3, we sorted reporter Thy1.1^+^ cells from both cultures, labeled the cells with CTV, re-cultured the cells with IL-21+R848 for 3 more days, and analyzed the cells for proliferation and ASC differentiation on D4, D5 and D6 (Fig. S6J). As expected, non-transduced cells did not express Thy1.1. However, > 50% of both Be_R848_LVThy1 and Be_R848_LV*IRF1*_Thy1_ cells expressed Thy1.1 on D3 (Fig. S6K). Consistent with our prediction, Thy1.1^+^ Be_R848_LV*IRF1*_Thy1_ cells underwent more proliferation compared to Thy1.1^+^ Be_R848_LVThy1 cells (Fig. 7T, Fig. S6L). Moreover, the Thy1.1^+^ Be_R848_LV*IRF1*_Thy1_ cells also differentiated more efficiently into ASC compared to Thy1.1^+^ Be_R848_LVThy1 cells (Fig. 7U, Fig. S6M). Thus, IRF1, an IFNγ-elicited TF, synergized with TLR7 signals to shift the IRF4:IRF8 rheostat and support human B cell proliferation and ASC differentiation. When taken together, the data in both mouse and human show that IRF1 appears to skew the IRF4:IRF8 balance in multiple B cell subsets, including ASC, ABC, GCB cells, GC emigrants and MBC, in favor of a terminal effector inflammatory ASC fate – often at the expense of preserving more stem-like, resting or regulatory populations. These results therefore suggest that IRF1 serves as a key molecular switch for B cell activation and effector differentiation, particularly in the setting of IFNγ-driven inflammatory autoimmune disease. The relevance of IRF1 as a central node in the autoimmune IFNγ-TLR7 axis is discussed.

## Discussion

Here, we identify IRF1 as a central transcriptional and epigenomic node that couples inflammatory IFNγ and TLR7 signaling to B cell fate decisions in the setting of systemic autoimmune disease. Using a combination of mouse and human B cell models, we demonstrated that IRF1 is both necessary and sufficient to drive the IFNγ- and TLR7-dependent expansion and differentiation of naïve B cells, ABC and MBC into ASC. Mechanistically, we showed that IRF1 shapes chromatin accessibility and transcriptional circuitry during key B cell fate decisions, most notably by tuning the balance of IRF4 and IRF8 to bias B cells toward a terminal inflammatory effector (ASC) differentiation program at the expense of quiescent, stem-like memory and regulatory ASC programs. Therefore, we argue that IRF1 serves as a molecular rheostat for B cell activation and effector differentiation in the setting of IFNγ-driven inflammatory autoimmune disease.

Prior studies showed increased systemic IFNγ levels in autoimmune patients and mice^8,9^ and at least in mice, it is appreciated that IFNγ signals are important drivers of autoAb production and tissue damage.^45,46^ Our data along with our previous reports^14,48^ demonstrate that IRF1, while not substantially induced following stimulation with IFNα or TLR ligands, is potently upregulated in response to IFNγ signals in mouse and human B cells. Thus, it was not completely unexpected to find that global deletion of *Irf1* in lupus-prone Yaa.*Fcgr2b*^−/−^ (YaaFc.*Irf1*^−/−^) mice curtailed splenomegaly, reduced ABC and ASC accumulation, decreased anti-dsDNA ASC frequencies and ANA load, and attenuated glomerular IgG deposition and renal pathology, culminating in significantly improved survival. However, these effects were not simply due to a global collapse of B cell immunity. Instead, our analysis of B cell chimeric animals revealed that *Irf1* expressed specifically by B cells was necessary for establishment of autoAb-mediated clinical disease and was intrinsically required for development of pathogenic ASC from effector-like ABC and MBC precursors. In fact, *Irf1* deficiency in YaaFc B cells shifted homeostasis of the B cell compartment away from the extrafollicular ABC and the inflammatory ASC axis that is recognized as a dominant driver of lupus-like autoimmunity^6,7,19^ toward a stem-like or resting MBC program. These data place IRF1 within the core B cell circuitry that determines whether chronic IFNγ-TLR7 stimulation will escalate into destructive humoral autoimmunity.

Our single-cell multiomics analyses revealed that IRF1 does not merely scale B cell activation globally but instead reconfigures the composition and internal wiring of antigen-experienced B cell subsets in a structured manner. Prior work established IRF4 and IRF8 as opposing regulators of B cell fate, with graded IRF4 levels driving ASC differentiation and IRF8 constraining this program and promoting GC and MBC fates.^38–40,42^ *Irf1* expression in B cells favored the emergence of ASC subsets endowed with a canonical high-output ASC program characterized by robust *Irf4* expression, ER/UPR and secretory machinery, translation and RNA-processing modules, and metabolic/stress adapters that support sustained Ab production.^77^ These cells also exhibited strong enrichment of proinflammatory type I and II interferon response signatures and increased chromatin accessibility at *Irf4* regulatory elements containing IRF1-binding motifs – all of which are consistent with an IFN-licensed, IRF1-dependent effector program. By contrast, B-YaaFc.*Irf1*^−/−^ mice lacked most ASC. Within the remaining ASC compartment, we identified a small population of B-YaaFc.*Irf1*^−/−^ ASC that were enriched for a regulatory signature^62,63^ and exhibited increased chromatin accessibility surrounding the *Il10* locus and increased *Il10* transcript levels. Thus, IRF1 appears to function as a fate-specifying switch that tilts ASC heterogeneity away from regulatory-like, restrained subsets and toward fully licensed, IFN-driven secretory ASC effectors.

A similar fate-switch could be seen in the ABC compartment, which represents an inflammatory “activated” extrafollicular B cell subset that is considered analogous to the IFNγ-driven T-bet⁺ DN2 cells that are expanded in SLE patients and are poised to rapidly differentiate into autoAb-producing plasmablasts in response to TLR7, IFNγ, and IL-21.^19,20,111^ We showed that *Irf1* regulates the size, molecular attributes, and functional properties of the mouse ABC compartment. Subclustering of ABC revealed that a transcriptionally distinct ABC subset was enriched in *Irf1*-sufficient B-YaaFc cells. This ABC subset, which was distinguished by high expression of *Tbx21* and strong enrichment of chromatin T-box and IFN response TF binding motifs, was transcriptionally similar to the IFNγ- and TLR7-driven ABC/DN2 cells that have been previously implicated in mouse and human lupus.^6,7^ Deletion of *Irf1* selectively in B cells resulted in decreased ABC in lupus-prone mice whether identified phenotypically or transcriptionally and the ABC that remained appeared enriched for an alternate transcriptional program that featured reduced effector gene expression and a diminished IFN signature. Although *Irf4* was not highly expressed by ABC, *Irf4* chromatin accessibility and gene expression was significantly elevated in *Irf1* sufficient YaaFc ABC relative to that seen in B-YaaFc.*Irf1*^−/−^ ABC. Consistent with this, ABC isolated from YaaFc.*Irf1*^−/−^ mice poorly differentiated into ASC following activation with IFNγ and TLR7 ligand. These data therefore suggest that IRF1 acts upstream of ASC formation to reprogram precursor ABC toward an IFNγ-licensed ASC effector path.

We further observed that B-YaaFc GCB cells and germinal center emigrant (GCe) cells displayed transcriptional signatures associated with ASC commitment or function. By contrast, B-YaaFc.*Irf1*^−/−^ GCB and GCe were enriched for expression of genes associated with PreMem and MBC programs.^85–87^ Consistent with *Irf1*-dependent skewing of the ASC:MBC balance in GCe cells, we observed that *Irf1* deficiency within the mature MBC compartment favored development of more stem-like, resting MBC that exhibited a low *Irf4*:*Irf8* ratio, enrichment of chromatin accessible binding motifs for EBF and KLF family factors, and upregulation of *Foxp1*, *Pax5*, *Cd55*, and *Bank1 –* all of which collectively reinforce B cell quiescence, preserve lineage identity, and brake terminal differentiation.^96–98,112,113^ However, in *Irf1* sufficient B-YaaFc mice, we found enrichment for MBC with an effector-like transcriptional profile. These MBC expressed effector-related genes such as *Cd80*, *Zeb2*, *St6gal1*, and *Irf4*, and exhibited reduced expression of *Pax5* and other TFs that maintain MBC identity.^78^ This *Irf1*-regulated MBC subset appeared transcriptionally and epigenetically poised for rapid ASC differentiation, with enrichment for Hallmark UPR, MTORC1, and OXPHOS signatures and effector B/T cell gene programs.^69,81,83,84,91,100–104^ Notably, chromatin accessibility at IRF1 binding motif-containing *Irf4* cis regulatory regions was higher in the *Irf1* sufficient MBC relative to B-YaaFc.*Irf1*^−/−^ MBC. These findings suggest that IRF1 qualitatively skews the MBC compartment toward an effector-biased, ASC-competent state. This result is consistent with recent data suggesting that MBC occupy a central, and often underappreciated, niche in SLE pathogenesis by serving as a durable reservoir of autoreactivity.^4^ While SLE pathogenesis is often framed in terms of short-lived plasmablasts and plasma cells, our findings support the concept that IFNγ-elicited IRF1 can program MBC to function as a durable “immunologic archive” of autoreactivity capable of reseeding pathogenic ASC responses.

Our mouse data indicated that IRF1 is a key node controlling IFNγ-dependent B cell fate decisions. Our analysis of human B cells strongly suggested that this IRF1-driven transcriptional network is highly conserved in B lymphocytes. Indeed, bioinformatic analysis of the transcriptome of antigen-experienced peripheral B cells from the rare individuals who lack a functional *IRF1* gene^108^ revealed that these *IRF1* defective B cells expressed reduced *IRF4* levels. By contrast, *IRF1* and *IRF4* levels in peripheral blood B cells from SLE patients were coordinately and consistently increased relative to healthy control B cells. The SLE B cells also displayed increased chromatin accessibility surrounding IRF1 binding motifs, consistent with heightened IRF1-dependent regulatory activity in SLE patient B cells. When primary human naïve B cells were activated with the defined stimulation cocktail^15^ that supports (i) acquisition of an activated DN2 profile, (ii) robust proliferation, and (iii) development of CD38^+^CD27^+^ ASC, we found that IFNγ synergized with TLR7/8 ligation to induce robust IRF1 expression. Expression of IRF1 in the absence of IFNγ signals was sufficient to support *IRF4* upregulation as well as proliferation and differentiation of human B cells. Thus, both mouse and human B cell data indicate that IFNγ licenses TLR7/8 driven proliferation and differentiation in an IRF1-dependent fashion that appears to be mediated, at least in part, through the regulation of *IRF4* expression.

Although IRF1 expression in myeloid cells can be elicited by other inflammatory signals, including IFNα,^24,25^ we observed that in both mouse^48^ and human B cells, IRF1 expression is predominantly induced by IFNγ and not by IFNα. This suggests that IRF1 specifically and selectively bridges IFNγ and TLR7/8 signals to drive B cell differentiation. In SLE patients, it is well appreciated that elevated IFNγ levels are associated with clinical disease manifestations.^8^ It is also known that elevated systemic IFNγ levels are an early predictor of SLE disease as elevated IFNγ can be observed years before clinical disease onset^10,11^ and well before IFNα levels rise in these patients.^11^ Given that IFNγ is required for the development of DN2 cells *in vitro*,^15,20^ for the differentiation of DN2 cells into ASC,^15,20^ and for development of ABC and autoAb in multiple mouse models of SLE,^45,46^ it is tempting to speculate that IRF1 is a key driver of many of these IFNγ-dependent autoimmune B cell defects. Interestingly, the human *IRF1* gene resides within the 5q31 cytokine cluster, which has been implicated as an autoimmune risk locus enriched for non-coding variants that modulate immune gene expression.^31–34^ Given our data showing that IRF1 rewires the chromatin and transcriptional circuitry of antigen-experienced B cells to favor an IFNγ-licensed effector B cell state, we postulate that IRF1 is not merely associated with autoimmune risk but may explain how some genetic variants within the 5q31 region could contribute to IFNγ-driven pathogenic human B cell fates.

In summary, our study reveals IRF1 as a pivotal IFNγ-inducible TF that rewires the transcriptional and epigenomic landscape of antigen-experienced B cells to favor IFNγ-licensed ASC, ABC, and effector MBC subsets that sustain systemic autoimmunity. By tuning the IRF4:IRF8 balance and remodeling chromatin at key fate-determining loci, IRF1 biases B cell responses toward terminal effector trajectories at the expense of resting, stem-like MBC and regulatory ASC programs. These insights not only reposition IRF1 as a central node in autoimmune B cell biology but also nominate the IRF1-IRF4 axis as a promising target for next-generation therapies aimed at durably reprogramming humoral immunity in lupus and related autoimmune diseases.

### Limitations of the study

First, our mechanistic analysis showing IRF1 regulation of B cell fate decisions was anchored in a single TLR7-driven mouse model of lupus. Although the results of this study were largely recapitulated with *in vitro* analyses of human B cells, it will be important to extend the molecular analyses to additional lupus and autoimmune B cells and to tissue-resident B cell niches (e.g., kidney, lung, tertiary lymphoid structures) in order to define the full spectrum of IRF1 activity in B cells. Second, while our studies focused on B cell-intrinsic roles for IRF1 in autoimmunity, IRF1 also modulates myeloid, T cell, and stromal compartments that shape the inflammatory milieu and modulate B cell behavior. In the future, it will be important to integrate B cell-targeted and global immune system perturbations in IRF1 to fully understand how IRF1 orchestrates systemic immunity in SLE. However, we did observe the loss of B cell-directed autoimmune responses in animals that were globally deficient in *Irf1*, suggesting that therapies targeting the IFNγR and IRF1 nexus would be expected to attenuate the autoreactive B cell response independent of the impact of the therapy on other cell types. Third, we did not directly address whether the IRF1-linked circuit to ABC and ASC formation is active during canonical acute immune responses. We suspect it may not be, since we did not observe a requirement for B cell-intrinsic IRF1 expression for IgG ASC and Ab response after acute influenza infection.^48^ Consistent with this, individuals with inherited IRF1 deficiency^108^ reported no history of severe acute viral disease, including SARS CoV 2. However, given that B cell-specific IRF1 deficiency did attenuate class-switched ASC responses in mice chronically infected with gammaherpesvirus,^114^ we speculate that sustained or chronic TLR7 and IFNγ signaling may be required to keep IRF1 levels elevated and skew the B cell fate toward effector ASC and ABC. Fourth, although our data bridge IRF1 function from mouse to human, larger and longitudinal human studies will be needed to determine how IRF1-dependent B cell states track with clinical phenotypes, treatment responses, and relapse risk in diverse autoimmune patient populations. Finally, while our multiomics analyses strongly implicate direct IRF1 engagement at *Irf4* regulatory regions, we know that *Irf1* deficiency in B cells alone results in global changes in the epigenome of lupus-prone YaaFc B cells. Thus, we anticipate that IRF1 will more broadly reprogram B cells. Future experiments using IRF1 ChIP-seq or CUT&RUN/CUT&Tag to map the direct IRF1 engagement at *Irf4* and other regulatory regions will be key to understanding how IRF1 integrates IFNγ and TLR7/8 signaling in autoimmunity.

## RESOURCE AVAILABILITY

### Lead Contact

Requests for further information and resources should be directed to and will be fulfilled by the lead contact, Frances E. Lund.

### Materials Availability

All reagents and mouse strains will be made available on request after completion of a Materials Transfer Agreement.

### Data and Code Availability

The single cell multiome datasets and bulk RNAseq datasets generated in this study have been deposited in the Gene Expression Omnibus (GEO) under accession numbers XXX and YYY. The datasets will be publicly available upon publication of this study. This study did not generate custom analysis code.

## ACKNOWLEDGEMENTS

We thank Thomas Scott Simpler, Shanrun Liu (Associate Director of the UAB Single Cell Core), Vidya Sagar Hanumanthu (Associate Director of the UAB Flow Cytometry Core), Chiao-Wang Sun (UAB Single Cell Specialist), Madhubanti Basu (UAB Single Cell Specialist) and the Emory Integrated Genomics Core (RRID:SCR_023529) for technical support. Funding for experiments was provided by the NIH: R01 AI110508 and R01 AI53365 (to FEL), R01 AI150664 and R01 AI152476 (to TDR), and P01 AI125180 (to FEL, CDS and TDR). NIH P30 CA013148, P30 AI027767, and NIH S10 OD032296 provided support for the UAB Flow Cytometry and Single Cell Core. G20RR022807-01 provided support for the UAB Animal Resources Program X-irradiator. EWO is supported by the UAB Heersink School of Medicine STAR21 Award.

## AUTHOR CONTRIBUTIONS

FEL and TDR conceived the idea for the project and secured initial funding. FEL and TDR supervised all UAB authors. FEL, EWO and EZM designed the experiments that were performed by EWO, AD, SC, EZM and JNP. JBF performed the histology analysis. GY made the lentivirus constructs. RB and KB provided technical support for animal experiments. Bioinformatics analyses were performed by EWO with guidance from FEL and CDS. All data were analyzed by EWO, EZM, and FEL. EWO and FEL wrote the manuscript and prepared final figures. Critical feedback on the project and manuscript was provided by all authors, and all authors have reviewed the manuscript.

## DECLARATION OF INTERESTS

The authors declare no competing interests.

### Experimental model and study participant details

#### Human subjects

Remnant human tonsil tissue was obtained by the University of Alabama at Birmingham (UAB) O’Neal Comprehensive Cancer Center Tissue Procurement Core Facility from patients undergoing routine tonsillectomy for non-malignant indications. The de-identified tonsil tissue was provided by the Core to the laboratory for *in vitro* B cell assays. The UAB Institutional Review Board (IRB) determined this study to be non-human subjects research (#N120216001).

#### Mice

All experimental animals were bred and maintained in the UAB animal facilities. All procedures involving animals were approved by the UAB Institutional Animal Care and Use Committee (IACUC, protocol-22088) and were conducted in accordance with the principles outlined by the National Research Council. Mouse strains used in these experiments were originally obtained from either Jackson Laboratory (B6.129S2-*Ighm^tm1Cgn^*/J (μMT), and B6.129S2-*Irf1*^tm1Mak^/J (*Irf1*^−/−^)) or from Dr. Sylvia Bolland at NIH (B6.SB-Yaa/J.B6;129S-*Fcgr2b^tm1Ttk^*/J (referred to as YaaFc mice)).^21^ New mouse strains were generated at UAB by intercrossing YaaFc mice with μMT mice to generate B cell deficient YaaFc.μMT mice^15^ and by intercrossing *Irf1*^−/−^ mice with YaaFc mice to generate *Irf1* deficient YaaFcc.*Irf1*^−/−^ mice. Experiments were performed on mice between 5-10 months of age. Male mice were utilized for all YaaFc experiments as the TLR7 duplication that drives autoimmune disease in these animals is Y chromosome linked.

### Method details

#### Human tonsillar naïve B cell purification

Human tonsil tissue was collected, dissected, mechanically dissociated, and filtered through a 70μm cell strainer. Mononuclear cells were isolated by density-gradient centrifugation over Lymphocyte Separation Medium (CellGro). Naïve B cells were purified by negative selection using the EasySep Naïve B Cell Enrichment Kit (STEMCELL Technologies), followed by positive selection with IgD magnetic microbeads (Miltenyi Biotec) according to the manufacturer’s instructions.

#### Naïve human B cell cultures

Purified naïve B cells were cultured in complete human B cell medium (Iscove’s modified Dulbecco’s medium supplemented with 10% heat-inactivated fetal bovne serum (FBS), 2mM L-glutamine, penicillin (200μg/mL), streptomycin (200μg/mL), and Normocin (InvivoGen)). When specified, cells were labeled with CellTrace Violet (CTV; Thermo Fisher) for 10 min at 37°C, quenched with complete medium, washed, and resuspended in complete media. Naïve B cells were plated at 1×10^6^ cells/mL in 96-well plates and stimulated with polyclonal anti-IgM F(ab’)_2_ (5μg/mL, Jackson ImmunoResearch) and recombinant human IL-2 (5ng/mL, PeproTech), in the presence or absence of R848 (0.1-10μg/mL, InvivoGen). When noted, recombinant human IFNγ (20ng/mL, PeproTech) was added to cultures. On day 3 (D3), cultures were washed, replated at 0.2×10^6^ cells/mL, and stimulated with IL-21 (50ng/mL, PeproTech) in the presence or absence of R848. In some experiments, human B cells were transduced on the indicated day with different lentiviral vectors (see below) in the presence of protamine sulfate (0.027ng/mL). For ASC differentiation experiments, cells were harvested on D6, counted, and analyzed by flow cytometry or ELISPOT.

#### Lentivirus constructs and virus production

The parent self-inactivating lentiviral vector pRRL-MND-eGFP^115^ (Addgene #36247, referred to as LVGFP) was modified to remove eGFP and the remaining plasmid backbone was used to generate vectors expressing human *IRF1* linked to reporter genes (GFP or mouse Thy1.1 (*Thy1a*)). Briefly, gene blocks were constructed containing (i) an internal ribosome entry site (IRES) and mouse *Thy1a* (GenBank accession AY445633.1); (ii) human *IRF1* (GenBank accession NM_002198.3) cloned upstream of the IRES-*Thy1a* cassette or (iii) *IRF1* cloned upstream of an IRES_GFP cassette. The gene blocks were cloned into the plasmid backbone to generate LV_IRES_*Thy1a* (referred to at LVThy1), LV_*IRF1*_IRES_GFP (referred to as LV*IRF1*_GFP_), and LV_*IRF1*_IRES_*Thy1a* (referred to as LV*IRF1* _Thy1_). All constructs were verified by Sanger sequencing.

Each construct described above was transiently transfected with packaging plasmid (psPAX2, Addgene #12260) and envelope plasmid (pMD2G, Addgene #12259) into 293T cells (ATCC, Manassas VA) using polyethylenimine (Polysciences) according to standard procedures. Transfected cells were maintained in DMEM plus L-glutamine, 10% FBS, and penicillin+streptomycin for 4 days. Supernatants were harvested, clarified and concentrated by low-speed centrifugation to enrich for viral particles, which were subsequently titrated as previously described.^116^

#### Human B cell flow cytometry

Single cell suspensions were blocked with 2% human AB serum or 10μg/mL human gamma globulin (Jackson ImmunoResearch, Cat#. 009-000-002) and stained with fluorochrome-conjugated antibodies (Table S6). 7-AAD or LIVE/DEADTM Fixable Dead Cell Stain Kits (ThermoFisher Scientific) were used to identify viable cells. For transcription factor staining, the Transcription Factor Staining Buffer Set (ThermoFisher Scientific) was used according to the manufacturer’s instructions. Stained cells were analyzed on a FACSCanto II (BD Biosciences). Cell sorting was performed on a FACSAria (BD Biosciences) in the UAB Comprehensive Flow Cytometry and Single Cell Core. Data were analyzed using FlowJo v10.

#### Human ELISPOT

Serial dilutions of D6 *in vitro* activated human B cells were transferred to anti-IgG-coated PVDF ELISPOT plates (Millipore) and incubated for 6 hr at 37°C. Plates were washed, incubated with alkaline-phosphatase-conjugated polyclonal anti-human IgG Ab (1:1000 dilution, Jackson ImmunoResearch) for 1-2 hr at 37°C, washed and incubated with alkaline phosphatase substrate (Moss, Inc.) according to manufacturer’s instructions. ELISPOTs were detected using a CTL ELISPOT reader.

#### Human *IRF1* RT-PCR

Total RNA was isolated from cultured human B cells using the Single Cell RNA Purification Kit (Norgen BiotekTM, Cat#.51800). RNA quantity and quality were assessed using an Nanodrop 6000 and an Agilent 2100 Bioanalyzer. cDNA was synthesized from total RNA using the SuperScriptTM Double-Stranded cDNA Synthesis Kit (InvitrogenTM, Cat#.11917010) with random hexamers, according to the manufacturer’s instructions. Real-time PCR was performed using TaqManTM Gene Expression Master Mix (Applied BiosystemsTM), under the following cycling conditions: 50°C for 2 min, 95 °C for 10 min, followed by 45 cycles of 95 °C for 15 s and 60 °C for 1 min. Pre-designed TaqManTM Gene Expression Assays (Applied BiosystemsTM) were used (*IRF1*: Hs00971965_m1; *GAPDH*: Hs02758991_g1). Gene expression levels were normalized to the housekeeping gene *GAPDH* and relative expression of *IRF1* was calculated using the 2− ddCT method, with the control sample set to 1.0.

#### Generation of mouse bone marrow chimeras

Bone marrow (BM) chimeric mice were generated using male B cell-deficient YaaFc.μMT recipients that were lethally irradiated with 850 rads from a high-energy X-ray source, administered as a split dose delivered ≥4 hours apart. Following irradiation, recipient YaaFc.μMT mice^15^ were reconstituted via intravenous (i.v.) injection with male donor BM cells (5×10^6^-1×10^7^ BM cells) that were mixed at an 80:20 ratio with 80% YaaFc.μMT BM + 20% YaaFc BM (B-YaaFc mice) or with 80% YaaFc.μMT BM + 20% YaaFc.*Irf1*^−/−^ BM (B-YaaFc.*Irf1*^−/−^ mice). B-YaaFc chimeras were competent to express *Irf1* in all cell types. In B-YaaFc.*Irf1*^−/−^ chimeras, the *Irf1* gene was deleted in 100% of B cells but was intact in 100% of all radiation-resistant cell types and in 80% of all radiation-sensitive hematopoietic cell types. Chimeras were analyzed between 5-10 months post-reconstitution.

#### Cell isolation from mouse tissues

Spleen and mediastinal lymph nodes (medLN) were collected into staining media and single-cell suspensions were prepared by disrupting tissues on fine wire mesh screens. BM cells were obtained by flushing femurs with PBS. Resultant single cell suspensions from all tissues were filtered (70μm nylon mesh filter), red blood cells were lysed (Hybri-MAX, Sigma-Aldrich), cells were washed in sterile PBS containing 2% FBS, filtered and counted to determine total viable cell numbers per-tissue.

#### Flow cytometry analysis of mouse cells

Cell suspensions were incubated with Fcγ receptor-blocking Ab (2.4G2; 10 μg/mL) diluted in FACS staining buffer (PBS, 2% FBS, 2 mM EDTA) for 15 min at 4°C. Cells were stained with fluorochrome-conjugated Abs specific for cell surface markers (See Table S6 for list of Abs used in flow cytometry analyses) and with LIVE/DEAD Fixable Aqua Dead (Invitrogen), for 15 min at 4°C then washed with staining buffer. In some analyses, stained cells were fixed and permeabilized using the eBioscience™ Fixation/Permeabilization buffer set (00-5523-00, Thermo Fisher Scientific), incubated for 1 hour (hr) at room temperature (RT) with Abs specific for intracellular proteins. Cells were washed and resuspended in FACS buffer for acquisition. Flow cytometric data were acquired on a FACSCanto II (BD Biosciences) or Attune NxT (Invitrogen, Thermo Fisher Scientific) and analyzed using FlowJo (v10, Tree Star). B cell subset gating strategies are shown in Fig. S7. The frequency of each B cell subset was determined and the absolute cell number per subset within a tissue was calculated by multiplying the frequency of each population by the total viable cell count for that tissue.

#### *In vitro* differentiation of mouse B cells

Splenic FOB (CD4^−^CD8^−^CD138^−^GL-7^−^CD19^+^CD21^−^CD23^+^), MBC (CD4^−^CD8^−^CD19^+^CD138^−^GL-7^−^CD38^+^IgD^−^) and ABC (CD4^−^CD8^−^CD19^+^CD138^−^GL-7^−^CD11b^+^CD11c^+^) subsets were sort-purified on a FACSAria (BD Biosciences) in the UAB Comprehensive Flow Cytometry and Single Cell Core. Purified cells were cultured in complete mouse B cell media (RPMI, 10% heat-inactivated FBS, 1% each of non-essential amino acids, sodium pyruvate, and HEPES buffer, 2% penicillin/streptomycin, 0.001% 2-Mercaptoethanol) in 96 well plates (50,000-100,000 cells/ml) in the presence of 10μg/mL anti-Kappa (Fab’2) (SouthernBiotech), 1μg/mL R848 (InvivoGen), 100U/mL IL-2 (PeproTech) 100ng/mL IL-21 (PeproTech) and 20 ng/ml IFNγ (PeproTech). For some experiments, the cells were stained for 10 min at 37°C with PBS diluted CellTrace Violet (Molecular Probes, Thermofisher) and washed with complete media prior to culture. On day 4 (D4), cells were harvested and analyzed by flow cytometry or ELISPOT to enumerate ASC.

#### Mouse ELISPOT

To detect Ig or IgG producing ASC, ELISPOT plates (Millipore, MAHAS4510) were pre-coated with anti-mouse IgG Ab (SouthernBiotech) overnight at 4°C. To detect dsDNA-specific ASC, ELISPOT plates were coated with methyl-BSA (10μg/ml, Sigma) for 2 hr at 37°C, washed with PBS/0.01% Tween 20, coated with calf thymus DNA in PBS (10μg/ml, Sigma) and incubated overnight at 4°C. Coated plates were washed and blocked with complete media. Cell suspensions in complete media were added (2000 cells/well) to the wells and incubated for 6 hr at 37°C. ELISPOT plates were washed and alkaline phosphatase-conjugated goat anti-mouse heavy chain-specific IgG Ab (Southern Biotech) was added (100μl/well) and incubated for 1-2 hr at 37°C. Plates were washed, developed with BCIP/NBT (Moss Substrates) and ELISPOTs were visualized using a CTL ELISPOT reader. The number of spots per well (after correction for non-specific background) were recorded using a CTL Immunospot S6 Macroplate Imager Reader (New Life Scientific) and counted manually. Data reported as the total number of ASC in the spleen generated by back-calculation total number of spots to the total number of splenic cells.

#### Mouse kidney histopathology

YaaFc, YaaFc.*Irf1*^⁻/⁻^, B-YaaFc, and B-YaaFc.*Irf1*^⁻/⁻^ kidneys were collected at the indicated timepoints, fixed in 10% neutral buffered formalin for ≥ 24 hr at RT at a tissue-to-fixative ratio of ≥1:10 and then processed and analyzed by the UAB Animal Resources Program Comparative Pathology Laboratory. Briefly, samples were paraffin-embedded, sectioned at 5 μm, mounted on glass microscope slides and stained with hematoxylin and eosin (H&E). Stained kidney sections were evaluated by a board-certified veterinary pathologist who was blinded to the genotypes of the mice, using a subjective scoring system adapted from the method of Austin and colleagues^117^ as modified by Kevil *et al*.^118^ The pathologist assessed glomerular cellularity, cell necrosis, neutrophil accumulation, glomerular capillary basement membrane changes, sclerosis, interstitial mononuclear inflammatory cell accumulation, tubular changes, and interstitial fibrosis. Each feature was scored on a 0-3 scale, corresponding to normal (0), mild (1), moderate (2), or severe (3) lesions. Scores for necrosis and crescent formation were weighted by a factor of 2 when calculating composite lesion scores. For each mouse, digital images of 6-15 glomeruli with adjacent tubules and interstitium were evaluated, with equal representation of glomeruli from superficial, mid, and deep cortex. The lesion score for each animal was calculated as the mean of the summed individual scores for each image, incorporating the weighted values for necrosis and crescent formation. Brightfield images showing multiple non-overlapping fields encompassing both glomerular and tubulointerstitial regions were acquired on a Nikon Eclipse Ci microscope using a 20× objective. Images were analyzed using NIS-Elements software (Nikon), and all samples from a given experiment were imaged under identical illumination and acquisition settings to permit qualitative comparison of renal pathology across genotypes. Representative images were exported into Adobe Illustrator for figure assembly.

#### Mouse kidney immune complex deposition

YaaFc and YaaFc.*Irf1*^⁻/⁻^ kidneys were harvested, rinsed in ice-cold PBS, embedded in OCT (Tissue-Tek), snap frozen on dry ice, and stored at -80°C until sectioned. Serial 7μm sections were cut on a cryostat, deposited onto Superfrost Plus microscope slides (Thermo Fisher Scientific), fixed in acetone (-20°C) for 10 min, and stored under dessication at 4°C. Tissue sections were incubated for 20-30 min at RT in blocking buffer (10% FBS (v/v in PBS)) containing 10μg/mL Fcγ receptor blocking Ab, 2.4G2. Slides were stained for 45-60 min at RT with 50μL FITC-conjugated anti-mouse IgG antibody (BioLegend, 1:100). Slides were washed in PBS, rinsed in distilled water, mounted with ProLong™ Diamond Antifade Mountant (Thermo Fisher Scientific) and coverslipped. Images were acquired at RT on a Nikon Eclipse Ti inverted fluorescence microscope using the NIS-Elements AR v4.20.02 software. FITC signal was collected using a FITC filter set. Within each experiment, all samples were imaged with identical exposure times and acquisition settings. Scores were assigned using a semiquantitative scoring system adapted from Sang Won Lee *et al*.^119^ IgG deposition intensity was graded on a 0 to 3 scale, where 0 indicates no/negligible deposition and 3 indicates intense deposition. At least 10 fields (∼50 glomeruli) were examined per mouse. Representative images were exported for figure preparation in Adobe Illustrator. Post acquisition processing was limited to figure assembly.

#### Mouse ANA detection

Serum anti-nuclear Ab (ANA) was detected using Kallestad HEp-2 cell slides (Bio-Rad). Briefly, mouse sera was serially diluted (starting at 1:40) in PBS and incubated in individual HEp-2 slide wells for 1 hr at RT. Slides were washed, incubated with 25μL FITC-conjugated goat anti-mouse IgG (Invitrogen, diluted 1:50-1:1600 in PBS + 1% BSA) for 1hr at RT in the dark, washed, mounted with ProLong™ Diamond Antifade Mountant with DAPI (Thermo Fisher Scientific) and coverslipped. Images were acquired on a Nikon Eclipse Ti inverted fluorescence microscope equipped with FITC/Alexa Fluor 488 filter sets using 20× or 40× objectives. Within each experiment, all samples were imaged using identical exposure settings and representative fields were captured for each serum sample. ANA seropositivity was defined by the presence of a clear nuclear/peri-nuclear/cytoplasmic fluorescence pattern in HEp-2 cells at one or more serum dilutions compared with PBS-only negative control wells. Representative images were exported to Adobe Illustrator for layout and figure preparation. Only uniform linear adjustments to contrast were applied across the full field, followed by panel assembly. Identical processing parameters were applied to all images within an experiment, and no region-specific adjustments were performed.

#### Single cell multiome

FACS-sorted splenic IgD^neg^ B cells (See Table S6 for sort panel Ab) were washed with PBS with 0.04% BSA followed by cell lysis with 200μL of chilled Lysis Buffer (10mM Tris-HCl, 10mM NaCl, 3mM MgCl2, 0.10% Tween-20, 0.10% IGEPAL, 0.01% digitonin, 1% BSA, 1mM DTT, 1U/µL RNase inhibitor (Protector RNase inhibitor, Sigma)), for 5 min on ice. Chilled Wash Buffer (10mM Tris-HCl, 10mM NaCl, 3mM MgCl2, 0.10% BSA, 0.10% Tween-20, 1mM DTT, 1U/µL RNase inhibitor (Protector RNase inhibitor, Sigma)) was added (200µL) to lysed cells and mixed five times by pipetting. After washing with chilled Wash Buffer. Nuclei were stained with TotalSeq™-A-anti-Nuclear Pore Complex Proteins Hashtag Ab in chilled Wash Buffer for 30 min on ice (See Table S6 for hashtag Abs), then washed, pooled, resuspended in 1X Nuclei Buffer (10x Genomics PN 200020) and processed with Single-cell Multiome Kit (10x Genomics PN-1000230). Gene expression, ATAC and feature barcoding (totalseq A antibody) libraries were prepped following manufacturers’ instructions. All libraries were sequenced using the Illumina NovaSeq6000 in the UAB Heflin Genomics Core.

Gene expression libraries were sequenced using 28 cycles for Read 1 and 90 cycles for Read 2, with 10 cycles each for the i7 and i5 index reads. ATAC libraries were sequenced using 50 cycles for Read 1 and 49 cycles for Read 2, with 8 cycles for the i7 index read and 24 cycles for the i5 index read. Sequencing was performed to target depths of 20,000 reads per nuclei for the gene expression library, 25,000 reads per nuclei for the ATAC library, and 10,000 reads per nuclei for the feature barcoding library.

#### Single-cell multiome data analysis

Raw base-call (BCL) files generated by the 10x Genomics Chromium platform were converted to FASTQ format using the Cell Ranger mkfastq function (v7.1.0). For single-cell multiome (gene expression + chromatin accessibility) libraries, paired nuclei gene expression and ATAC FASTQ files were processed with the Cell Ranger ARC pipeline (v2.0.2) using the 10x Genomics mouse reference genome mm10-2020-A-2.0.0. The count function was used to perform read alignment, filtering of low-quality reads, barcode and UMI counting, peak calling, and generation of gene expression and peak-by-barcode matrices. Hashtag oligonucleotide (HTO) libraries were processed separately using the Cell Ranger Count pipeline (v9.0.1), with both gene expression and feature barcode FASTQ files used as input to generate HTO count matrices.

Downstream analysis was performed in R using Seurat^79^ (v5.2.1) and Signac^120^ (v1.14.0) and in python (v3.11.13) using Scanpy^121^ (v1.11.4) and MIRA^71^ (v2.1.1). A Seurat object containing the RNA, ATAC, and HTO assays was created and low-quality nuclei were removed based on standard quality-control metrics: only nuclei with ≥1,000 unique ATAC fragments per nuclei and ≥200 detected genes in the RNA assay with <25% of the RNA reads mapping to mitochondrial genes were retained for further analysis. HTO-based sample demultiplexing was carried out with Seurat’s HTODemux function using default parameters and cells classified as doublets or “negative” for all HTOs were excluded. Additional doublets were identified and removed using DoubletFinder (v2.0.4).

For the RNA modality, counts were normalized and scaled using standard Seurat workflows, and highly variable genes were identified for downstream dimensionality reduction. For the ATAC modality, peak counts were processed with Signac, including term frequency-inverse document frequency (TF-IDF) normalization and latent semantic indexing (LSI) to obtain a reduced set of chromatin-accessibility dimensions. A joint low-dimensional representation was computed using Seurat’s weighted nearest neighbor (WNN) analysis, which integrates RNA PCA and ATAC LSI embeddings to construct a shared neighbor graph. Clustering was performed on the WNN graph using the Louvain algorithm implemented in Seurat’s FindClusters function at empirically chosen resolutions, and uniform manifold approximation and projection (UMAP) was used to generate two-dimensional embeddings for visualization of the integrated RNA-ATAC landscape. Clusters were annotated as different B cell subsets based on the expression of a curated list of canonical markers of major B cell subsets.

#### ROGUE scoring

Transcriptional homogeneity within clusters was quantified using the ROGUE^55^ (Ratio of Global Unshifted Entropy) package (v1.0) with the RNA assay set as the default expression slot in the Seurat object. Raw UMI count matrices were extracted and converted to a gene-by-cell expression matrix. Data was filtered with *matr.filter* to retain genes detected in at least 10 cells and cells expressing at least 10 genes and global expression entropy across genes was computed using the *SE_fun* function. Overall dataset-level ROGUE scores were derived using *CalculateRogue* with platform = “UMI”. To assess homogeneity at the cluster level, the rogue function was applied to the filtered expression matrix, using the Seurat clustering identities (resolution 0.8; wsnn_res.0.8) as the cluster labels and the sample of origin (group) as the samples factor, with platform = “UMI” and a smoothing span of 0.6. This yielded per-sample, per-cluster ROGUE scores that were subsequently summarized by computing the mean ROGUE value for each cluster (excluding NA values). Data was visualized by plotting the relationship between mean expression and entropy using *SEplot*, and the distribution of ROGUE scores across samples for each cluster using boxplots, with clusters ordered according to their average ROGUE score.

#### Single cell differential gene expression analysis

Differentially expressed genes (DEGs) analysis for the RNA assay was performed using the presto R package^122^ using the on log-normalized expression values. Presto was used to compute Wilcoxon rank sum statistics and associated p-values using a Gaussian approximation, together with the area under the receiver operating characteristic curve (auROC) as an effect size metric. P-values were adjusted for multiple testing using the Benjamini Hochberg procedure. Genes or peaks with |avg_log2FC| ≥0.1, FDR < 0.05 and expressed in ≥10% of cells per cluster were considered significant.

#### Differential chromatin accessibility

Differential chromatin accessibility was assessed using Seurat/Signac’s implementation of *FindMarkers* on the ATAC assay, employing the logistic regression framework (test.use = “LR”). Peaks with an |avg_log2FC| ≥0.2, FDR < 0.05 and expressed in ≥10% of cells per cluster/group were defined as significantly differentially accessible.

#### Single nuclei ATAC peak calling with MACS2

Chromatin accessibility peaks were identified using MACS2^123^ peak caller within the *CallPeaks* function in the Signac R package. ATAC fragments across all retained nuclei were aggregated by Signac to generate a pseudo-bulk ATAC profile. Enriched regions of accessibility were identified by MACS2 with default parameters.

#### Gene activity scores

To infer gene-level chromatin accessibility from ATAC-seq signal, we computed gene activity scores^120^ using the Signac implementation of the *GeneActivity* function applied to the ATAC assay. A gene-by-cell “gene activity” count matrix, which was generated by mapping ATAC fragments to promoter-proximal and annotated regulatory regions for each gene, was imported into the Seurat object as a separate assay. Gene activity counts were normalized using Seurat LogNormalize method, with gene activity counts in each cell scaled by a factor equal to the median RNA UMI count (nCount_RNA) across all cells and then log-transformed.

#### Module Scoring for ASC commitment scores

To assess ASC commitment module scores, we used the *AddModuleScore* function in Seurat^79^ on a curated gene set (Table S3) that was assembled from a published dataset.^80^ Module scores were computed on log-normalized RNA expression values using Seurat default parameters.

#### Transcription factor motif enrichment and motif activity analysis

TF motif enrichment and motif activity were analyzed using Signac chromVAR package.^120^ CORE vertebrate motif position frequency matrices were obtained from JASPAR2020^124^ via TFBSTools and added to ATAC peaks with *AddMotifs*. Peak level sequence features computed using *RegionStats*. Motif enrichment on DARs was assessed using *FindMotifs*, with multiple testing correction by Benjamini-Hochberg and significance defined as FDR < 0.05. Motif activity at single cell resolution was quantified using chromVAR via the *RunChromVAR* function.

#### Transcription factor motif footprinting

TF Footprints were computed using the ATAC assay and the BSgenome.Mmusculus.UCSC.mm10 reference genome. The Signac *Footprint* function was used to aggregate Tn5 insertion events across genomic instances of each motif and to generate an insertion profile centered on the motif with flanking sequence on both sides. To reduce technical confounding from transposase sequence preferences, observed insertion profiles were normalized by an expected background model that accounts for Tn5 sequence bias, yielding a bias corrected footprint signal for each motif. Footprint signals were stored in the ATAC assay for downstream analysis and visualization. Motif footprints were visualized using the *PlotFootprint* function.

#### Peak-gene linkage analysis

Peak to gene linkages were inferred using Signac. Peak level sequence features were computed using the *RegionStats* function using the mm10 genome, and peak-gene associations were identified with the *LinkPeaks* function. Peak-gene links were visualized using the *CoveragePlot* function.

#### Single cell Gene Set Enrichment Analysis (GSEA)

GSEA was performed using gene sets obtained from the Molecular Signatures Database (MSigDB) curated collections (Hallmark and C7 immunological gene sets) and custom gene sets derived from published datasets. These gene sets were used to query pre-ranked single nuclei DEG lists. For each comparison, DEGs were ranked by a signed enrichment score defined as the product of the average log_2_ fold change and the -log_10_ of the adjusted p value. Pre-ranked GSEA was carried out using the GSEA implementation in the clusterProfiler R package (v4.10.1),^125^ with default parameters. For each analysis, the enrichment score (ES) and normalized enrichment score (NES) were computed based on permutations of the gene labels, and multiple-testing correction was performed using the Benjamini-Hochberg method to control the false discovery rate. Pathways with an adjusted p value < 0.05 were considered significantly enriched. All published gene sets used in these analyses that are not found in the Molecular Signatures Database^126,127^ are found in Table S3.

#### Multimodal models for integrated regulatory analysis

Single-cell gene expression and chromatin accessibility profiles were modeled using MIRA^71^ (v2.1.1) to obtain a shared low-dimensional representation of cellular state. For the RNA modality, a topic model on highly variable genes was constructed using raw counts and exogenous/endogenous dispersion covariates to separate technical from biological variability. The number of expression topics was selected by combining gradient-based topic contribution diagnostics with Bayesian hyperparameter tuning (min 5, max 50 topics), and the final model was used to derive per-cell topic loadings and corresponding UMAP features. An analogous topic model was fit to the ATAC modality using highly variable peaks as features, a dedicated accessibility encoder, and a similar gradient/Bayesian tuning procedure (centered around 15 topics). For both modalities, MIRA “UMAP feature” representations were used as inputs to Scanpy (v1.11.4) to construct k-nearest-neighbor graphs (k = 20) and UMAP embeddings (min_dist = 0.1). A joint multimodal embedding was obtained by aligning the expression- and accessibility-derived topic spaces with mira.utils.make_joint_representation, yielding a shared “X_joint_umap_features” representation used for downstream visualization and clustering.

To link TF binding to gene regulation, MIRA Regulatory Potential (RP) framework was applied.^71^ Candidate RP genes were defined as the union of all highly variable genes in the RNA topic model and the top topic-associated genes (top 200 per topic), restricted to genes with annotated transcription start sites (TSS) in the mm10 reference. For these genes, LITE regulatory potential models were trained to predict gene expression from local chromatin accessibility around the TSS using matched RNA and ATAC AnnData objects. Probabilistic *in silico* deletion (pISD) was performed by integrating these RP models with CistromeDB-processed ChIP–seq peak sets.^70^ For each TF-gene pair, pISD quantified the change in the regulatory potential predicted for the gene when all chromatin peaks overlapping the TF ChIP-seq peaks were computationally “deleted” (masked) from the RP model to simulate a TF-specific perturbation.

#### Bulk RNA-seq library preparation and analysis

D3 *in vitro* activated human B cells were collected and placed in RL buffer (Norgen Biotek) supplemented with 1% β-mercaptoethanol, snap-frozen on dry ice, and sent to Emory Genomics Core for library preparation and sequencing. Total RNA was isolated using the Quick-RNA Microprep kit (Zymo Research) and used as input for the SMART-seq v4 cDNA synthesis kit (Takara) with 12 cycles of PCR. cDNA was quantitated and 200 pg of material was used with the NexteraXT kit and NexteraXT Indexing primers (Illumina, Inc) in 12 cycles of PCR to generate libraries. Samples were quality checked on a bioanalyzer, quantitated by Qubit fluorometer, and pooled at equimolar ratios for sequencing on an Illumina S4.

Raw sequencing image files (.bcl) generated on the Illumina platform were converted to FASTQ format and demultiplexed by sample using Illumina’s bcl2fastq software (v2.17), with default settings for base calling and index assignment. Residual adapter sequences and low-quality bases were removed from the resulting FASTQ files using Skewer (v0.2.2)^128^ and only reads passing Skewer’s internal quality filters were retained for downstream analysis. Trimmed reads were aligned to the human reference genome (hg38) using STAR (v2.7.8a)^129^ with the UCSC hg38 KnownGene annotation as the reference transcriptome. Alignment was performed with default parameters and coordinate-sorted BAM files were generated for each sample. PCR duplicate reads were identified and flagged using Picard MarkDuplicates (v3.3.0). Gene-level counts were computed in R (v4.4.0) using the GenomicRanges package (v1.54.1)^130^ by intersecting aligned reads with annotated gene models from the KnownGene table and summarizing overlaps to obtain raw counts per gene for each sample. Reads Per Kilobase of transcript per Million mapped reads (RPKM) values were calculated from these counts. DEG analysis was performed using DESeq2 (v1.42.1).^131^ Raw gene counts were imported into DESeq2, and a generalized linear model based on the negative binomial distribution was fit with the experimental group as the primary design factor. Size factors and dispersion parameters were estimated using DESeq2 standard workflow, and Wald tests were used to evaluate differential expression between conditions. Resulting p values were adjusted for multiple testing using the Benjamini-Hochberg procedure to control the false discovery rate (FDR). Genes with an absolute log_2_ fold change >1 and FDR <0.05 were considered significantly differentially expressed.

#### Analysis of published datasets

Single cell RNA sequencing data from Rosain *et al*.^108^ (GSE216489) was retrieved from the NCBI GEO database and the Perez *et al*. dataset^110^ was downloaded from cellxgene. Both data were imported into R, Seurat object was created, B cells were subsetted and processed using the standard Seurat workflow: normalization, identification of highly variable genes, scaling and linear and non-linear dimensionality reduction. For the Lino *et al*. (Regulatory ASC) gene set,^63^ downloaded affymetrix CEL files for IL-10eGFP^pos^CD138^hi^ and IL-10eGFP^neg^CD138^hi^ cells (GSE103458)^63^ were background corrected, quantile normalized and summarized to expression values using RMA.^132^ Differential expression for the contrast IL-10eGFP^pos^CD138^hi^ over IL-10eGFP^neg^CD138^hi^ was computed with limma,^133^ and probes were mapped to gene symbols using the Bioconductor mouse4302.db annotation package.^134^ To generate gene sets, significantly upregulated genes were selected using an FDR <0.05 and log_2_ fold change (log_2_FC) > 1 cutoffs. These gene lists were exported in GMT format. For the Shi et al. ASC vs GCB (GSE60927),^80^ Tellier et al. WT vs *Blimp1*KO-WT (GSE70981) and WT vs *Xbp1*KO-WT (GSE70981)^88^ gene sets, raw count tables were imported into R and DEG analysis was performed using DESeq2.^131^ Log_2_FC shrinkage was applied using apeglm^135^ to stabilize effect size estimates. Gene sets were defined from DEGs using an FDR <0.05 and |log_2_FC| ≥ 3 for GSE60927 and |log2FC| ≥ 1 for GSE70981. The gene sets were exported in GMT format. In both cases, the GMT files were used as input for GSEA. Bulk RNA-seq and ATAC-seq data (GSE118256)^109^ were analyzed as previously described.^109^ Genes were considered detected if expression > 3 rpkm in least one experimental group. DEGs were identified using edgeR v3.18.1^136^ and a gene was considered significant based on FDR <0.05 and |log2FC| ≥ 2. For the ATAC-seq data, motifs within accessible chromatin peaks were annotated using HOMER v4.8.2 (annotatePeaks.pl, option “-size given”). Motif footprints were generated by computing read depth across motif instances and flanking sequences using GenomicRanges v1.22.4 and custom R and Bioconductor scripts (https://github.com/cdschar), with per-base coverage visualized as histograms or summarized as box plots.

### Statistical analyses

Statistical analyses were performed using GraphPad Prism v.10.1.2 (GraphPad Software). Comparison of two groups was performed using unpaired two-tailed Student’s t tests. Comparison of three or more groups was performed using One-way ANOVA. Correlations were tested using Pearson’s correlation. Analyses of bulk and single-cell datasets were conducted in R (v4.4.0 and v4.3.0) and python (v3.11.13), with additional details provided in the relevant Methods subsections. DEGs were identified using auROC and Wilcoxon p-value based on Gaussian approximation (presto). Genes with |avg_log2FC| ≥0.1, FDR <0.05 and expressed in ≥10% of cells per cluster or group were considered significant. Peaks with an |avg_log2FC| ≥0.2, FDR < 0.05 and expressed in ≥10% of cells per cluster/group were considered DARs. GSEA was performed on a pre-ranked gene list (bulk RNA-seq) or pre-ranked DEG list (single-cell RNA-seq). Normalized enrichment scores (NES) and adjusted p value for multi-comparisons were determined using the GSEA package.

