## supplement figures and legends for "IFNγ-induced IRF1 synergizes with TLR7 signals to tune the IRF4-IRF8 axis and drive pathogenic effector B cell fate"

#### Figure S1

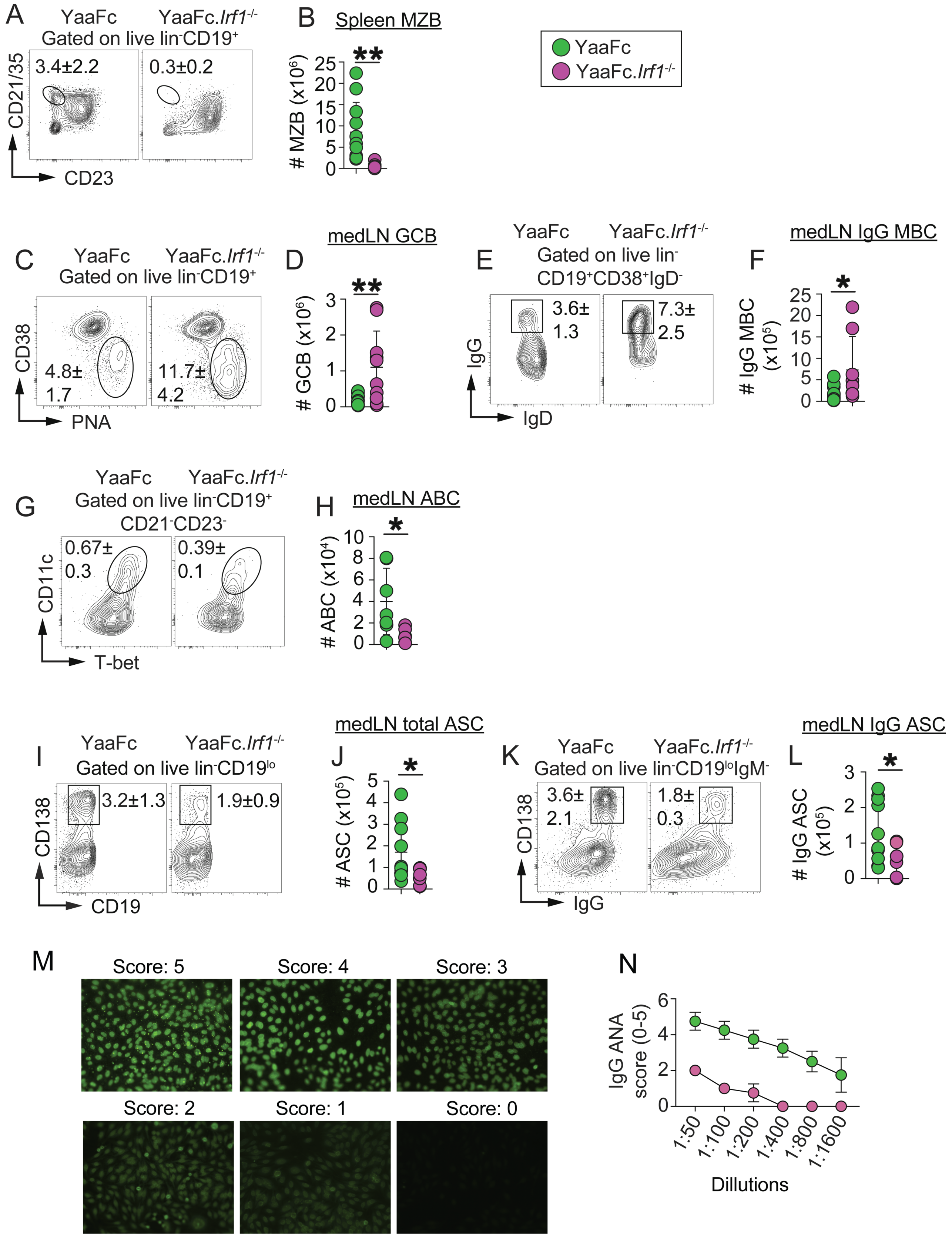
A
YaaFc
YaaFc.Irf1-/-
Gated on live lin-CD19+
3.4±2.2
0.3±0.2
CD21/35
CD23
B
Spleen MZB
### MZB (x10^6)
**
YaaFc
YaaFc.Irf1-/-
C
Gated on live lin-CD19+
4.8± 1.7
11.7± 4.2
CD38
PNA
D
medLN GCB
### GCB (x10^6)
E
Gated on live lin- CD19+CD38+IgD-
3.6± 1.3
7.3± 2.5
IgG
IgD
F
medLN IgG MBC
### IgG MBC (x10^5)
G
Gated on live lin-CD19+ CD21-CD23-
0.67± 0.3
0.39± 0.1
CD11c
T-bet
H
medLN ABC
### ABC (x10^4)
I
Gated on live lin-CD19lo
3.2±1.3
1.9±0.9
CD138
CD19
J
medLN total ASC
### ASC (x10^5)
K
Gated on live lin-CD19loIgM-
3.6± 2.1
1.8± 0.3
IgG
L
medLN IgG ASC
### IgG ASC (x10^5)
M
Score: 5
Score: 4
Score: 3
Score: 2
Score: 1
Score: 0
N
IgG ANA score (0-5)
1:50
1:100
1:200
1:400
1:800
1:1600
Dillutions

**Figure S1. *Irf1* regulates homeostasis of the B cell compartment and autoreactive ASC responses in lupus-prone Yaa.*Fcgr2b*$^{-/-}$ mice, related to Figure 1 and Figure 2.**

(**A-B**) Representative flow cytometry plots (**A**) showing splenic MZB cells from 5-month YaaFc and YaaFc.*Irf1*$^{-/-}$ mice. Data reported as frequency (**A**) and absolute number (**B**) of cells.

(**C-L**) Flow cytometric quantification of B cell subsets in medLN from 5-month YaaFc and YaaFc.*Irf1*$^{-/-}$ mice. Representative flow plots shown and data reported as frequencies and absolute number of GCB cells (**C-D**), IgG$^{+}$ MBC (**E-F**), CD11c$^{+}$T-bet$^{+}$ ABC (**G-H**), CD138$^{+}$ ASC (**I-J**) and IgG$^{+}$ CD138$^{+}$ ASC (**K-L**).

(**M-N**) Binding of serum IgG anti-nuclear Abs (ANA) to HEp-2 cells with representative ANA images showing the scoring system used (**M**) and serially diluted (1:50-1:1600 dilution) binding assay data reported as ANA scores for each dilution (**N**) of serum from 5-month YaaFc and YaaFc.*Irf1*$^{-/-}$ mice.

Data representative of two independent experiments (n ≥ 5 mice/group) and reported for individual mice in each group. Significance determined by unpaired two-tailed Student's t tests. *$p < 0.05$, **$p < 0.01$.

### Figure S2

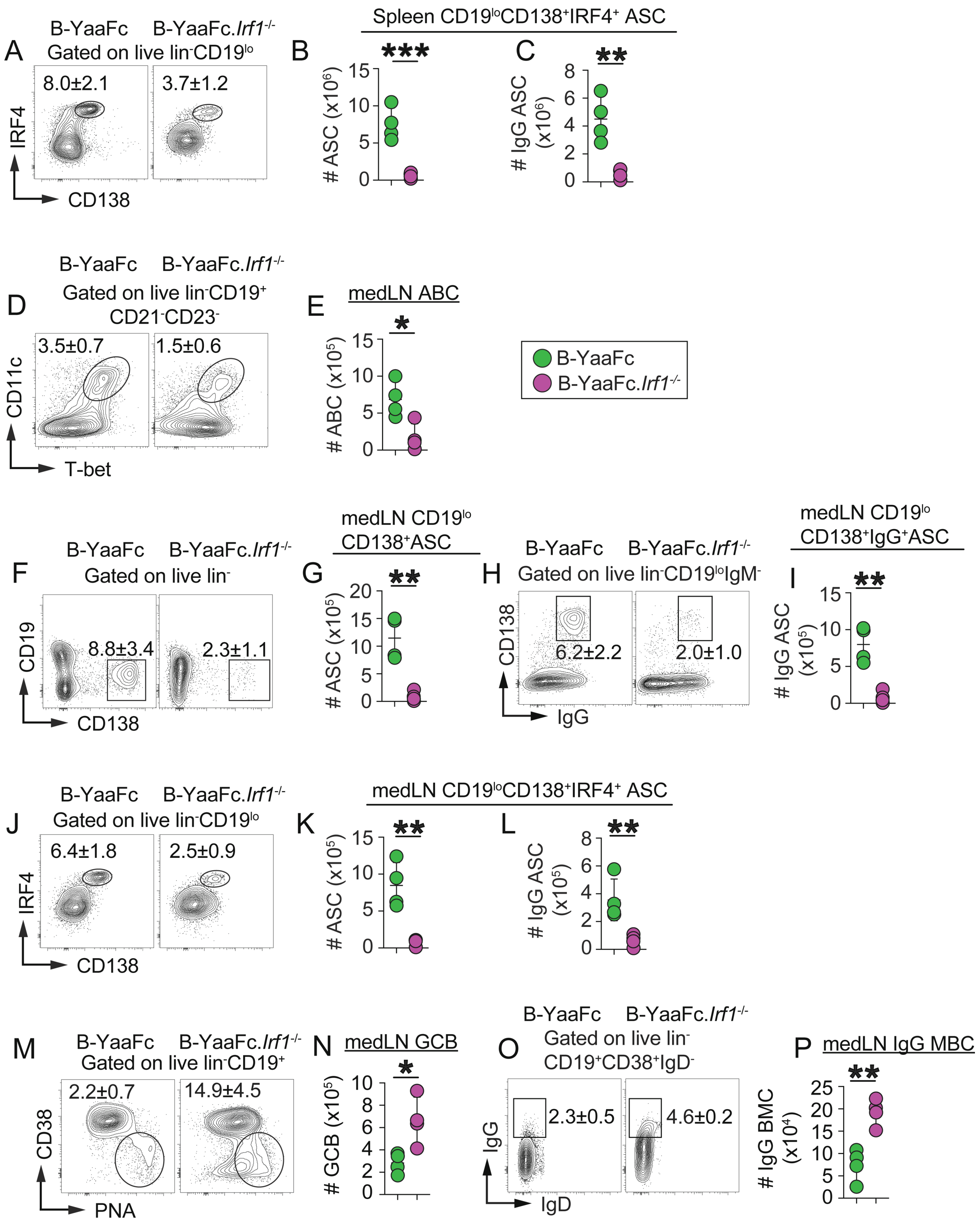

A
B-YaaFc
B-YaaFc.Irf1-/-
Gated on live lin-CD19lo
8.0±2.1
3.7±1.2
IRF4
CD138
Spleen CD19loCD138+IRF4+ ASC
B
***
### ASC (x10^6)
C
**
### IgG ASC (x10^6)
D
Gated on live lin-CD19+ CD21-CD23-
3.5±0.7
1.5±0.6
CD11c
T-bet
E
medLN ABC
*
### ABC (x10^5)
B-YaaFc
B-YaaFc.Irf1-/-
F
Gated on live lin-
8.8±3.4
2.3±1.1
CD19
CD138
G
medLN CD19lo CD138+ASC
**
### ASC (x10^5)
H
Gated on live lin-CD19loIgM-
6.2±2.2
2.0±1.0
CD138
IgG
I
medLN CD19lo CD138+IgG+ASC
**
### IgG ASC (x10^5)
J
Gated on live lin-CD19lo
6.4±1.8
2.5±0.9
IRF4
CD138
medLN CD19loCD138+IRF4+ ASC
K
**
### ASC (x10^5)
L
**
### IgG ASC (x10^5)
M
Gated on live lin-CD19+
2.2±0.7
14.9±4.5
CD38
PNA
N
medLN GCB
*
### GCB (x10^5)
O
Gated on live lin- CD19+CD38+IgD-
2.3±0.5
4.6±0.2
IgG
IgD
P
medLN IgG MBC
**
### IgG BMC (x10^4)

**Figure S2. B cell-intrinsic *Irf1* is required for expansion of pathogenic B cell subsets and autoreactive ASC in lupus-prone mice, related to Figure 3.**

(**A-C**) Representative flow cytometry plots (**A**) showing splenic ASC from 10-month B-YaaFc and B-YaaFc.*Irf1*$^{-/-}$ chimeras. Data reported as frequency (**A**) and absolute number of CD19$^{lo}$CD138$^{+}$IRF4$^{+}$ ASC (**B**) and absolute number of IgG$^{+}$ CD19$^{lo}$CD138$^{+}$IRF4$^{+}$ ASC (**C**).

(**D-P**) Flow cytometric quantification of B cell subsets in the medLN from 10-month B-YaaFc and B-YaaFc.*Irf1*$^{-/-}$ chimeras. Representative flow plots shown and data reported as frequency and absolute numbers of ABC (**D-E**), CD19$^{lo}$CD138$^{+}$ASC (**F-G**), IgG$^{+}$ CD19$^{lo}$CD138$^{+}$ASC (**H-I**), CD19$^{lo}$CD138$^{+}$IRF4$^{+}$ ASC (**J-K**), IgG$^{+}$ CD19$^{lo}$CD138$^{+}$IRF4$^{+}$ ASC (**L**), GCB cells (**M-N**) and IgG$^{+}$ MBC (**O-P**).

Data representative of two independent experiments ($n \geq 4$ mice/group) and reported for individual mice in each group. Significance determined by unpaired two-tailed Student's t tests. *$p < 0.05$, **$p < 0.01$, ***$p < 0.001$.

### Figure S3

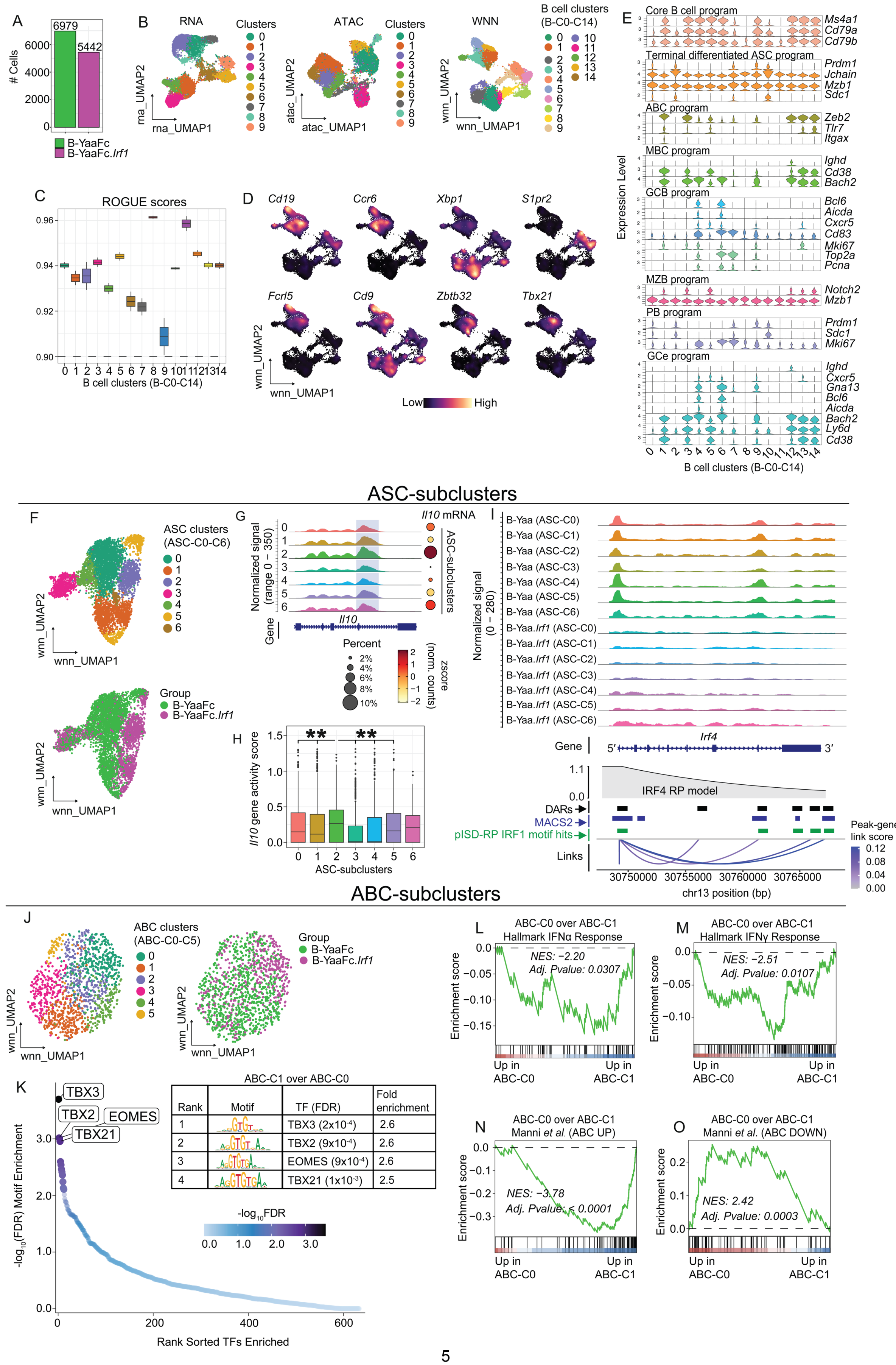


ABC-C1 over ABC-C0

| Rank | Motif | TF (FDR) | Fold enrichment |
|---|---|---|---|
| 1 | | TBX3 (2x10⁻⁴) | 2.6 |
| 2 | | TBX2 (9x10⁻⁴) | 2.6 |
| 3 | | EOMES (9x10⁻⁴) | 2.6 |
| 4 | | TBX21 (1x10⁻³) | 2.5 |

**Figure S3. IRF1 regulates the transcriptome and epigenome of splenic ASC and ABC from lupus-prone mice, related to Figure 4.**

(**A-E**) Paired snRNA-seq and snATAC-seq multiomics profiling of IgD$^{neg}$ splenic B cells from 6-month post-reconstitution B-YaaFc and B-YaaFc.*Irf1*$^{-/-}$ chimeras. Shown are the number of cells/nuclei of each genotype that passed all quality control steps (**A**) and were used in downstream bioinformatic analyses. UMAP visualization of the B cell clusters (**B**) defined based on RNA modality only (left), ATAC modality only (middle) and WNN integrated multimodal analysis (right), which resulted in 15 integrated B cell clusters (B-C0 to B-C14) that were highly homogeneous based on ROGUE scores (**C**). Expression of canonical marker genes (**D-E**) was used to annotate major B cell subsets. B-C0, B-C2, B-C8, B-C10 and B-C11 clusters were annotated as non-proliferating terminally-differentiated ASC based on lower *Cd19* and higher *Prdm1*, *Jchain*, *Xbp1*, *Mzb1*, and *Sdc1*. Cluster B-C1 was annotated as ABC-like cells with higher *Zeb2*, *Tlr7*, *Itgax*, and *Tbx21*. Clusters B-C3, B-C13 and B-C14 were annotated as MBC based on lower *Ighd* and higher *Cd38*, *Bach2*, *Ccr6*, and *Zbtb32*. Clusters B-C4 and B-C6 were annotated as GCB cells based on expression of *Bcl6*, *Aicda*, *S1pr2*, and *Cxcr5*. B-C4 was annotated as light zone (LZ) GCB cells based on high *Cd83* expression, while B-C6 was annotated as dark zone (DZ) GCB cells that expressed *Mki67*, *Top2a*, and *Pcna*. Cluster B-C5 was annotated as MZB cells based on higher *Cd19*, *Cd9*, *Notch2*, *Fcrl5* and *Mzb1*. Cluster B-C7 was annotated as proliferating ASC (plasmablast-PB) based on high expression of *Prdm1* and *Sdc1* along with residual *Cd19* and high *Mki67*. The B-C9 cluster was annotated as GC-emigrant GC-exit cells (GCe) based on lack of *Ighd*, retention of *Cxcr5* and *Gna13*, downregulation of *Bcl6* and *Aicda*, and higher expression of *Bach2*, *Ly6d*, and *Cd38*. Cluster B-C12 was annotated as naïve B cells that express *Ighd*.

(**F-I**) Cells assigned to the non-proliferating ASC clusters (B-C0, B-C2, B-C8, B-C10 and B-C11) were combined, unsupervised sub-clustering was performed, and 7 ASC subclusters (ASC-C0 to ASC-C6) were identified. UMAP (**F**) showing ASC-subclusters grouped by cluster ID (top) or genotype (bottom). (**G-H**) *Il10* gene analysis in ASC subclusters. Chromatin accessibility surrounding the *Il10* gene (**G**), expression of *Il10* (**G,** right) and *Il10* gene activity score (**H**) in the individual ASC-subclusters shown. *Irf4* gene analysis (**I**), showing chromatin accessibility within each ASC-subcluster, split by ASC genotype showing DAR (I, black bars), MACS2-called peaks (**I**, blue bars), predicted IRF1 binding sites in the *Irf4* locus using multimodal integrated regulatory potential modeling[70] following probabilistic *in silico* deletion of TF binding regions (pISD-RP)[71] (**I**, green bars) and peak to gene links.

(**J-O**) Unsupervised subclustering was performed on the B cells assigned to the ABC cluster (B-C1). UMAP (**J**) showing the 6 ABC-subclusters (ABC-C0 to ABC-C5) grouped by ABC-subcluster ID (left) or by genotype (right). TF motif enrichment analysis (**K**) comparing cells in clusters ABC-C1 over ABC-C0. GSEA (**L-O**) using the Hallmark Type I IFN (**L**), the Hallmark Type II IFN (**M**) response gene set, and a curated[37] geneset (GSE99480) of genes upregulated (**N**) or downregulated (**O**) in ABC to query the ranked DEG list of ABC-C0 over ABC-C1.

Significance (**H**) was assessed using Kruskal-Wallis test, followed by pairwise Wilcoxon rank sum tests with Benjamini-Hochberg (BH) adjustment for multiple comparisons. Significant over-represented TF binding motifs (**K**) defined as motifs with adjusted p-value <0.05 and fold enrichment > 1.2. Adjusted p value and NES scores for each GSEA indicated (see also **Table S3**). See **Table S2** for DEGs of total B clusters, number of cells per cluster among total B cells, and average expression of genes used for annotating B cell clusters. **p < 0.01.

### Figure S4

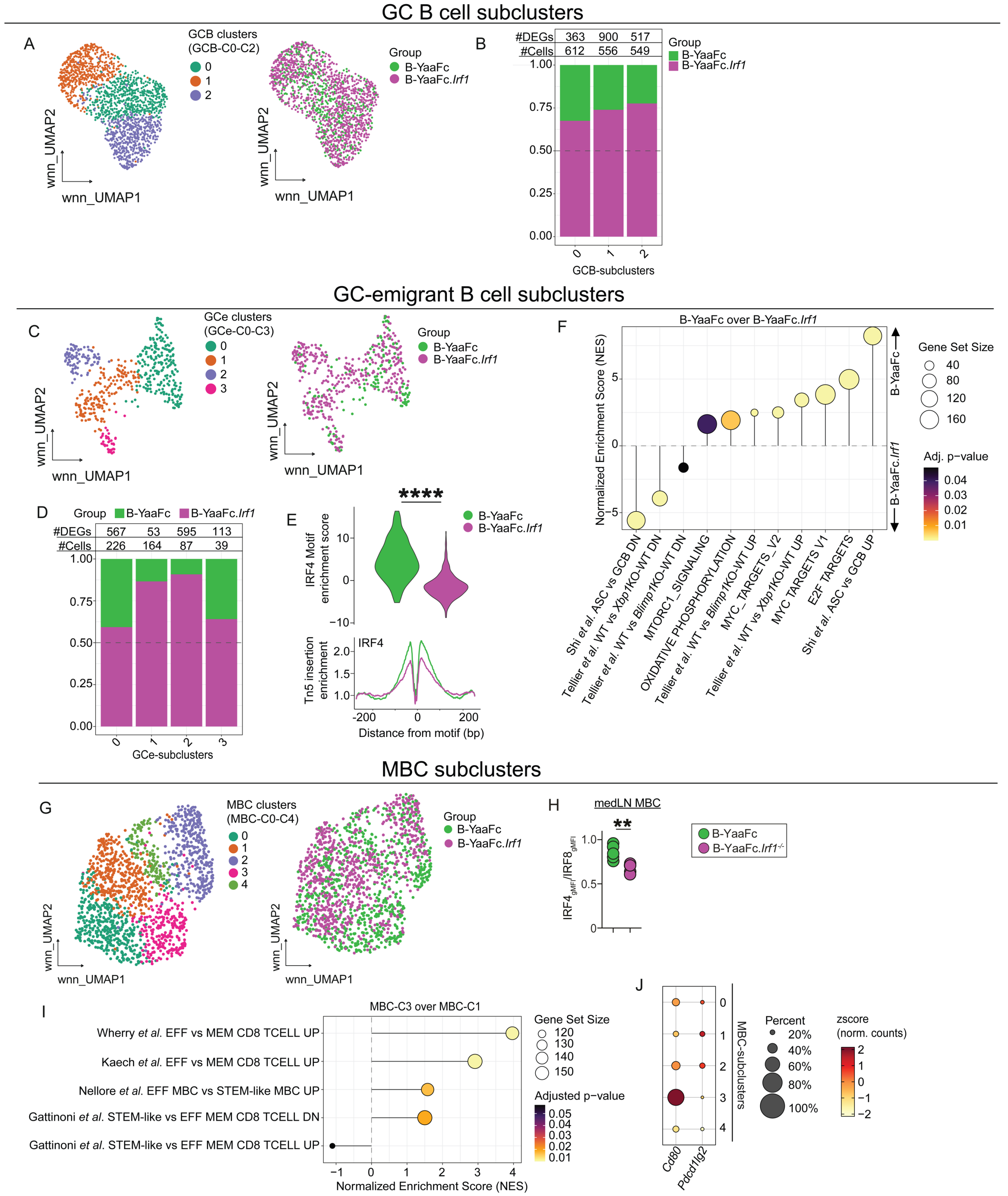

GC B cell subclusters
A
GCB clusters
(GCB-C0-C2)
0
1
2
wnn_UMAP2
wnn_UMAP1
Group
B-YaaFc
B-YaaFc.Irf1
B
#DEGs 363 900 517
#Cells 612 556 549
1.00
0.75
0.50
0.25
0.00
0 1 2
GCB-subclusters
GC-emigrant B cell subclusters
C
GCe clusters
(GCe-C0-C3)
0
1
2
3
D
#DEGs 567 53 595 113
#Cells 226 164 87 39
GCe-subclusters
E
****
IRF4 Motif enrichment score
10
0
−10
IRF4
Tn5 insertion enrichment
2.0
1.5
1.0
-200 0 200
Distance from motif (bp)
F
B-YaaFc over B-YaaFc.Irf1
Normalized Enrichment Score (NES)
Gene Set Size
40
80
120
160
Adj. p-value
0.04
0.03
0.02
0.01
Shi et al. ASC vs GCB DN
Tellier et al. WT vs Xbp1KO-WT DN
Tellier et al. WT vs Blimp1KO-WT DN
MTORC1_SIGNALING
OXIDATIVE PHOSPHORYLATION
Tellier et al. WT vs Blimp1KO-WT UP
MYC_TARGETS_V2
Tellier et al. WT vs Xbp1KO-WT UP
MYC TARGETS V1
E2F TARGETS
Shi et al. ASC vs GCB UP
MBC subclusters
G
MBC clusters
(MBC-C0-C4)
0
1
2
3
4
H
medLN MBC
**
IRF4gMFI/IRF8gMFI
1.0
0.5
0
B-YaaFc.Irf1-/-
I
MBC-C3 over MBC-C1
Wherry et al. EFF vs MEM CD8 TCELL UP
Kaech et al. EFF vs MEM CD8 TCELL UP
Nellore et al. EFF MBC vs STEM-like MBC UP
Gattinoni et al. STEM-like vs EFF MEM CD8 TCELL DN
Gattinoni et al. STEM-like vs EFF MEM CD8 TCELL UP
−1 0 1 2 3 4
Normalized Enrichment Score (NES)
Gene Set Size
120
130
140
150
Adjusted p−value
0.05
0.04
0.03
0.02
0.01
J
MBC-subclusters
Cd80
Pdcd1lg2
Percent
20%
40%
60%
80%
100%
zscore
(norm. counts)
2
1
0
−1
−2

**Figure S4. IRF1 regulates the *Irf4*:*Irf8* balance and supports effector over stem-like transcriptional programs in post-GC B cells in lupus-prone mice, related to Figure 5.**

(**A-B**) Cells assigned to the GCB cell clusters (B-C4 and B-C6) were combined and unsupervised sub-clustering was performed to identify 3 GCB subclusters (GCB-C0 to GCB-C2). UMAP (**A**) showing GCB-subclusters grouped by cluster ID (left) or genotype (right). Data in (**B**) reported as the number of cells and DEG assigned to each GCB cell subcluster and the relative abundance of the B-YaaFc and B-YaaFc.*Irf1*$^{-/-}$ B cells within the different GCB cell subclusters.

(**C-F**) Unsupervised subclustering was performed on the B cells assigned to the GC-emigrant (GCe) cluster (B-C9). UMAP (**C**) showing 4 GCe subclusters (GCe-C0 to GCe-C3) grouped by cluster ID (left) or genotype (right). Data in (**D**) reported as the number of cells and DEG assigned to each GCe subcluster and the relative abundance of the B-YaaFc and B-YaaFc.*Irf1*$^{-/-}$ B cells within the different GCe subclusters. Chromatin accessibility surrounding IRF4-binding motifs in GCe cells (**E**) showing IRF4 motif enrichment scores and the IRF4 binding footprint between B-YaaFc and B-YaaFc.*Irf1*$^{-/-}$ GCe cells. GSEA (**F**) using gene sets comparing ASC to GCB,[80] WT vs *Blimp1*-KO B cells,[88] WT vs *Xbp1*-KO B cells,[88] and Hallmark gene sets relevant to ASC differentiation/function to query the ranked DEG list of B-YaaFc GCe cells over B-YaaFc.*Irf1*$^{-/-}$ GCe cells.

(**G-J**) Multiomics (**G, I-J**) and flow cytometric (**H**) analyses of MBC from B-YaaFc and B-YaaFc.*Irf1*$^{-/-}$ mice.

(**G, I-J**) Cells assigned to the MBC clusters (B-C3, B-C13 and B-C14) were combined and unsupervised sub-clustering was performed to identify 5 MBC subclusters (MBC-C0 to MBC-C4). UMAP (**G**) showing subclusters grouped by cluster ID (left) or by genotype (right). GSEA (**I**) using gene sets comparing effector vs memory CD8 T cells,[101,102] stem-like vs effector memory CD8 T cells,[103] and human FCRL5$^+$ effector vs stem-like MBC,[100] to query a ranked DEG list of MBC-C3 over MBC-C1 cells. Expression of *Cd80* and *Pdcd1lg2* (**J**) by the MBC-subclusters.

(**H**) Flow cytometric analysis showing IRF4 and IRF8 protein expression by CD38$^+$IgD$^{neg}$ MBC from medLN of 6-month post-reconstitution B-YaaFc and B-YaaFc.*Irf1*$^{-/-}$ chimeras. The ratio of IRF4 to IRF8 protein expression is shown.

DEGs (**B**, **D**) based on auROC and Wilcoxon p-value based on Gaussian approximation. Significance in (**E**, **H**) was assessed using Wilcoxon Rank Sum (**E**) or unpaired two-tailed Student's t (**H**) tests. Adjusted p value and NES scores for each GSEA indicated (see also **Table S3**). **p < 0.01, ****p < 0.0001.

### Figure S5

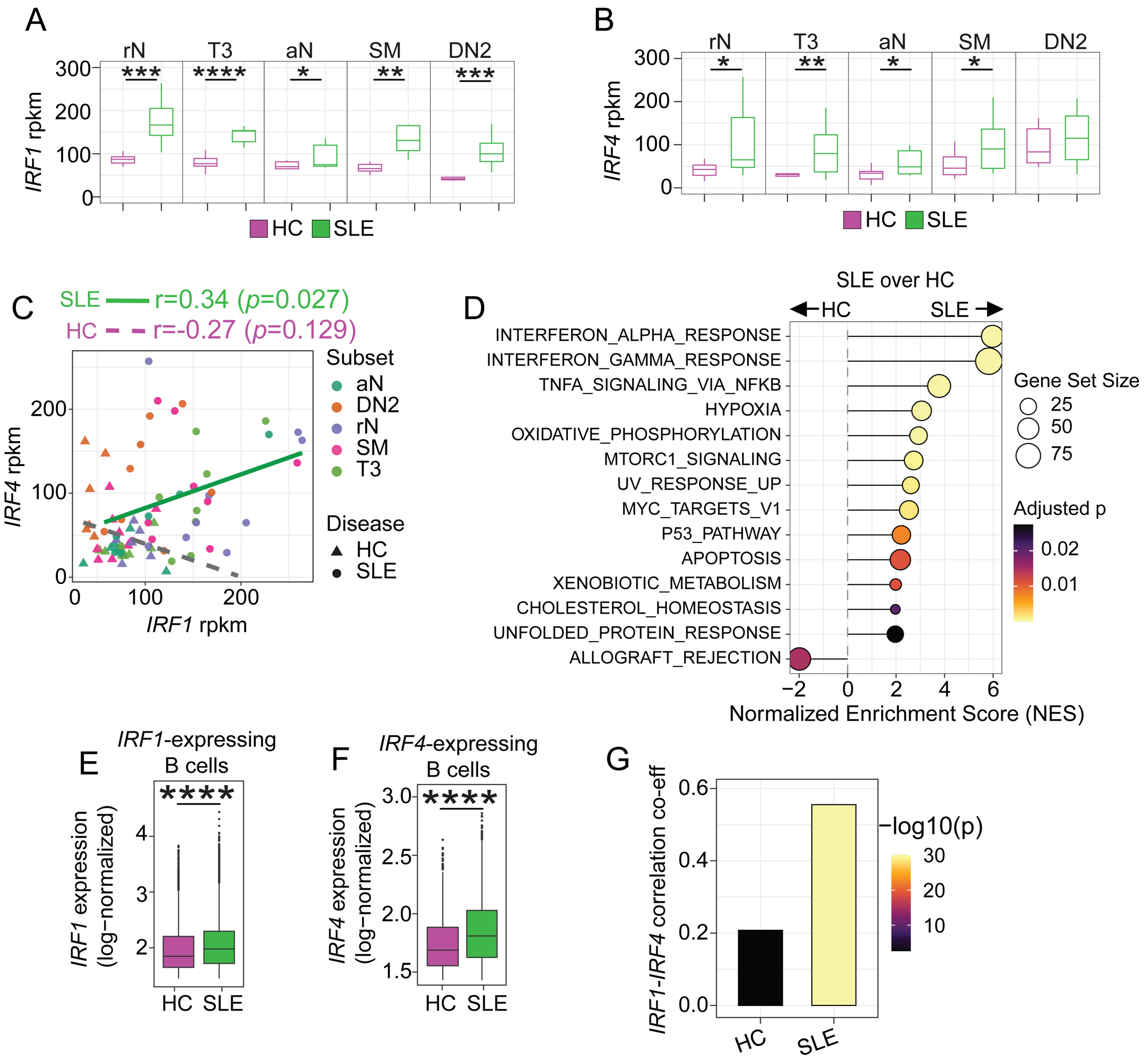
A
rN
T3
aN
SM
DN2
***
****
*
**
***
IRF1 rpkm
300
200
100
0
HC SLE
B
rN
T3
aN
SM
DN2
*
**
*
*
IRF4 rpkm
300
200
100
0
HC SLE
C
SLE r=0.34 (p=0.027)
HC r=-0.27 (p=0.129)
IRF4 rpkm
IRF1 rpkm
0
100
200
Subset
aN
DN2
rN
SM
T3
Disease
HC
SLE
D
SLE over HC
HC
SLE
INTERFERON_ALPHA_RESPONSE
INTERFERON_GAMMA_RESPONSE
TNFA_SIGNALING_VIA_NFKB
HYPOXIA
OXIDATIVE_PHOSPHORYLATION
MTORC1_SIGNALING
UV_RESPONSE_UP
MYC_TARGETS_V1
P53_PATHWAY
APOPTOSIS
XENOBIOTIC_METABOLISM
CHOLESTEROL_HOMEOSTASIS
UNFOLDED_PROTEIN_RESPONSE
ALLOGRAFT_REJECTION
−2 0 2 4 6
Normalized Enrichment Score (NES)
Gene Set Size
25
50
75
Adjusted p
0.02
0.01
E
IRF1-expressing B cells
****
IRF1 expression (log−normalized)
4
3
2
HC SLE
F
IRF4-expressing B cells
****
IRF4 expression (log−normalized)
3.0
2.5
2.0
1.5
HC SLE
G
IRF1-IRF4 correlation co-eff
0.6
0.4
0.2
0.0
−log10(p)
30
20
10
HC
SLE

**Figure S5. *IRF1* expression is increased in SLE B cells and its expression correlates with *IRF4* expression, related to Figure 7.**

(**A-C**) Analysis of published bulk RNAseq[109] datasets from purified B cell populations isolated from healthy controls (HC) and SLE patient blood samples. Box plots showing expression levels (reported as rpkm) of *IRF1* (**A**) and *IRF4* (**B**) in $CD19^{+}IgD^{+}CD27^{-}MTG^{-}CD24^{+}CD38^{+}$ resting naïve, $CD19^{+}IgD^{+}CD27^{-}$Mitotracker $Green^{+}CD24^{mid/+}CD38^{-}$ transitional T3, $CD19^{+}IgD^{+}CD27^{-}$Mitotracker $Green^{+}CD24^{-}CD38^{-}$ activated naive, $CD19^{+}IgD^{-}CD27^{+}$ isotype-switched memory, and $CD19^{+}IgD^{-}CD27^{-}CXCR5^{-}$ DN2 B cells. Center lines indicate the median, lower and upper bounds of boxes indicate the 1st and 3rd quartiles and whiskers indicate the upper and lower limits of the data. Correlation analysis (**C**) comparing *IRF1* and *IRF4* mRNA expression levels in different B cell populations isolated from HC and SLE patient blood samples.

(**D-G**) Analysis of published[110] single cell RNAseq datasets from HC and SLE B cells isolated from blood. GSEA (**D**) using Hallmark gene sets relevant to ASC differentiation and function to query a ranked DEG list of SLE B cells over HC B cells. *IRF1* and *IRF4* expression levels in HC and SLE B cells (**E-F**) showing *IRF1* expression levels within the *IRF1*$^{+}$ HC and SLE B cells (**E**) and *IRF4* expression levels (**F**) within the *IRF4*$^{+}$ HC and SLE B cells. Correlation analysis (**G**) between *IRF1* and *IRF4* expression levels within single *IRF1*$^{+}$ cells from HC and SLE donors. *IRF1*$^{+}$ and *IRF4*$^{+}$ B cells were defined as cells expressing at least one transcript of the gene.

Significance determined by unpaired two-tailed Student's t tests (**A-B**, **E-F**) or Pearson's correlation analysis (**D**). Adjusted p value and NES scores for each GSEA indicated (see also **Table S3**). *$p \leq 0.05$, **$p \leq 0.01$, ***$p \leq 0.001$, ****$p \leq 0.0001$.

### Figure S6

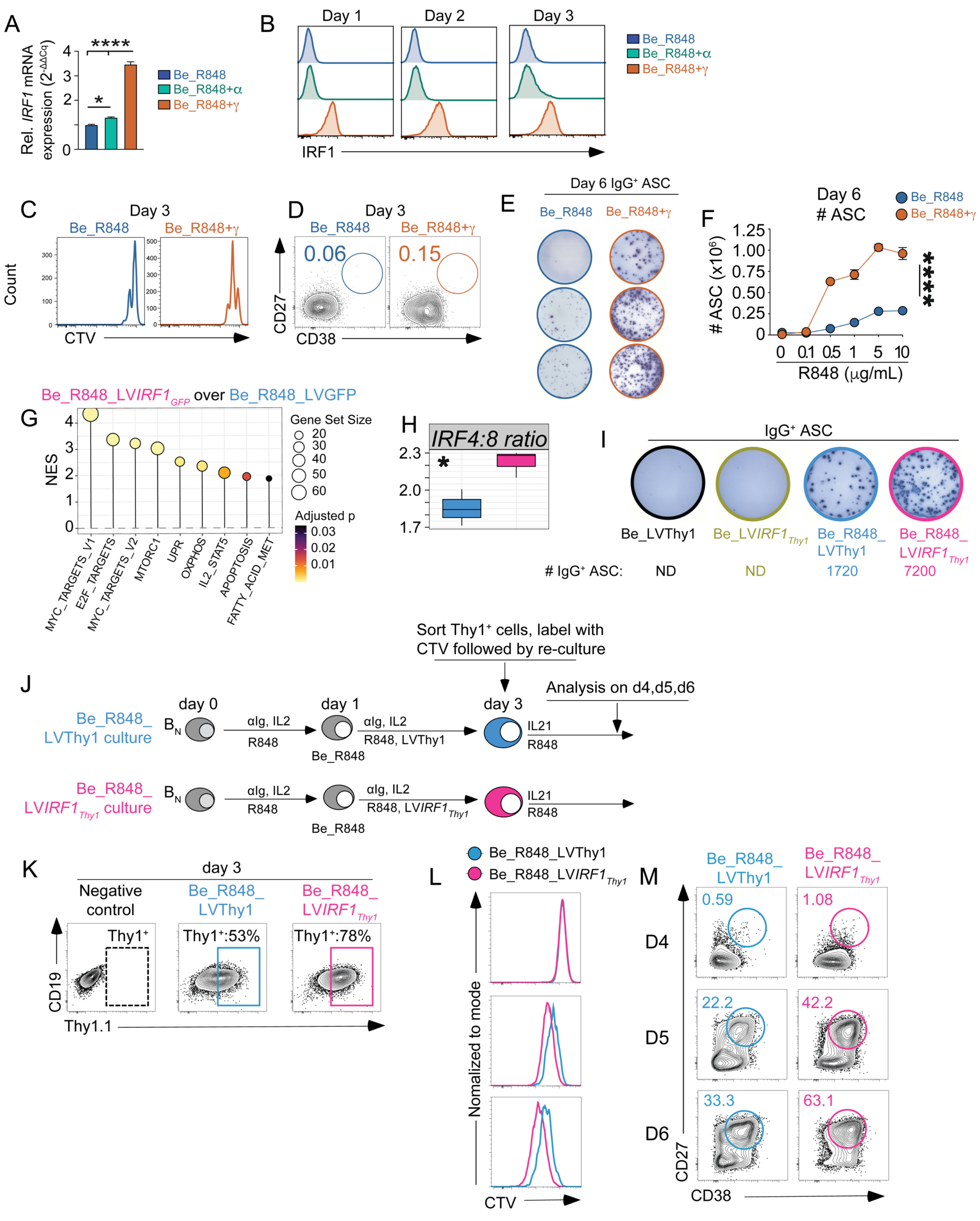

A
Rel. IRF1 mRNA expression (2^-ΔΔCq)
****
*
Be_R848
Be_R848+α
Be_R848+γ
B
Day 1
Day 2
Day 3
IRF1
C
Day 3
Be_R848
Be_R848+γ
Count
CTV
D
Day 3
0.06
0.15
CD27
CD38
E
Day 6 IgG⁺ ASC
F
Day 6
### ASC
### ASC (x10⁶)
R848 (μg/mL)
****
G
Be_R848_LVIRF1GFP over Be_R848_LVGFP
NES
Gene Set Size
Adjusted p
MYC_TARGETS_V1
E2F_TARGETS
MYC_TARGETS_V2
MTORC1
UPR
OXPHOS
IL2_STAT5
APOPTOSIS
FATTY_ACID_MET
H
IRF4:8 ratio
I
IgG⁺ ASC
Be_LVThy1
Be_LVIRF1Thy1
Be_R848_LVThy1
Be_R848_LVIRF1Thy1
### IgG⁺ ASC:
ND
ND
1720
7200
J
Sort Thy1⁺ cells, label with CTV followed by re-culture
Analysis on d4,d5,d6
Be_R848_LVThy1 culture
Be_R848_LVIRF1Thy1 culture
day 0
day 1
day 3
BN
αIg, IL2
R848
R848, LVThy1
R848, LVIRF1Thy1
Be_R848
IL21
K
day 3
Negative control
Thy1⁺
Thy1⁺:53%
Thy1⁺:78%
CD19
Thy1.1
L
Be_R848_LVThy1
Be_R848_LVIRF1Thy1
Nomalized to mode
CTV
M
D4
D5
D6
0.59
1.08
22.2
42.2
33.3
63.1
CD27
CD38

**Figure S6. *IRF1* cooperates with TLR7 signaling to induce IRF4 upregulation and promote human ASC differentiation, related to Figure 7.**

(**A-B**) Analysis of Be_R848, Be_R848+γ and Be_R848+α cultures. Cultures containing naïve B cells were set up as described in **Fig. 7E**. A similar culture in which IFNγ was replaced with IFNα was also included. *IRF1* mRNA (**A**) in D3 cells, quantitated by qPCR and normalized to *GAPDH.* IRF1 protein levels analyzed by flow cytometry (**B**) in D3 cells.

(**C-E**) Analysis of Be_R848 and Be_R848+γ cultures. Cultures containing CTV-labeled naïve B cells were set up as described in **Fig. 7E** and were analyzed on D3 (**C-D**) or D6 (**E**). Proliferation (**C**), as measured by CTV dilution, and $CD27^{+}CD38^{+}$ ASC (**D**), as enumerated by flow cytometry, are shown. ELISPOT analysis (**E**) of D6 $IgG^{+}$ ASC from each culture.

(**F**) Analysis of Be_R848 and Be_R848+γ cultures (see **Fig. 7E**) that were stimulated with increasing concentrations of R848 (0-10 μg/ml). Numbers of $CD27^{+}CD38^{+}$ ASC, as measured by flow cytometry, in D6 cultures, are shown.

(**G-H**) GSEA (**G**) using Hallmark gene sets related to ASC differentiation/function were used to query the ranked gene list of D3 Be_R848_LV*IRF1*$_{GFP}$ over Be_R848_LVGFP cultures. (**H**) *IRF4* to *IRF8* expression in D3 Be_R848_LVGFP and Be_R848_LV*IRF1*$_{GFP}$ cells reported as ratio.

(**I**) ELISPOT of D6 cultures from **Fig. 7P**.

(**J-M**) Analysis of IRF1- and TLR7-dependent *in vitro* human B cell proliferation and differentiation cultures. Experimental design schematic (**J**). Primary tonsil-derived HC naive B cells were activated with anti-IgM+IL2+R848 (Be_R848) and transduced on D1 with LV encoding mouse Thy1.1 (Be_R848_LVThy1, (blue)) or with LV encoding IRF1 and Thy1.1 (Be_R848_LV*IRF1*$_{Thy1}$ (pink) cultures). On D3, transduced $Thy1.1^{+}$ cells from both cultures were sorted, labeled with CTV, and re-cultured at equal numbers in the presence of IL-21+R848. Cells were analyzed for proliferation and ASC differentiation on D4, D5, and D6. Representative flow cytometry plots (**K**) showing the percentages of $Thy1.1^{+}$ cells at D3. Negative controls for Thy1.1 expression were non-transduced Be_R848. Flow cytometry plots showing CTV dilution profiles (**L**) and frequencies of $CD27^{+}CD38^{+}$ ASC (**M**) on D4, D5, and D6 in the cultures.

Significance (**F**, **H**) was determined by comparing AUC values between Be_R848 and Be_R848+γ cultures using unpaired two-tailed Student's t test (**F**) or unpaired two-tailed Student's t tests (**H**). Adjusted p value and NES scores for each GSEA indicated (see also **Table S3**). *$p \leq 0.05$, ****$p \leq 0.0001$.

### Figure S7

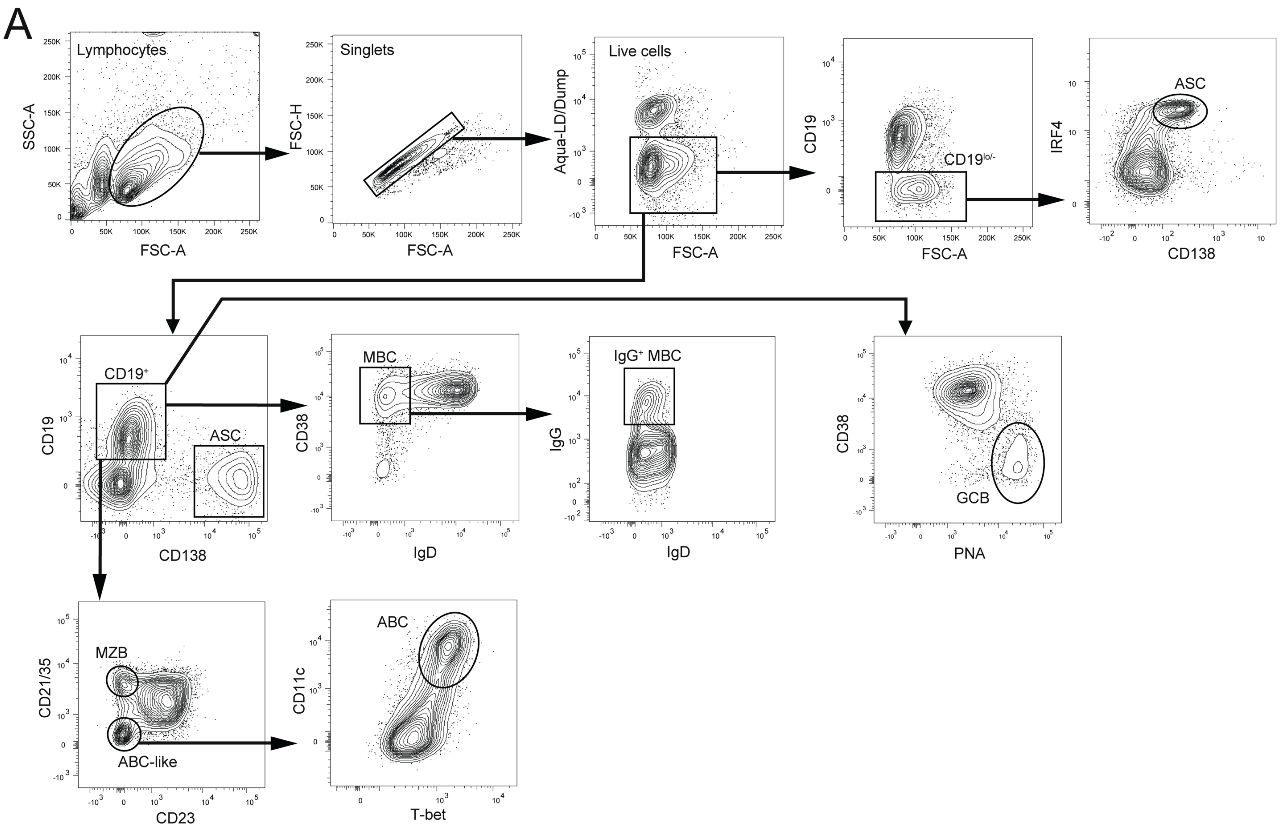
A
Lymphocytes
SSC-A
FSC-A
Singlets
FSC-H
FSC-A
Live cells
Aqua-LD/Dump
FSC-A
CD19
CD19lo/-
FSC-A
ASC
IRF4
CD138
CD19+
CD19
ASC
CD138
MBC
CD38
IgD
IgG+ MBC
IgG
IgD
CD38
GCB
PNA
MZB
CD21/35
ABC-like
CD23
ABC
CD11c
T-bet

**Figure S7. Gating strategy for flow cytometric analysis of mouse B cell subsets, related to Figures 1, 3, S1-2.** ASC are live$^{+}$CD19$^{lo/neg}$CD138$^{hi}$ or live$^{+}$CD19$^{lo/neg}$CD138$^{hi}$IRF4$^{+}$. MBC are live$^{+}$CD19$^{+}$IgD$^{neg}$CD38$^{+}$ or live$^{+}$CD19$^{+}$IgD$^{neg}$CD38$^{+}$IgG$^{+}$. GCB cells are live$^{+}$CD19$^{+}$CD38$^{lo}$PNA$^{hi}$. MZB cells are live$^{+}$CD19$^{+}$CD21$^{hi}$CD23$^{neg}$. ABC are live$^{+}$CD19$^{+}$CD21$^{neg}$CD23$^{neg}$CD11c$^{+}$Tbet$^{+}$.

**Supplemental Table Legends**

**Table S1**: Serologic autoreactivity and kidney histopathology in lupus model mice, related to **Figure 2** and **Figure 3**.

IgG ANA titers across serial serum dilutions in YaaFc and YaaFc.*Irf1*$^{-/-}$ mice, renal IgG deposition scores in YaaFc and YaaFc.*Irf1*$^{-/-}$ mice, renal histopathology scores from H&E stained kidney sections in YaaFc and YaaFc.*Irf1*$^{-/-}$ mice, and B-YaaFc and B-YaaFc.*Irf1*$^{-/-}$ mice. Significance was determined by unpaired two-tailed Student's t tests.

**Table S2**: Cluster composition, marker gene expression, and differential expression analyses across B cell, ASC, and ABC clusters, related to **Figure S3** and **Figure 4**.

DEGs across total B cell clusters, number of cells per cluster among total B cell clusters, average expression of marker genes used for annotation of B cell clusters, DEGs across ASC subclusters, DEGs comparing ASC-C0 and -C4 versus ASC-C2, DEGs across ABC subclusters, average expression of select marker genes across ABC subclusters and TF binding motif enrichment analysis of ABC-C1 over ABC-C0. DEGs identified using auROC and Wilcoxon p-value based on Gaussian approximation. Genes with |avg_log2FC| ≥0.1, FDR <0.05 and expressed in ≥10% of cells per cluster or group were considered significant.

**Table S3**: Gene set enrichment analyses across ASC, GC, MBC, and human SLE B cell comparisons, related to **Figure 4**, **Figure S4**, **Figure 5**, **Figure S5** and **Figure S6**.

Results of GSEA of ASC-C2 over ASC-C4 and -C0 using the Lino *et al*. regulatory ASC gene set,[63] ASC-C4 and -C0 over ASC-C2 using the Hallmark type I and type II IFN gene set, ABC-C0 over ABC-C1 using the Hallmark type I and type II IFN and public ABC[37] gene sets, B-YaaFc GCB over B-YaaFc.*Irf1*$^{-/-}$ GCB using the Good *et al*. plasma cell versus memory B cell upregulated gene set,[75] B-YaaFc GCe over B-YaaFc.*Irf1*$^{-/-}$ GCe using Hallmark and public gene sets,[75,85,88] MBC-C3 over MBC-C1 using Hallmark and public dataset-derived gene sets,[84,92,100-103] MBC-C1 over MBC-C3 using public dataset-derived gene sets,[100,104] SLE B cells over healthy control B cells from the Perez *et al*. 2022 dataset[110] using Hallmark gene sets, Be_R848_LV*IRF1*$_{GFP}$ over Be_R848_LVGFP using Hallmark gene sets, and the curated public dataset gene sets used for GSEA that are not found in the Molecular Signatures Database.[126,127]

**Table S4**: Integrated multiome clustering, differential expression, and motif enrichment across GC and MBC states, related to **Figure S4** and **Figure 5**.

Number of clusters and cells per cluster defined by WNN integration of RNA and ATAC profiles in GCB cells from B-YaaFc and B-YaaFc.*Irf1*$^{-/-}$ mice, DEGs of B-YaaFc GCB cells vs B-YaaFc.*Irf1*$^{-/-}$ GCB cells, DEGs of B-YaaFc GCe vs B-YaaFc.*Irf1*$^{-/-}$ GCe, average expression of select genes in GCe cells by genotype, DEGs across MBC subclusters, average expression of select genes across MBC subclusters, results of TF binding motif enrichment analysis of MBC-C3 over MBC-C1 and MBC-C1 over MBC-C3. DEGs were identified using auROC and Wilcoxon p-value based on Gaussian approximation. Genes with |avg_log2FC| ≥0.1, FDR <0.05 and expressed in ≥10% of cells per cluster or group were considered significant.

**Table S5**: Bulk transcriptomic expression and differential gene expression analyses in human B cell perturbation, related to **Figure 7**.

RPKM expression values and DEGs analysis for Be_R848_LVGFP versus Be_R848_LV*IRF1*$_{GFP}$ cells.

**Table S6**: Antibody reagents for flow cytometry, multiome cell sorting and nuclei hashing, related to **Figure S3-4** and **Figure 4-5**.

Antibodies used for flow cytometry, FACS sorting for multiome experiments, and hashtag antibodies used for nuclei labeling for multiome experiments.
